# Long-read analysis of complete human 5S rDNA arrays reveals a zoned organization with two-state unit methylation linked to chromatin architecture

**DOI:** 10.64898/2026.11.08.765174

**Authors:** Hyungmin Kim

## Abstract

Despite the fundamental importance of the 5S ribosomal DNA (rDNA) to all cellular life, the landscape of its units by position along its array has remained poorly understood. Here, recent telomere-to-telomere long-read sequencing is used to resolve 5S rDNA arrays from flank to flank and to follow the state of each unit along single molecules. The array is zoned at both the sequence and the epigenetic level. Notably, methylation is bimodal at the level of the unit, and the combination of low-methylation and high-methylation units gives rise to the overall methylation pattern of the array. Moreover, the average number of low-methylation units remains relatively constant regardless of array length. Chromatin fiber sequencing (Fiber-seq) data further show that low-methylation units are more accessible, have lower nucleosome occupancy and are more frequently footprinted at the TFIIIB-binding site, consistent with analyses of ATAC-seq and ChIP-seq data from cell lines. Across species, this pattern is shared among apes, whereas other species show diverse patterns. These results provide a unit-resolved view of the human 5S rDNA array and reveal a zone-specific organization of unit methylation that is associated with chromatin architecture.

## Introduction

Ribosomes carry out protein synthesis in all cellular organisms (1). Among their RNA components, 5S rRNA, a molecule of about 120 nucleotides (2), is present in every cytoplasmic ribosome and is transcribed by RNA polymerase III (Pol III) (3). Beyond its role in the ribosome, 5S rRNA can also assemble into the 5S ribonucleoprotein particle, which stabilizes p53 (4,5).

In line with these cellular roles, misregulation of 5S rRNA is linked to abnormal cellular states and human disease (1,6,7), and altered expression of 5S rRNA is observed in cancer and in immortalized cells (8–10). Regulating the production of 5S rRNA is therefore essential, all the more so since the copy number of 5S rDNA varies widely between individuals (11). Several factors that act on 5S rRNA transcription as a whole have been identified (3,12,13). However, at the unit level, it remains unknown whether active and inactive units are divided within the same array, what distinguishes them if so, and how such a pattern is distributed along the array.

DNA methylation, although a well-established mechanism of control in the 45S rDNA (14,15), has a controversial and species-dependent relationship with expression in the 5S rDNA (16–20). Earlier studies that resolved 5S rDNA methylation relied on methylation-sensitive restriction digestion (21) or on bisulfite sequencing of cloned fragments about 250 bp long (18), which are too short to cover the 2.2 kb human 5S rDNA repeat unit continuously. However, methylation analysis using recent long-read PacBio HiFi and Oxford Nanopore sequencing data, with reads of about 20 kb to more than 100 kb (22–25), can read the methylation of an entire unit, and even of multiple neighboring units, on a single molecule.

This capability makes long-read sequencing particularly well suited to the study of the human 5S rDNA array. The human 5S rDNA repeat unit is about 2.2 kb long (this study), and the units are arranged in a single tandem array on the long arm of chromosome 1 at 1q42 (26–28). Because these units are nearly identical, the 5S rDNA array long remained unassembled (29), and most studies have therefore measured its copy number per individual (11,30,31) or examined sequence variation among copies (32,33) without placing each unit at its position along the array. As a result, how units are organized and patterned within the array has received little attention. Long-read assemblies now make it possible to place each unit along the array, and thus to examine how sequence variation and the state of units differ with position and how units are regulated relative to one another.

Here 5S rDNA arrays were assembled from flank to flank from long reads and the assemblies were assessed for reliability. The long reads were then realigned to the assembled arrays, placing each unit within its molecule at its position in array. The array proves to be divided into zones by both sequence variation and methylation, and its units take one of two methylation states, low or high. Fiber-seq reads aligned to the same arrays further relate these states to chromatin accessibility and to footprints at transcription-factor binding sites, and ATAC-seq and ChIP-seq data from cell lines support this relationship.

## Materials and methods

### Data and cohorts

Long reads were taken from four public ONT sources, the HPRC (release 2 data (34) and, for 100 of the 302 individuals, Oxford Nanopore reads in the consortium’s working area but not part of a release (35)), GIAB (36) (its open data release giab_2023.05 (37)), the 1kGP ONT release of the 1000 Genomes Project Long-Read Sequencing Consortium (24,38), and cancer cell lines. The HPRC and 1kGP material is Epstein-Barr-virus-transformed lymphoblastoid cell lines. The reference set is 612 haplotypes from 310 individuals (HPRC 604/306, GIAB 8/4) as unique (individual, haplotype) keys, from a registry of 616 such keys from 313 individuals, which is the count the supplementary tables report. Supplementary Table S1 gives how each reference was obtained and with which assembler (34,39–41).

Four tiers inside the HPRC cohort are subsets of it: H0, the whole cohort (604/306); H1, the variant reference (557/301), dropping references invalid for variant calling and haplotypes whose reads come from outside the HPRC ONT submissions; H2, the methylation reference (510/280), on a six-condition slot quality rule covering placement agreement, covered fraction, depth and complete-unit count; and H3 = H1 ∩ H2 (490/274). Neither H1 nor H2 contains the other; the four counts are in Supplementary Table S1. Figures 2 and S2 are drawn on the 541 haplotypes of 299 individuals with a unit at every depth 1 to 40 from both junctions, 51,219 units, and panels by superpopulation on the 533 of those matched to the 1000 Genomes table, 295 individuals. The tier behind every panel is given in Supplementary Table S1.

Two methylation cohorts share no individual: H2 (510/280, 733,288 read-units, 45,544 units passing the filter below) and the 1kGP set of 18 haplotypes from 9 individuals. The 1kGP set is single-chemistry and single-basecaller (R10.4.1/LSK114, dorado 0.9.0, model v5.0.0); H2 mixes R9 and R10 with two basecallers, broken down in Supplementary Table S1. Absolute levels are therefore compared within one cohort, and between individuals only where the text says so. The consortium report on its first 100 samples (24) describes R9.4.1 material and states that later sequencing moved to R10.4.1, from which the 18 haplotypes used here come. In the 1kGP set, profiles along an array use only assembly segments whose two ends both reach flanking sequence, with no unit 31-mer within 500 bp of either segment end. Seventeen haplotypes have no ONT reads.

Fiber-seq was analyzed for K562 and GM12878, processed with different versions of the m6A caller (K562 with SMRT Tools v23, GM12878 unestablished), so the two lines are compared by the direction of the within-line contrast. The matched tumor-normal pairs include one produced as a reference material (42). That the Fiber-seq input is chromatin was checked on the data, m6A density in open segments exceeding that in nucleosomal segments 32.1-fold in K562 and 28.1-fold in GM12878. Haplotypes are labeled A and B; for a cohort individual these are fixed registry slots, for a cell line a naming convention, so cell-line labels bind to unit counts, stated wherever used.

Eight species were analyzed from Nanopore whole-genome data, five apes (Sumatran orangutan, Bornean orangutan, gorilla, bonobo, chimpanzee), zebra finch, zebrafish and *Arabidopsis thaliana*. Of 65 cataloged 5S arrays 49 were analyzed, the remainder being arrays of three to five copies and the single siamang array. In one bonobo sample the same flowcells basecalled with Guppy 6.3.7 and 6.3.8 differ by 0.13 in methylation level, so only the 6.3.8 flowcells are used for that species.

### Array references, read extraction and read placement

Cohort 5S reads were extracted per individual by streaming each BAM through one BBDuk (43) k-mer gate (BBTools 40.02, k = 25, hdist = 2, mkh = 2, both strands) that preserves the MM/ML base-modification tags (44), which are present on 597 of 616 haplotypes over 1,019,408 read-unit rows, the 19 without calls being the 19 with no ONT reads. Cell-line short reads were put through the same gate against the array reference with its AluY segments masked and 5 kb of flank on each side, at k = 21 and hdist = 0.

For individuals with a public assembly the array was cut from that assembly; otherwise it was assembled with hifiasm (45,46) (--ont) from that individual’s extracted 5S reads alone, except for two references from 1kGP-LRSC reads (24,38). Both routes apply the same cutting rule, the array-bearing contig being found by screening the assembly for eight 25-mers from four positions of the 5S gene on both strands. Two 50 kb probes cut from CHM13v2.0 (22) immediately outside the two array-to-flank junctions, chr1:227,694,884-227,744,883 proximal and chr1:228,025,901-228,075,900 distal, are aligned to the contig with minimap2 (-x asm20), and the span between their two inner ends is the array; a contig at which only one probe aligns is discarded. The routes differ only in the probe alignment required, at least 20 kb of matching bases for a public assembly and at least 5 kb for one made here. For a public assembly secondary hits are discarded (--secondary=no); otherwise they are kept (-N 20 -p 0.1), segments shorter than 4 kb are dropped and contigs whose lengths agree to within 200 bp are treated as one. Units are counted as 5S-gene blastn HSP spacings of 2,100-2,400 bp, the gene search using blastn (47) (-task blastn -dust no - word_size 9 -evalue 1e-5), with -task blastn, which finds the short repeat-embedded gene. Both array ends carry a truncated partial unit, which is excluded in every haplotype.

Alignment used minimap2 (48,49) with SAMtools (50) and BLAST+ (47); the two places where the pipeline ran carried different releases, and both sets are in Supplementary Table S1. Assembly-to-assembly alignment used minimap2 (-x asm20), and reads were placed with -x map-ont for ONT and -x map-hifi for PacBio HiFi.

Reads are placed on one reference holding both haplotypes together with the two 50 kb flank probes, so a read cannot be drawn to the wrong haplotype by the absence of the right one and a read reaching out of the array aligns to flank. Alignment used minimap2 (-a -- secondary=no -y), -y carrying the MM/ML tags, and only primary alignments are used; the two haplotypes are about 99.6% identical, so an ambiguous read carries MAPQ 0. A read counts as reaching the flank when at least 1 kb of it aligns to a flank probe within 50 bp of the junction, the 50 bp being the trough of an observed bimodal distribution of junction distance. Haplotype is then assigned by two routes. A read that reaches a junction is assigned by a vote at the positions where the two haplotype flanks differ, used only where at least five discriminating positions exist, the false-positive rate under permuted labels being 14.3% below five positions, 2.4% at five to nine and 0.0% at ten or more. A read lying wholly inside the array takes its haplotype from the primary alignment between the two haplotypes of its own individual at MAPQ ≥ 20, and that placement is tested by a map-margin difference: the read is placed as a chain of K consecutive units against the map of every individual in turn and kept only when the margin on its own individual’s map exceeds the best margin on any other map by at least 10. The difference is the discriminating axis, the median margin on another individual’s map being 1 against 25 on the right one. The chain length is fixed before the data are seen, K = 16 spanning at least 36 kb for whole-genome ONT reads, K = 12 and 26,880 bp for Fiber-seq ONT reads, and K = 4 with a margin difference of 5 and 8,960 bp for the shorter PacBio reads. Where the two routes disagree the chain is followed.

The unit frame is 2,240 bp with the transcription start site at 2,122: AluY 1-318, spacer-1 319-1,178, GA-rich 1,179-1,551, spacer-2 1,552-2,031, the 5′-flanking region 2,032-2,121 and the 5S rRNA gene 2,122-2,240 (119 bp, 9 CpG). A second frame of 2,241 bp with the start site at 2,123 is used in parts of the accessibility analysis, the gene body being 119 bp in both. Along the array, position 1 is the proximal end, the end nearer the centromere, and centromere positions were determined here from the annotated centromeric satellite of the T2T assemblies for the apes, CEN180 arrays (51) for *A. thaliana*, Tgut191A and Tgut716A arrays (52) for the zebra finch, and the satellite-density peak of the UCSC GenArk repeatMasker satellite track (53) for zebrafish. Depth is the position of a unit counted from a junction, depth 1 being the first complete unit next to it, and truncated units excluded. The frame and the axis direction that apply to each panel are given in Supplementary Table S1.

### Sequence variation

Variants are called per unit against a majority-allele reference unit built with a > 0.5 rule, which gives 6.87 variants per unit over H1. Unit sequences are compared by pairwise global alignment with parasail 1.3.4 (54) (nw_trace_striped_16, match +2, mismatch −3, gap open 10, gap extend 1). For the origin-class panels each (haplotype, variant) pair is assigned one of three classes, using haplotypes of at least 30 units: multi-individual shared when the variant is carried by at least 10% of those haplotypes, within-array multiple when it occupies at least two units of that array, and single-unit when it occupies one. Trimming the longest-read span ratio at 0.45, the fifth percentile of that ratio over the cohort, defines the reassembly test of Figure S1. A transmitted pair is a parent-offspring haplotype pair at a mean variant Jaccard of at least 0.95, units being matched by aligning the two arrays from either end with a shift of up to 5 units and keeping the alignment of highest mean Jaccard; the non-transmitted comparison uses the other haplotype of the same parent, and the hg003 reference was inferred and not used for transmission. Co-sharing around a rare variant is measured in the central window (0.2-0.8 of the array) of haplotypes with at least 23 window units, over pairs of units that both carry a variant A of cohort unit frequency below 1% against control pairs at the same separation in which only one unit carries A; over substitutions and deletions only. A terminal-type unit carries at least 10 of the 12 diagnostic variants of Figure S2I, neighbor and sandwiched-unit comparisons are made against controls at the same array depths in arrays that carry no terminal type, and the terminal-to-terminal class is omitted (3 neighbor pairs).

The variant-based zoning of Figure 2 has two steps and methylation β is not among its features. Boundaries were first placed in each haplotype of at least 40 units by binary segmentation (55) on the first five principal components of unit features centered within each haplotype, with a cost equal to the within-segment sum of squares, a penalty of 1.0 × d × log n (d = 5, n = units in that haplotype) and a minimum segment of three units, the features being 20 variant characters (criterion B) or individual variants carried by 0.5-99.5% of units (criterion A). Cohort zones were then set by binary segmentation of the profile of the 20 variant characters averaged at each depth from 1 to 40 on each side, z-standardized along depth, with a penalty of 1.0 × 20 × ln 40 and a minimum segment of two depths, which cut behind depths 3 and 17 on the proximal side and behind depth 10 on the distal side. Each unit was also described by its counts of variant characters standardized against middle units (deeper than 30 units from both junctions): the proximal-ness and distal-ness axes are the unit vectors from the mean of up to three randomly chosen middle units per haplotype to the mean of units within depth 3 of the proximal or distal junction, the score of a unit is its projection on an axis, and the threshold of each axis is the 95th percentile of the scores of those sampled middle units, so that 5% of middle units are expected to exceed it and the two quantities are reported as the fraction of units above the threshold minus 0.05. Compartments were obtained by clustering units with k-means (56) (k = 12, 20 starts, random state 0) on the same position-free features, a cluster being kept when its distribution along the array, in 20 bins of relative position, departed from uniform by more than the 97.5th percentile of 200 shuffles of unit order within each haplotype.

### Methylation calling and the state cut

Methylation calls are those of the submitting consortia, basecalled with dorado (57) or guppy (58). Every CpG call is a hard call, counted 1 when the methylation likelihood reaches ML ≥ 128 and 0 otherwise. A read-by-unit observation was kept when at least 150 of the unit’s 199 canonical CpG positions were called and at least 90% of that read’s calls fell on canonical CpG positions, and haplotype-level tables additionally require 5 such reads per unit; the 0.9 is a call-purity ratio and so a check on placement. β is the call-weighted mean Σ5mC ⁄ Σcalls of one read × unit. A haplotype-by-unit row enters the positional analyses only when that unit carries at least five reads, and the array-length strata need at least 40 units per haplotype.

Unit methylation is split at β = 0.508, the valley of the unit-level distribution in H2 (47,467 units), and that cut was adopted for every analysis here. A valley is a property of the ruler and of the aggregation unit, so a cut is quoted with both; over three measures, three state definitions and two platforms all 18 combinations keep the same sign. The proximal-to-distal span of the hard-called β mean is +0.4020 overall against +0.3895, +0.3979 and +0.4250 in the R9.4.1, the R10.4.1 and the unrecorded stratum, so H2 is used uncorrected. Three further cuts are used and none is interchangeable with it: β = 0.4611 in the 1kGP set, fitted by the EM algorithm (59) and split by likelihood ratio; β = 0.5 on gene-body CpGs, which labels units in the cell-line comparisons and agrees with the whole-unit label for 91.6% of units; and β = 0.35 in the cross-species comparison, where the distributions are binned differently for each array and no common valley exists. For DXZ4 the two modes are the active and the inactive X chromosome, and no cut is used.

Within each array every repeat unit carries two 1-based positions, ap counted from the proximal end and ad from the distal end, with ap + ad = N + 1 in an array of N units. Each unit was assigned to whichever end it lies nearer and the array was cut into five zones whose boundaries are fixed in units and not in fractions of array length: proximal 1-3 (ap ≤ 3), proximal 4-10 (4 ≤ ap ≤ 10), distal 1-10 (ad ≤ 10), distal 11-21 (11 ≤ ad ≤ 21), and the interior. For each haplotype the low-methylation stretch at each end was also delimited by a single change point in gene-body CpG methylation, located by an exhaustive scan over every split position, taking the position at which the sum of two per-segment binomial log-likelihoods is largest, with the segment required to run from low to high methylation and with no penalty term and no minimum segment length. Array types are those defined by carriage of variant tokens, and the methylation comparison keeps the haplotypes of those types that also fall in H3 and carry a unit with at least five reads in every zone, the distance between two haplotypes being the mean |Δβ| over the five zones with pairs within one individual excluded. In the trio comparison the methylation of a haplotype in a zone is the median of its unit β over the units of that zone, taken on units with at least two reads.

A block of one state is a stretch of one state within a run of at least three consecutive units on one read of H2 that has a β for all six sub-unit elements, an element being high-methylation above β = 0.508, and stretches touching either end of the run are excluded; spacing is the distance in elements between successive starts of blocks of the same state within a run, against one in six expected by chance. A boundary window is six consecutive units on one read of H2, each with at least 20 CpGs, in which three units of one state are followed by three of the other at a unit β of 0.508, read from the distal toward the proximal end, and methylation across a boundary pools the hard calls of all windows in 50 bp bins of the unit coordinate; because windows are chosen by these six unit states, part of the contrast between the two central units comes from the selection; the example molecules are drawn from windows whose three high units all have β of at least 0.70 and whose three low units all have β of at most 0.25. A step is the difference in β between two adjacent complete units on one read going inward from the array end, within 25 units of the distal end of 5S rDNA and within 10 units of the proximal end of D4Z4, a read counting as rising when its steps sum to more than 0.15, and the signed steps of all rising reads are divided into three classes by absolute size (below 0.1, 0.1 to below 0.3, and 0.3 or more). In the 1kGP set the crossing is the first adjacent pair of units that passes β = 0.4611 from below, at position 3 or more from the array end. Element-level mixtures use a two-component binomial mixture fitted by the EM algorithm (59) without using the unit’s class, an element entering the fit when it has at least 12 CpGs for AluY, 30 for spacer-1, 12 for the GA-rich segment, 20 for spacer-2, 6 for the 5′ flank and 5 for the 5S gene, and a unit is counted when all six elements were assigned.

Dispersed 5S-like loci were taken from the ONT reads of 307 individuals that the read gate had extracted from outside the array. Two concatenated copies of the repeat unit were aligned to each read with blastn (47) (-task blastn -word_size 11 -dust no -evalue 1e-5), with the 119 bp 5S gene as a second query (-word_size 9 -evalue 1, at least 40 bp). Non-overlapping segments of at least 50 bp were kept in order of bit score with at most 20 bp of overlap, segments within 2,000 bp on the read form one locus, and a locus needs at least 50 bp in spacer-1, spacer-2 or the 5S gene. Each locus with up to 10 kb of read on either side was placed on CHM13 with minimap2 (48,49) (-x map-ont -c --secondary=no), and loci with a mapping quality of at least 20 were joined into sites by single linkage within 3 kb on each chromosome. The analyses use loci covering at least 50 bp of the 5S gene and sites with at least 200 such loci from at least 10 individuals, which gives 33 sites; one site overlapping the array itself (chr1:228.00 Mb) was removed, leaving 29 sites that carry at least three elements. Locus β is the hard-called methylation over the CpGs of spacer-1, spacer-2 and the 5S gene of one locus, which needs at least 10 such CpGs. For each site locus β was binned at 0.02, the valley being the emptiest bin between 0.2 and 0.8 and the low and high modes the fullest bins below and above it; a site is bimodal when the low mode is below 0.35, the high mode above 0.65, the share of loci above the valley between 5% and 95%, and the valley holds at most half the density of the smaller mode. The flank tone of an individual is the mean methylation of the CpGs on the same reads lying more than 500 bp outside the 5S-like segment, and for the comparison of single molecules within one individual flank beta is the same quantity measured on one read, for reads carrying at least 20 such CpGs, a molecule being one dispersed locus on one read. Array read-units and dispersed loci were assigned to states by one rule, β at 0.508 on at least 10 CpGs in a calling window of positions 319-1178 and 1552-2240 of the unit.

In the cross-species analysis the array is the unit of analysis. A unit is complete when it carries at least a per-array minimum of CpG calls, min(20, max(3, round(0.5 × the median CpG per unit of that array))), and when at least 90% of its reference CpGs are observed; in zebra finch the maternal and paternal copies of one array are pooled by summing calls, and for *A. thaliana* CG, CHG and CHH were measured separately, the β reported for that species outside Figure S11C being the CG value. The end slots are the first and the last repeat unit of the array and the interior is the part beyond the twentieth unit from either end, defined only for an array longer than 40 units, and flank windows pool at least 300 CpG calls. For the within-unit profiles the units of an array are split into a high-methylation and a low-methylation group, by the likelihood ratio of two fitted components where the array is bimodal and otherwise at the cut whose achieved low fraction is closest to the human 5S low fraction (*f* = 0.2191, 5,619 of 25,644 units in the 1kGP set), and no unit is discarded. Profiles are coverage-weighted sliding means over ±min(50, max(5, unit length/20)) bp, widened locally where a window would hold fewer than 40 calls, and the gene window is the last 120 bp of the unit. Ashman’s *D* (60), in the unequal-variance form (61), is √2 |μ1 − μ0| / √(σ0² + σ1²) from two Gaussian components fitted to unit β by the EM algorithm (59).

### Fiber-seq, footprints and cell-line chromatin

Accessibility was measured by Fiber-seq (62). The cohort is HPRC Fiber-seq, 52 individuals on ONT and 52 on PacBio; on PacBio the m6A panels rest on 48, because the submitted calls for two individuals carry 5mC only and two further individuals were held out whose m6A was called with fibertools-rs (63) rather than with jasmine (64). The per-individual tests also require at least 500 called bases per zone per state, and two PacBio haplotypes whose read unit numbering disagreed with the canonical array map are held out of the per-unit statistics. The two platforms are kept apart, m6A labeling efficiency differing between them, and agreement in direction is the replication.

Nucleosome and MSP (methyltransferase-sensitive patch) segments were called with FiberHMM 2.16.8 (65–67) (--enzyme hia5), the per-platform nuc_recall_policy being given in Supplementary Table S1. A later 3.x release retires both options and changes the Hia5 nucleosome calls; phase_nrl is estimated per sample, not fixed, and the values are in Supplementary Table S1. The fiber stacks of Figure 5A were called instead with ft add-nucleosomes (fibertools-rs 0.13.0, default parameters) (63), which gives a median segment matching one nucleosome where the HMM merges neighboring particles.

Three metrics are used, each with its own denominator. m6A fraction at a base is the number of molecules carrying m6A there divided by the coverage at that base. Large-MSP density is the bases covered by an MSP longer than 150 bp (68) divided by the called bases, the 150 bp being the methods value of the reference work (68) rather than one derived here and set in that work on fibertools-family segments (63), so the threshold is a reference value while the segmentation is not; the MSP length distribution in these data shows neither a mode nor an inflection near it, the median patch being 15 bp on ONT and 25 bp on PacBio and patches longer than the threshold accounting for 1.1% and 2.9% of patches, and the direction of every comparison is unchanged across seven thresholds spanning 100-300 bp. Nucleosome occupancy is nuc_bp / (nuc_bp + msp_bp) over called bases; nucleosome positions are inferred from the m6A pattern on the same molecule, so analyses removing the nucleosome contribution bound it from above. Molecules carrying a low-methylation unit take up m6A 2.80-fold more readily than molecules carrying a high-methylation unit, so raw comparisons between the two states cannot be separated from labeling efficiency; where the comparison matters it is paired within one molecule, using only molecules that pass through both a low- and a high-methylation unit, so that exposure to the enzyme and, the two unit classes being the same repeat sequence, adenine density are identical. Such molecules are 31.8% of ONT and 20.9% of PacBio molecules and are concentrated in the transition zones.

Footprint positions were found by a blind scan. m6A-protected intervals called by FiberHMM 2.16.8 were counted at every offset of a grid across the unit in size classes of 10 bp, within a window fixed before the scan to −150 to +118 relative to the transcription start site. The grid is smoothed with an unweighted moving average whose width and edge handling differ by panel, and a ratio is shown only where the permutation null is itself above a floor; those per-panel values are given in Supplementary Table S1. A second, caller-independent readout is the ratio of raw m6A call density in a flanking window to that in the core window. The offsets of those two windows around a center are those of Dubocanin et al. (68), and the center is taken from the caller output, independent of the data being tested.

The cell-line arrays were assembled with hifiasm 0.25.0-r726, K562 with --ul (69) from array-region HiFi reads plus the ONT 5S reads of BioProject PRJNA1371648, GM12878 without that flag (45) from GIAB HG001 (36) PacBio HiFi reads of the array region (70).

Fiber-seq was analyzed for both lines on all reads from each run, array placement discarding the majority of K562 reads; those pools keep every read touching the array, use no read placer, include nucleosome-covered bases, and the units were cut by anchor sequence. Nucleosomes in these pools were called with FiberHMM 2.16.8 (--enzyme hia5, conservative nuc_recall_policy), the per-sample phase_nrl being given in Supplementary Table S1. The two Fiber-seq files differ in kind: for K562 the submitted BAM of the archived run was read, so its base modification calls are those made when the data were produced, whereas for GM12878 the processed FIRE file of that release was read, whose m6A was called with fibertools-rs 0.1.4 (ft predict) and whose accessibility calls were made with fibertools-rs 0.3.8 (ft fire). The coordinates of neither file were used; for both lines the 5S reads were placed on the array reference used here.

A zone-exclusive marker is a sequence variant present only in units of one zone. Short fragments from ATAC-seq and ChIP-seq were passed through an alignment gate that keeps only reads matching the array and not its dispersed relatives, a read having to score highest on the array against dispersed 5S decoys and higher than any position outside the array in CHM13. A zone was scored as Σ(observed marker-allele reads) ⁄ Σ(read depth × q), q being the marker-allele fraction in whole-genome sequence from the same line through the identical gate, summed over the zone, read depth varying by orders of magnitude between markers. The interaction statistic is the proximal-to-interior ratio of the target divided by that of the control track. The control track of each cell line is one pooled track, in K562 ENCSR000FAK (input) and ENCSR000EHM (IgG), in GM12878 ENCSR000EYX and ENCSR000EAF, so the expected proximal count of the denominator is one number shared by every target, 1.086 in K562 at base quality 20 or more. An interaction is reported as NC (not computable) when either zone has fewer than two markers scored in both tracks. Two counting rulers appear in the figures and are labeled in each panel, a count without a base-quality filter and a base-quality ≥ 20 count. ChIP-seq targets were included, before any outcome was examined, when their expected proximal count over the proximal markers was at least 3 and the control track reached at least 1. Because an assembly can under-represent copies, zone exclusivity was also tested on reads: contigs were assembled from marker-allele reads alone, requiring agreement on the marker’s unit slot, and each read was then classified as lying at its original site, at another site in the same zone, outside the assembly, or ambiguous.

### Statistical analyses

Tests are two-sided unless stated otherwise, intervals in parentheses are 95% intervals, randomized procedures are seeded in the source, and where a quantity was computed in more than one way both values are reported. Copy number is compared at the haplotype level by the Kruskal-Wallis test (71) with Dunn’s test (72) over all ten pairs of superpopulations and Bonferroni adjustment within that set, Dunn’s test computed from the rank sums directly; the two haplotypes of one individual being dependent, the same omnibus test at the individual level is given with it. Superpopulation labels come from the 1000 Genomes pedigree of 3,202 samples, agree with the phase 3 panel at all 276 shared individuals, and four individuals without a label are excluded. Paired comparisons of tissue, culture and tumor use the Wilcoxon signed-rank test (73), from the exact null for the per-individual tests and the normal approximation for the per-molecule tests; with seven pairs the smallest attainable *P* is 0.0156, stated wherever reached. The unpaired estimate of Figure S7B is tested over individuals, not over pooled units, a state running in blocks over many units. Zone-resolved recounts of the transmission comparison are corrected across the five zones by the Benjamini-Hochberg procedure (74) and the smallest q is reported, and presence/absence comparisons in the shallower cell line use Fisher’s exact test (75) with a one-sided 95% lower bound on the ratio, the zero cell making the ratio unbounded above.

Permutation tests state the number of permutations, and a value at the limit of the grid is given as a bound. The footprint null was built at three counts, 200 for the null mean, the curves and the thresholds, 1,000 for the blind scan, whose *P* values reach a floor of 1/1,001 and are reported as ≤ 0.0010, and 2,000 for the confirmatory tests; multiplicity was handled by taking, for each permutation, the maximum over the whole grid, and the thresholds, each the 95th percentile of the 200 maxima, were 2.94 for the caller-based axis and 1.03 for the caller-free axis, derived within each methylation stratum and not from the pooled scan. The distance-dependence of unit similarity is tested against a null in which unit order is shuffled within each haplotype, by an extreme-rank-length envelope test (76) with 2,499 simulations plus the data, so its *P* value has a floor of 1/2,500. The identity background against which unit-pair sharing is normalized is φ (77), the identity coefficient between units on different homologous chromosomes, 0.003134 over 2,083,965,584 unrelated pairs, and the exchanged-patch length is estimated by maximum likelihood under a gene-conversion tract-length model (78) and reported as a lower bound. Bimodality in the cross-species comparison was called by a parametric-bootstrap likelihood-ratio test (79) of one against two beta-binomial components behind six gates, testable, significant, resolved, minor-component size, peak separation, and reproduction across five split halves, the procedure of McLachlan having been developed for normal mixtures and applied here to beta-binomial mixtures. The population-exclusive token counts are medians over 200 subsamples matched on group size, and the AFR skew is tested by permuting population labels between individuals, the two haplotypes of a person moving together, over 1,000 permutations of 50 subsamples each on 503 haplotypes of 276 individuals, and the rarity-clumping comparison uses 999 permutations. Bootstrap intervals resample the unit stated with each measurement, 2,000 resamples of markers (80) drawn separately within the proximal and the interior zone for the zone ratios and their interactions, 400 resamples of haplotypes for the compartment regressions, 500 for the co-sharing comparison and 1,000 for the terminal-type exchange estimates.

### Code and software availability

All analyses were carried out in Python with NumPy (81) and SciPy (82). All analysis code, together with the result tables behind the figures and a manifest of inputs and outputs, is openly available at https://github.com/H-minKim/human-5S-rDNA-array under the MIT license and is archived with a permanent DOI at https://doi.org/10.5281/zenodo.24059050. Four result tables are too large for the repository and are in that same Zenodo record, P10_FRAME_ALL.tsv, P10_CPG_ADD3.tsv.gz, L1_READ_UNIT.parquet and P10_CPG_ALL.tsv.gz, and tables/README.md in the repository names them with their sizes. The transcription start sites on the annotation tracks rest on the published GM12878 GRO-cap data of Core and colleagues (83) (GEO accession GSE60453, the GRO-cap subseries of GSE60456), the 5S annotation tracks use the mature 5S rRNA (EMBL X12811) with the RefSeq 5S and U2 entries (NR_002716), the D4Z4 4q track uses the DUX4 open reading frame (NM_001306068) with the two RefSeq D4Z4 transcripts (NR_147172, NR_147173), and the zebra finch arrays were taken from the diploid bTaeGut7 assembly (84) (GCA_048771995.1 maternal plus Z with GCA_048772025.1 paternal autosomes), the array count being a property of that assembly version. Parts of the analysis were prepared with the assistance of a large language model (Anthropic Claude), used under the direction of the author, who checked every number reported here against the tables it was taken from and is responsible for all of it. No figure or image was generated by a model, and every panel is drawn from data by the deposited code.

## Results

### A flank-to-flank array reference with long-read sequencing data

The 5S rDNA array lies at chr1q42.13 (27–29) and is a tandem repeat of a 2,240 bp unit, tens to hundreds of units long in a single haplotype (Figure 1A). The 5S rRNA gene sits on the minus strand of chr1, so when the reference genome is laid out with coordinates increasing to the right, transcription within each unit runs from right to left, that is from the distal toward the proximal end.

**Figure 1.**
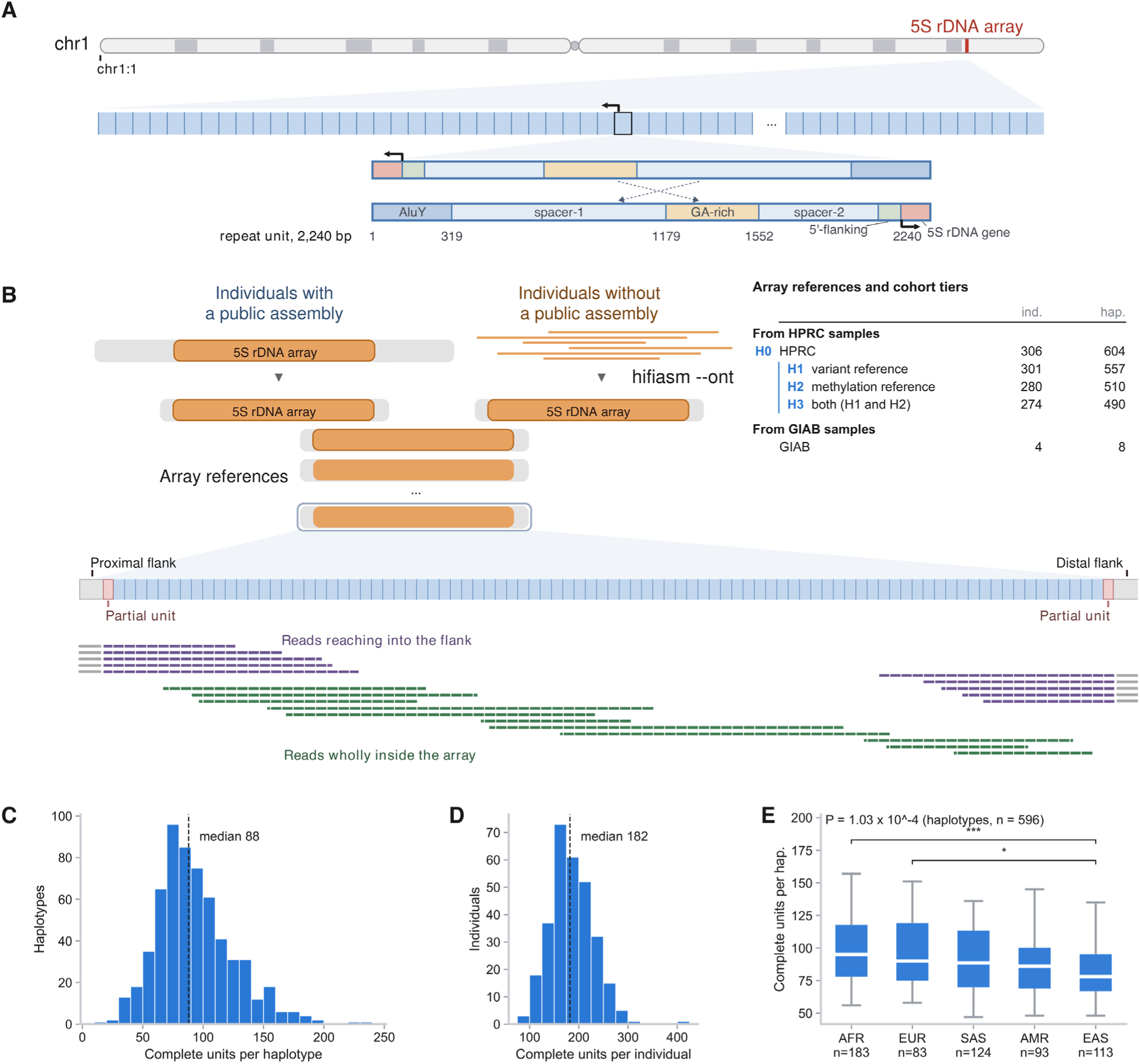
A flank-to-flank array reference using long-read sequencing data. (A) The repeat unit and its parts. The 2,240 bp repeat unit and its six sub-unit elements, AluY 1-318, spacer-1 319-1,178, GA-rich 1,179-1,551, spacer-2 1,552-2,031, 5′-flanking 2,032-2,121 and the 119 bp 5S rRNA gene 2,122-2,240, with the transcription start site at 2,122. (B) Construction of the array references, read placement, and the reference set. One array reference is built per haplotype and long reads are placed on it, and the table inside the panel gives the reference set and the cohort tiers from which the per-panel denominators are drawn. (C) Copy number per haplotype. Dashed line, the median. n = 604 (HPRC). (D) Copy number per individual. Dashed line, the median. n = 298 (HPRC), 8 individuals with only one haplotype assembled are excluded. (E) Copy number by 1000 Genomes superpopulation. Boxes, the interquartile range with the median. Whiskers, the 5th and 95th percentiles. *P* = 1.03 × 10⁻⁴, the Kruskal-Wallis test across the five groups, and the brackets above the boxes are the pairs that survive Dunn’s test over all ten pairs with Bonferroni adjustment, *** for *P* < 0.001 and * for *P* < 0.05. n = 596 labeled haplotypes (HPRC).

Seeing the landscape of the array as a whole first required the array in one piece, flank to flank. In individuals with a public long-read assembly, including those of HPRC, the main cohort here, two 50 kb stretches of CHM13 flanking sequence lying outside the junctions served as anchors and the sequence between them was cut out as the array. In individuals without a public assembly, the ONT reads carrying 5S array sequence were collected and assembled separately with hifiasm --ont (45,46), and the same rule was then applied to that assembly (Figure 1B). This yielded array references for 306 individuals and 604 haplotypes of HPRC and 4 individuals and 8 haplotypes of GIAB. The 5S rDNA array reads, ONT and PacBio HiFi alike, were then realigned to these references. A read carrying a flank was anchored on that flank. A read lying wholly inside the array was assigned to a position only where the variants it carries make that placement unique (Figure 1B).

Assemblies of this kind had not been validated for the 5S rDNA array. The next question was whether these assembled arrays are reliable. The unit count per array came first. Some haplotypes carry a read that crosses the array in one piece from flank to flank, and these reads were taken as an answer key against which the unit count of the assembly could be compared. Of the 195 haplotypes that have such a read, 181 (92.8%) agreed exactly and 194 (99.5%) agreed to within one unit, and only one disagreed by more (Figure S1A). Because the spanning read had itself gone into the assembly, a further test asked whether reads that span only part of the array length are by themselves enough to build a correct assembly: every read was trimmed to 0.45 of the array length, the fifth percentile of the fraction of the array covered by the longest read in these data, and the haplotype was then reassembled (Figure S1B, C). This test was run on the 32 individuals and 64 haplotypes chosen as the individuals in which both haplotypes have a spanning read. Of the 64 haplotypes, 62 (96.9%) were reassembled, and all 62 of those (100%) matched the length of the spanning read (Figure S1D). Reads covering only part of the array were therefore enough to rebuild the whole of it, and every rebuilt array came out at the length the spanning read gives. The next question was whether this reference is reliable at the sequence level, that is whether the variant content inside those units is reproduced by a consensus of PacBio HiFi reads. Variant recall was 99.54% for the public HPRC assemblies (n = 313) and 98.19% for the hifiasm assemblies made here (n = 119), which are built from ONT reads (Figure S1E). So even when a reference built from ONT reads was taken as the backbone and the consensus was built from PacBio reads, the sequence recovered was almost identical to the reference.

Structural error came next. Gross misassembly such as an inversion or a block move was sought with the structure test (Figure S1F), and because that test had low sensitivity to the insertion or deletion of a single unit, a finer test was added, the insertion and deletion test, which measures the distance between two unique anchor units (Figure S1G). In the structure test a constructed inversion of six units was caught plainly, but in the real arrays no inversion, block move or duplication was detected in any of the 591 haplotypes the test reached. Similarly, the insertion and deletion test flagged nothing in the great majority of the arrays. Of the 591 haplotypes, 579 (98.0%) carried no flagged site, and the remaining 12 haplotypes carried 13 flagged sites between them, one of the 12 carrying two of the sites. Twelve of the 13 sites were flagged by the insertion and deletion test and one was carried over from the earlier unit-count pipeline, and the structure test flagged none. Each flagged site was then re-examined read by read. Of the 12 haplotypes, six carry a true deletion or insertion and six are mosaic, with reads supporting the reference form and the event form at the same site, which may reflect the gain or loss of a unit during cell division (Figure S1H). Two references were corrected on this evidence, one haplotype of HG00133 from 153 units to 149 and one haplotype of HG03270 from 78 to 77. Worked examples are shown in Figure S1I.

On the results of these tests, the length and the sequence of the 5S arrays obtained in this way lie within a reliable range. On that footing the array references were first used to look at copy number. Its distribution at the level of the haplotype (median 88 units, n = 604, Figure 1C). Summed to the level of the individual, the median was 182 units (n = 298, Figure 1D). Both values fall within the ranges other studies report (32,85,86). The distribution of array length at the haplotype level differed between populations (Kruskal-Wallis test across the five superpopulations taken together, Figure 1E).

### Zoned sequence features along the array

Using the complete array assemblies described above, the sequence landscape of 5S rDNA was examined next at two levels that have received little attention before, that of the individual unit within a contiguous run of units and that of the array as a whole, to describe how sequence variation is organized along the array. Variants were called in each unit against a reference unit built from the cohort majority allele (> 0.5) at each position (Methods). The distribution of variants differed between unit elements. The per-base variant rate was high in the GA-rich region and spacer-1 and low in the 5S gene, the 5′ flank, spacer-2 and AluY (Figure 2A). To assess whether these calls reflect real sequence rather than assembly or calling errors, arrays assembled separately from parents and offspring in trios were compared. In the five parent-offspring pairs whose arrays matched along their whole length, 436 of 447 units (97.5%) were identical in sequence to the unit at the same position in the parent (Figure S2A).

The landscape within the array came next, namely how the units of one array differ from each other and how these differences are composed. Units of one array resembled each other more than units of different arrays (Figure 2B). The resemblance between the two homologous arrays of an individual exceeded that between unrelated individuals overall but was nearly the same as that between unrelated individuals of the same subpopulation. The resemblance of homologous arrays therefore reflects mainly a shared population origin, and arrays from one population share part of their variation. This within-array homogenization (concerted evolution) was stronger between nearby units (Figure 2C), proceeded through the exchange of patches within repeat units (Figure 2D) and differed in extent between elements (Figure S2B). To exclude the possibility that identical variants arose independently in different units at mutation-prone sites, pairs of units that share a rare variant were examined. These pairs also shared other variants more often than control pairs at the same separation (Figure S2C), so shared variants are better explained by copying between units than by recurrent mutation. The probability that a whole element was identical between two units of the same array was highest for the 5′ flank and the 5S gene (Figure S2B). These patterns agree with the earlier observation that the 5S gene is depleted of common variants and of gene conversion tracts (32). Units within an array are thus homogenized by concerted evolution.

At the level of the whole array, however, the frequency of variants was not constant along the array. To examine this further, variants were divided into three classes. Variants carried by at least 10% of haplotypes were classed as multi-individual shared. Of the rarer variants, those present in two or more units of one array were classed as within-array multiple and those present in a single unit as single-unit. Both the number of variants per unit and their origin composition changed with position in the array (Figure 2E). Within-array multiple variants are copies of one variant spread over several units of an array and, together with the resemblance between units (Figure 2B and C), provide evidence of within-array homogenization. Variants were also divided by substitution class (Figure 2F, left). To test whether positional differences remain once the effect of local homogenization is removed, single-unit variants were analyzed separately and again differed with position (Figure 2F, right). Positional differences within the array therefore not only reflect regional character shaped by local homogenization but are also present in rare variants confined to one unit. Variants present in a single unit thus also differ with position in the array.

Along the array, the proximal and distal ends each had a distinctive variant character (Figure 2G and H). Notably, as arrays lengthened, nearly all of the added length went to the central compartment (Figure 2I). Across length bins, the numbers of proximal and distal units stayed nearly constant, whereas the central region scaled with array length (Figure 2J). Unbiased change-point segmentation, which places boundaries in each haplotype separately, gave the same result (Figure 2K). The proximal and distal ends are thus regions of a nearly fixed number of units, irrespective of array length. On the proximal side the split gain marked clear boundaries at depths 3 and 17, dividing this end into proximal 1-3 and proximal 4-17. On the distal side the split gain was flat over depths 7 to 11, and depth 10 was taken to define distal 1-10 (Figure S2D, with the zone map in Figure 2K, bottom). The leading variant classes differed between these zones, among all variants and also among single-unit variants, in which C>T and G>A at CpG sites led in proximal 4-17 and the central compartment and C>T and G>A at non-CpG sites led in the distal end (Figure S2E). In every zone, the variants that distinguished zone boundaries were mostly multi-individual shared (Figure S2F).

The CpG bias of C>T substitutions illustrates differences within the array. After removal of multi-individual shared variants, the CpG bias of C>T substitutions on the distal side differed from that in other regions and was reversed (Figure 2L). The tendency of variation thus differs between regions of the array, and the array can be divided into zones by its variant character.

Differences between arrays came next. When haplotypes were clustered by carriage of variant tokens in the four zones (proximal 1-3, proximal 4-17, central and distal 1-10), the four largest clusters (28 to 99 haplotypes) each spanned four or five populations, whereas nine of the remaining ten clusters drew at least 80% of their haplotypes from one population (Figure 2M). These array types suggest that combinations of variants are inherited along population history and persist despite homogenization within arrays, with widely shared types possibly older than the population-restricted ones. The tokens that separate each cluster, and the number of zones in which they do so, are shown in Figure S2G. Tokens confined to one population were most numerous in the African population. Over 200 subsamples to the size of the smallest population, the median number of such tokens was 162.5 in AFR and 9.0 to 22.0 in the other populations. The AFR median exceeded that of all 1,000 permutations of population labels between individuals, giving *P* = 0.001, the smallest attainable value (Figure 2N). This agrees with the higher genetic diversity of African populations (87).

At the distal end, the variant count per unit peaked at depths 2, 5 and 7 (Figure 2E). These peaks came from units carrying a distinctive combination of variants. Units carrying at least 10 of 12 diagnostic variants define the terminal type, and each diagnostic variant was carried by at least 96% of terminal-type units (Figure S2I). Terminal-type units appeared from distal depth 2 onwards (Figure S2H) and were mostly 2 or 3 units apart (89% of 414 gaps). Gaps of 1 were rare (0.7%, against 28% when terminal-type units were placed freely, Figure S2J), and this depletion was not seen for other minority units (Figure S2K). Terminal-type units rarely exchanged sequence with neighboring units (Figure S2L and M). This low rate of recombination between terminal-type and non-terminal-type units could explain the depletion of gaps of 1 between terminal-type units, because forming two adjacent terminal-type units from a terminal-type unit with non-terminal-type units on both sides may require recombination between the two unit types. Terminal-type units of different individuals resembled each other even when the 12 diagnostic variants were excluded from the comparison, suggesting a common origin (Figure S2N). Carriage was lowest in African haplotypes (5.4%, against 41% to 55% in the other populations), and seven of the nine African carriers had terminal-type units at only one or two depths (Figure S2O and P). In cell-line assemblies that carried terminal-type units, these units were also spaced without gaps of 1 (Figure S2Q).

Taken together, these results describe variation in 5S rDNA units at the level of the array. Long-read assemblies resolved the combination of variants carried by each unit in its contiguous context. At the unit level, this showed how closely each unit resembles its neighbors and revealed the terminal-type unit, a unit with a characteristic combination of variants. At the array level, variation between the units of an array was not a uniform product of concerted evolution but had a distribution characteristic of each region. By variant content, the array had characteristic proximal and distal zones whose lengths were relatively fixed and not proportional to the total length of the array.

### Positional methylation of the 5S rDNA array

The long-read sequencing data used here match methylation data directly with sequence data, which allows the distribution of methylation across the 5S rDNA array to be asked next. Flank-to-flank methylation of the 5S rDNA array shows a pattern clearly separated from the surrounding flank, low at the proximal and at the distal end and high in the interior (Figure 3A), in the way that CpG methylation gradients have been read along the D4Z4 macrosatellite with long reads (88,89). One of those studies also reported an asymmetric methylation gradient at the two ends of the 5S rDNA array of chromosome 1 in a single control individual (88) and reported for the repeat content of the complete reference (90). The ends are the least methylated part of the array. The low methylation also reaches into the flank, the proximal flank being low 4 to 12 kb out and the distal low stretch continuing past the array into the flank. The same contour appeared in every data source (Figure S3B, C). Of note, the proximal partial unit was almost identical in sequence to a repeat unit and yet appeared far more methylated than the proximal unit beside it. Figure S3A shows CpG density over the same range.

**Figure 2.**
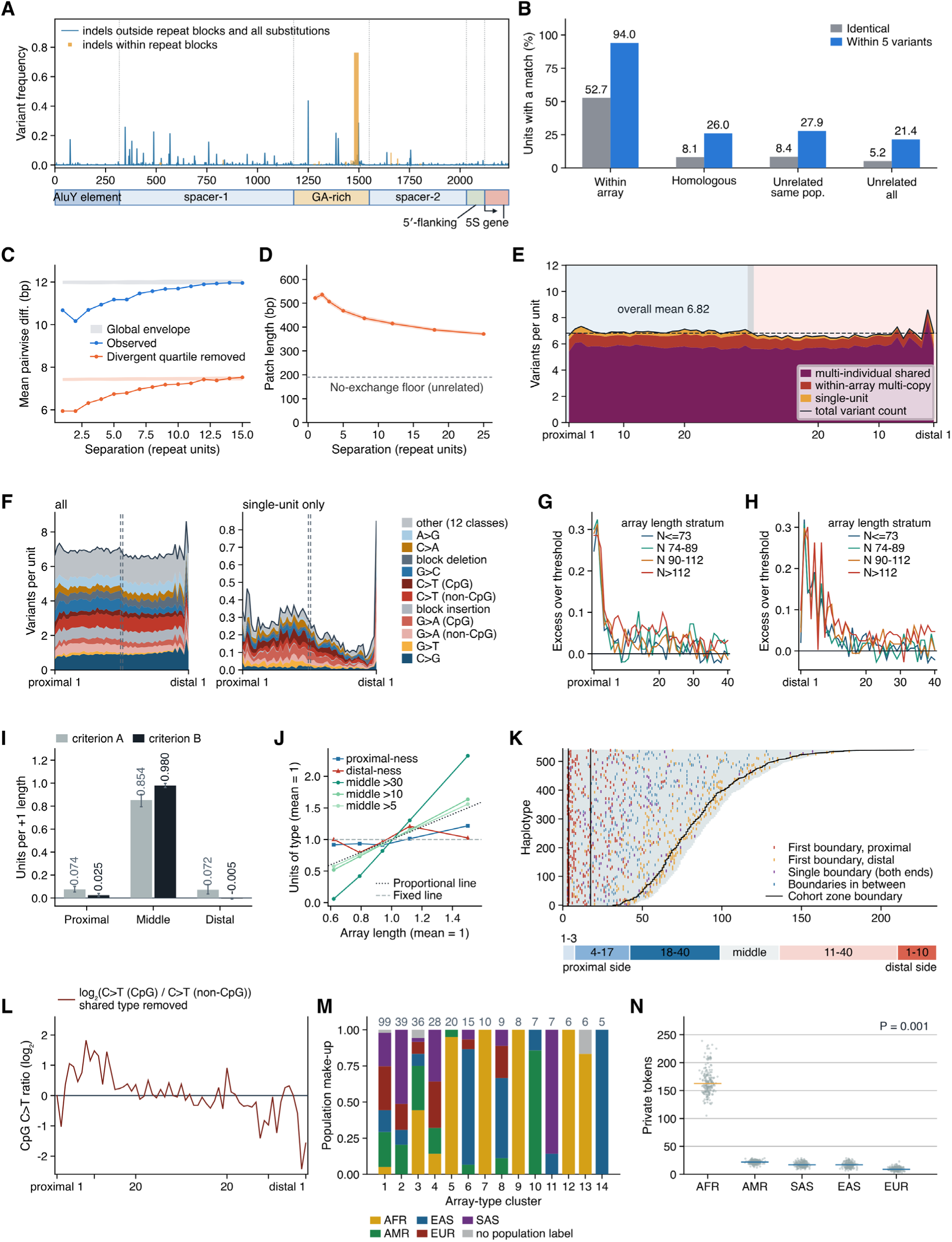
Sequence variation within the 5S rDNA unit and along the array. (A) Variant frequency along the unit. Fraction of units differing from the majority-allele reference unit at each position. Blue line, substitutions and, outside repeat blocks, insertions or deletions, per base (1,936 positions). Orange bars, insertions or deletions within repeat blocks, at block width. Bent arrow, 5S transcription start (2,122 bp). n = 51,219 units from 541 haplotypes of 299 individuals. (B) Unit resemblance within and between arrays. Fraction of units with a matching partner in each comparison set. Each set is size-matched to one haplotype (mean of 5 draws). n = 45,736, 40,797, 45,092 and 45,736 interior units in group order from 541 haplotypes of 299 individuals. (C) Unit difference by separation. Mean pairwise difference (bp) between units of one array. Blue, all units. Orange, most divergent quartile removed. Pale bands, 95% global envelope (extreme rank length) (76) of 2,499 within-array permutations. Central window 0.2-0.8. n = 541 haplotypes of 299 individuals. (D) Exchanged patch length by separation. Maximum-likelihood tract length (78) (marker spacing proxy 105.9 bp). Band, 95% profile-likelihood interval. Dashed, floor (191.9 bp) from pairs of unrelated individuals. n = 541 haplotypes of 299 individuals, with 4,000 pairs drawn per separation and 4,000 floor pairs. (E) Variants per unit along the array. Variants per unit against the majority-allele reference, stacked by origin class. Left, units 1-30 from the proximal junction. Right, 30-1 from the distal junction. Middle cell, mean over all other units, from the 484 haplotypes with at least 61 units. Multi-individual shared, shared frequency f ≥ 0.10 (includes the 12 terminal-type diagnostic variants). Within-array multiple, f < 0.10 and ≥ 2 copies per haplotype. Single-unit, f < 0.10 and one copy. Black line, total. Dashed, mean of the 61 positions (6.82). (F) Variant classes along the array. All variants (left) and single-unit variants (right, own y axis), x axis as (E). Gray dashed lines bound the middle cell (not zone boundaries). Stacked, the 11 classes in any zone’s top four. Gray, 12 others pooled. (G) Proximal-ness by depth. Fraction of units above the proximal-ness threshold, in excess of middle units, by depth from the proximal junction. Color, four array-length strata. Black line, zero. (H) Distal-ness by depth. As (G), from the distal junction. (I) Compartment size against array length. Slope, units added to the proximal, central and distal compartments per added unit of array length. Compartments are k-means clusters (k = 12) with positional structure above the null, on position-free features, by criterion A (gray, variants in 0.5-99.5% of units) or B (black, counts in 20 variant characters). Bars, bootstrap 95% intervals. (J) Region sizes across array lengths. Units per haplotype by region in five length bins (mean 59-142 units), relative to the mean (= 1). Proximal-ness and distal-ness, within depth 10 of each junction. Middle > 30, > 10 and > 5, deeper than that from both junctions. Dotted line, proportional. Gray dashed, fixed. Middle > 30 is empty for 55% of haplotypes in the shortest bin. (K) Zoning of the array. Top, boundaries per haplotype from change-point segmentation (penalty 1.0, criterion B of (I)) by kind, with the cohort boundaries (proximal 3|4 and 17|18, distal 10|11). Bottom, resulting zones (proximal 1-3, proximal 4-17, central, distal 1-10) on a mean-length (95-unit) array, proximal left. n = 1,151 boundaries in 541 haplotypes of 299 individuals. (L) CpG bias of C>T along the array. log₂(C>T at CpG / C>T at non-CpG) at the 61 positions of (E), multi-individual shared variants removed (f < 0.10, origin-f denominator 541). Line, zero. (M) Population make-up of array types. Single linkage (Jaccard ≥ 0.90) on carriage of the 40 most variable tokens in the four zones of (K). The 14 clusters of ≥ 5 haplotypes (counts above bars). Five haplotypes lack a population label. n = 295 of 510 haplotypes, from 213 individuals. (N) Population-private variant tokens. Tokens found in only one population over 200 subsamples to the smallest population size. A token counts if ≥ 3 carrier haplotypes are all in one population. Gray points, subsamples. Line, median. 1,283-1,475 testable tokens per subsample. *P* = 0.001 for the AFR median against 1,000 permutations of population labels between individuals, the smallest attainable value. n = 503 of 510 haplotypes, from 276 individuals with a population label. Where a panel gives no other count, n = 541 haplotypes of 299 individuals.

**Figure 3.**
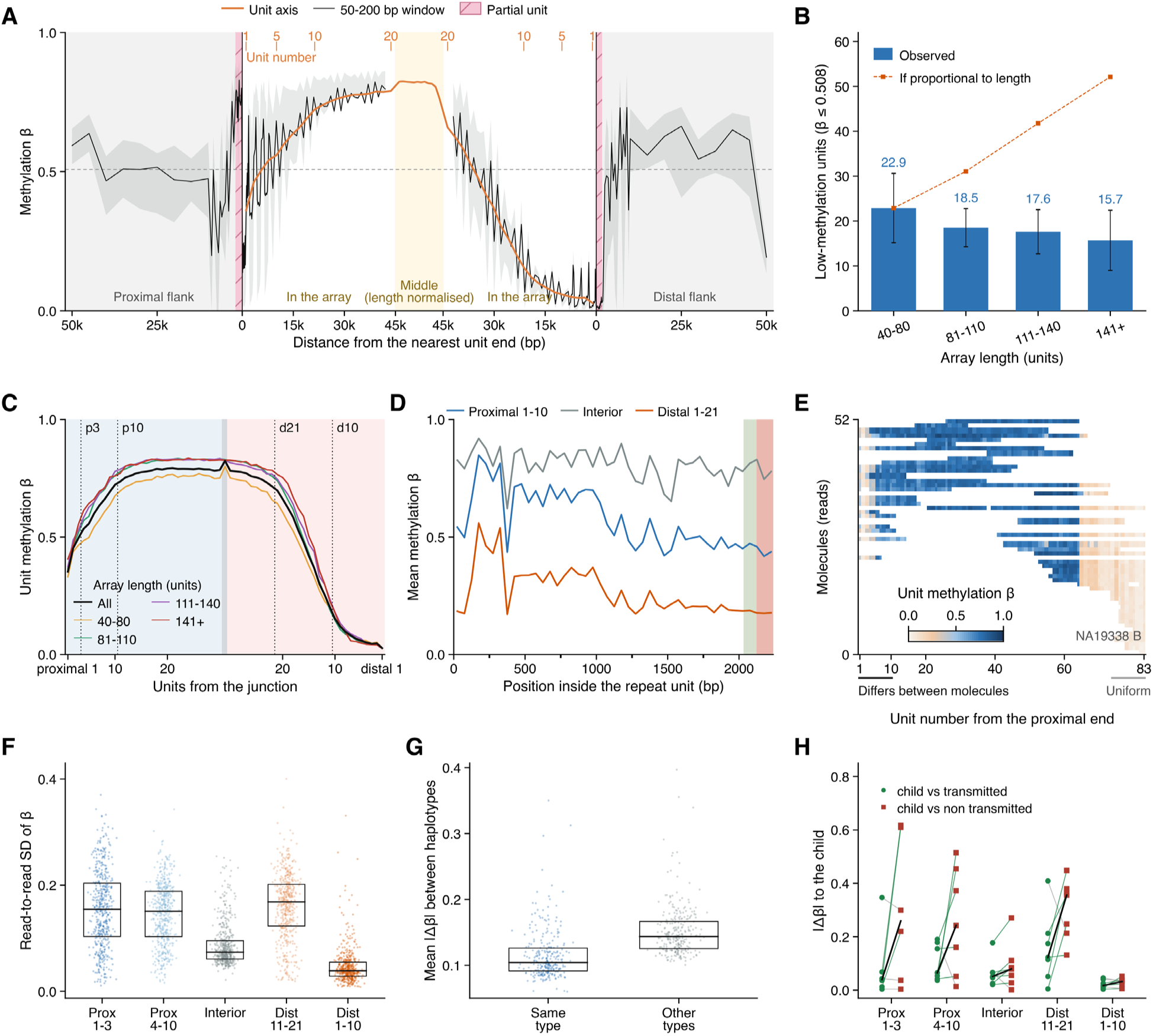
Methylation along the 5S rDNA array. (A) Methylation landscape of the whole array. Mean β along the array, laid out from both junctions, with one value per unit (orange) and a 50 to 200 bp window trace (black, with a quartile band). The x axis is distance from the nearest unit end. Orange ticks number complete units only. The pink hatched bands are the terminal partial units (proximal 1,928 bp, distal 1,635 bp) and the yellow band is the length-normalized middle. The dashed line, unit state cut at β = 0.508. n = 256 windows from 504 haplotypes of 276 individuals and 52 unit positions from 510 haplotypes of 280 individuals. (B) Low-methylation unit count by array length. Low-methylation units (β ≤ 0.508) per haplotype, averaged within four array length strata. Whiskers, standard deviation across haplotypes. The orange dashed line, count expected if these units scaled with array length, anchored on the shortest stratum. The 12 excluded carry fewer than 40 units. n = 180, 189, 92 and 37 haplotypes in the four length strata, 498 of 510. (C) Mean unit methylation by unit number. Unit mean β against unit number counted from each junction, for all haplotypes (black) and the same four length strata as in (B). The middle is units more than 25 units from both ends, and 610 of 45,544 (1.3%) fall on both sides. Four dotted vertical lines, the five zone boundaries, drawn at proximal 3, proximal 10, distal 21 and distal 10 on the same unit numbering as the x axis. n = 45,544 units from 510 haplotypes of 280 individuals. (D) Methylation by position inside one repeat unit. The 2,240 bp unit, oriented in the direction of 5S transcription, in 45 bins of 50 bp, each bin the sum of methylated calls over the sum of calls, for proximal 1-10, the interior and distal 1-21. Bands, the 5′ flank in green and the 5S gene body in red. n = 4,947 proximal, 30,314 interior and 10,283 distal units. (E) Single molecule methylation map of one haplotype. Every molecule of NA19338 haplotype B as a row. The staircase edge is coverage. n = 52 molecules over 83 units. (F) Read to read methylation variability by zone. One point per haplotype in each of five zones, each point the read-to-read standard deviation of β in that zone. Line, median. Box, interquartile range. Zone means 0.154, 0.146, 0.087, 0.160 and 0.047. n = 494, 500, 510, 503 and 496 haplotypes. (G) Methylation distance within and between array types. One point per haplotype, its mean |Δβ| over the five zones to its own array type of Figure 2M against the other types. Line and box as in (F). The within-type distance is the smaller of the two in 245 of 281 haplotypes, median of the per-haplotype difference −0.0375, paired Wilcoxon *P* = 5.8 × 10⁻³⁴. n = 281 haplotypes of 206 individuals in 14 array types. (H) Distance between the child and each parental haplotype in trios. Mean |Δβ| of the child against the transmitted (green circle) and the non-transmitted (red square) haplotype, one line per pair within a zone. Pooled over zones, 7 of 7 pairs are closer on the transmitted haplotype, median |Δβ| 0.072 against 0.165, within-pair median change −0.0950, *P* = 0.0156, which is the floor a two-sided Wilcoxon can reach with 7 pairs. Zone by zone no zone reaches 0.05, lowest BH *q* 0.195. n = 34 pair-zone comparisons from 7 child-parent pairs of 5 trios, HPRC and GIAB.

Interestingly, when the array was longer the low-methylation stretches at the two ends did not grow in proportion to it. Low-methylation units per haplotype stayed relatively steady as array length increased and in fact fell slightly (Figure 3B), and single molecule maps of three haplotypes of different length show the same thing (Figure S3D). A unit was taken as low in methylation below a unit β of 0.508 and high above it, the trough of the per-unit methylation distribution seen in Figure 4. On average, and independently of array length, there is therefore a pattern of change in methylation at the proximal and at the distal end, the length of the low-methylation stretch of an individual haplotype departing from it only a little, and the array divides into five zones by methylation, proximal 1-3, proximal 4-10, distal 11-21, distal 1-10 and the high-methylation interior that remains (Figure 3C), with the zone boundaries given in Methods. Methylation inside the unit is also shown for each part (Figure 3D).

**Figure 4.**
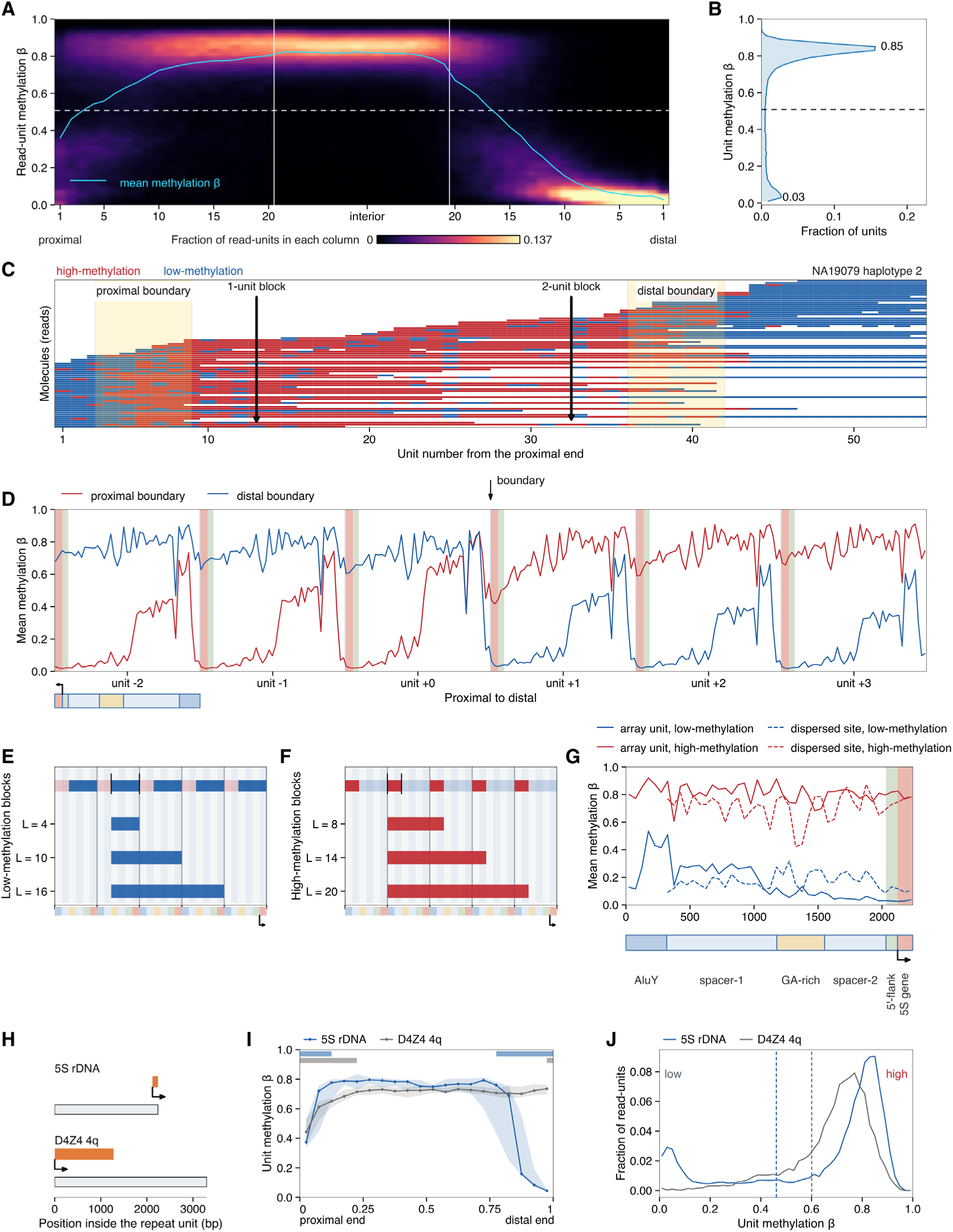
Methylation states of the 5S rDNA repeat unit. (A) Read-unit methylation along the array. Color is the density, the share of a column’s read-units in each β bin, from 0 to 0.137 on the color bar below the panel. Proximal units 1 to 20, the interior and distal units 20 to 1 lie on one axis. n = 733,288 read-units from 510 haplotypes of 280 individuals. (B) Distribution of unit methylation. Modes at 0.03 and 0.85. The dashed line, valley between them, β = 0.508, the cut that assigns units to the low- or high-methylation state (Methods). n = 47,467 units. (C) Unit states along single molecules. Reads of one haplotype (NA19079 haplotype B), one row each, with the proximal and distal boundary zones shaded and a 1-unit and a 2-unit block marked. n = 75 reads. (D) Mean methylation across a boundary. Six consecutive units in 50 bp bins, red across the proximal and blue across the distal boundary. Within each unit the axis runs from the 5S gene to AluY, as the color bar and the bent arrow show, and pink and green bands, the 5S gene and its 5′ flank. n = 3,590 proximal and 6,094 distal boundary windows. (E) Ends of low-methylation blocks. Blocks of 4, 10 and 16 sub-unit elements. n = 27,658 blocks. (F) Ends of high-methylation blocks. Blocks of 8, 14 and 20 elements. n = 6,928 blocks. In (E) and (F) the top row is the chain of the two states, and each bar is a length family of blocks that contain the 5S gene element, placed at its most common start element. Single-element blocks are not drawn. (G) Within-unit methylation of array units and dispersed 5S loci. Dispersed loci are from the 29 sites of Figure S5. Solid lines, array units. Dashed lines, dispersed loci. Blue, the low- and red the high-methylation state. The dispersed side is not drawn over AluY, which almost no dispersed locus covers. n = 867,958 read-units from 580 haplotypes of 301 individuals (array) and 534,880 of the 535,371 loci of Figure S5 (dispersed). (H) Transcribed parts of the 5S rDNA and D4Z4 4q repeat units. Gray bar, repeat unit. Orange bar, transcribed part. Bent arrow, transcription start and direction. (I) Methylation along the whole array in 5S rDNA and D4Z4 4q. Relative position from the proximal end (0) to the distal end (1). Points are the median over array segments of the segment mean in 20 equal bins, and the band is the interquartile range. Bars along the top mark each array’s transition zones, from each end to the first bin whose median reaches 95% of the interior median. n = 13 segments from 13 haplotypes of 7 individuals (5S rDNA) and 12 segments from 11 haplotypes of 7 individuals (D4Z4 4q), the 1kGP set (9 individuals of the 1kGP ONT release). (J) Unit methylation distribution in 5S rDNA and D4Z4 4q. For 5S rDNA (Ashman’s *D* = 3.00) the dashed line, likelihood-ratio cut between the two fitted components, β = 0.4611. D4Z4 4q has no valley to cut at, so its dashed line is a rank split that leaves the 5S share of units (21.9%) below it. n = 25,644 (5S rDNA) and 7,966 (D4Z4 4q) units, the 1kGP set.

The molecules of one haplotype were then read one at a time. Of note, the methylation state was not fixed between reads from the same haplotype, which may come from different cells or from sister chromatids (Figure 3E). Measured over the whole cohort (Figure 3F), the read-to-read standard deviation of β within a haplotype was higher in the three zones in which methylation changes steeply with position, proximal 1-3, proximal 4-10 and distal 11-21, than in the two flat zones, the interior and distal 1-10.

Moreover, comparing the lymphoblastoid cell line with peripheral blood mononuclear cells of the same donor, and normal with tumor, showed that a haplotype of the same individual can carry a different methylation pattern in a different cell state (Figure S3E, F). Gene-body β within high-methylation units fell by a within-pair median of 0.018 in 8 of 10 blood and cell line pairs (*P* = 0.037) and by 0.141 in 6 of 7 normal and tumor pairs (*P* = 0.047), six of those seven normals being cell lines and one patient tissue. The gene body carried the change while the unit β of the whole array moved little (Figure S3G, H).

Finally, arrays that share an array type also share a methylation pattern. Using the 14 array types of Figure 2M, a haplotype stood closer in methylation to the haplotypes of its own type than to the others (Figure 3G). Likewise, when the same haplotype passed from a parent to a child the methylation difference was usually smaller (*P* = 0.0156) than the difference to the haplotype that was not transmitted, although in some comparisons it was larger (Figure 3H). This also shows that methylation here is affected by the array’s own features but is not determined by it alone.

Taken together, the 5S rDNA array carries a methylation pattern that separates it from the flank and divides its inside into zones, and that division is, like the division by sequence variants, relatively insensitive to array length on average, so that the low-methylation stretches at the proximal and the distal end stay about the same size on average as array length changes. Moreover, beyond position within the array, which sets the overall methylation framework on average, several layers act together to form the methylation pattern of a unit in an individual array. These layers include the sequence features of the array itself, differences between individuals, differences between cell types, and, even within one tissue, differences in the state of individual cells, so that regulation can differ from cell to cell.

### Bimodal unit methylation of 5S rDNA

The next question was how the methylation pattern along the 5S rDNA array is distributed at the level of the repeat unit. Unexpectedly, unit methylation in the 5S rDNA array was bimodal, with units accumulating at a low-methylation and a high-methylation mode (Figure 4A, B). The distribution of unit β has its modes at 0.03 and 0.85, and the minimum between them lies at β = 0.508 (Figure 4B). Units were therefore divided into the low- and the high-methylation state at β = 0.508. This feature of the unit sets the boundary between the two states used throughout this work, including Figure 3. The same two-mode shape was observed in all three datasets (HPRC, 1kGP and GIAB), despite differences in submission and chemistry (Figure S4A). The methylation differences along the array seen in Figure 3 and Figure S3 therefore do not arise from a gradual change in the methylation of individual units, but from a proportion of low- and high-methylation units that differs from position to position.

When the reads of one haplotype were laid out unit by unit, units of both states occurred within the same molecule, and the regions in which the state changed within reads were marked as the proximal and distal boundaries (Figure 4C). Aligned on these boundaries and averaged at base resolution, methylation changed from one state to the other mostly within one unit (Figure 4D). On single molecules the change was sharper, taking the form of a step of tens to hundreds of base pairs rather than a ramp across the unit (Figure S4B). When the two states were fitted separately for each sub-unit element, the high-methylation state was similar in every element, whereas the low-methylation state became progressively deeper from AluY toward the 5S gene (Figure S4C). Within a single DNA molecule, units thus step between two states across a boundary.

Along the reads there were also isolated blocks of one or two units surrounded by the other state (Figure 4C). Most units were in one state over all six elements (Figure S4D), and the spacing between successive starts of blocks of the same state peaked at 6, 12, 18 and 24 elements, that is, at multiples of one unit (Figure S4E). Low- and high-methylation blocks began and ended at fixed elements and interlocked, each sharing both of its ends with its neighbor (Figure 4E, F). Methylation therefore does not change independently in each element but changes with the positions of the boundaries fixed within the unit.

The next question was whether these features, namely the bimodality of units, the element-specific methylation within the unit and the change along the array, arise from the 5S sequence or from the repetitive structure itself. To this end the array was first compared with 5S-like loci dispersed outside it and with other tandem arrays. Besides the 5S rDNA array, the human genome contains many 5S-like sequences scattered elsewhere (27). The dispersed loci that carry at least three elements of the 5S rDNA repeat unit (29 sites) were examined, and most of those sites (27 of 29) carry spacer-1 through the 5S gene, that is, the whole unit except AluY (Figure S5A). With the single exception of chr20:30.35 Mb, these loci were called single-mode (Figure S5B). At chr20:30.35 Mb, within the same individuals, molecules in the high-methylation state also carried more highly methylated flanking DNA on the same read than molecules in the low-methylation state, by 0.125 in flank beta in 192 of the 195 individuals that carry both states and with no shuffle of 10,000 reaching the observed value, suggesting that the bimodality of this locus reflects two methylation states of the surrounding region rather than of the 5S-like copy itself. The dispersed loci also differed from the array in their within-unit profile. In array units the low-methylation state fell close to zero from the GA-rich segment to the 5′ flank, whereas at dispersed loci it remained higher over the same elements. The high-methylation state was similar in both (Figure 4G). Therefore, the methylation pattern of the 5S array cannot be explained by sequence alone.

5S rDNA was next compared with five other tandem arrays, D4Z4 4q, D4Z4 10q, DXZ4, RNU2 and SST1 (Figure 4H-J and Figure S6). In lymphoblastoid cell lines these arrays are expected to differ in transcription. RNU2 is an actively transcribed array of Pol II-driven U2 snRNA genes (91,92), DXZ4 is transcribed from both strands, with longer antisense transcripts from the inactive X (93), D4Z4 is expected to be largely silent heterochromatin (94), and full-length DUX4 is not expressed in most somatic tissues (95), and SST1 is methylated in normal tissue (96). After annotating the repeat unit of each array (Figure 4H and Figure S6A, F, L, Q), none of the other arrays separated into two classes at the unit level (Figure 4J and Figure S6C, N, S). DXZ4 appeared bimodal in women (Figure S6H), but this reflected a mixture of the active and the inactive X chromosome (Figure S6G, I). Because arrays without a valley cannot be split by a β cut, their units were divided for the within-unit comparison by a rank split matched to the 5S shares of low- and high-methylation units, 21.9% and 78.1% (Figure S6D, O, T), and for DXZ4 the within-unit profile is drawn by X chromosome instead (Figure S6J). Methylation along these arrays was broadly uniform, except that D4Z4 4q and D4Z4 10q were lower at their proximal ends (Figure 4I and Figure S6B, M, R, with read-by-unit maps in Figure S6E, K, P, U). Of these arrays, D4Z4 4q showed the largest methylation difference along the array and is known to carry a proximal methylation gradient (88), so it was compared with 5S rDNA in detail. In both arrays methylation was low at the proximal end and rose toward the interior (Figure 4I, with read-by-unit maps in Figure S4F, G). The unit distribution, however, was bimodal in 5S rDNA and not in D4Z4 4q (Figure 4J). Aligned on the crossing and averaged at base resolution, 5S methylation moved between its two fitted state levels in one step at the boundary (Figure S4H, I), whereas in D4Z4 4q the whole unit rose gradually, without a jump between two separated levels (Figure S4J). Averaged within the unit after the units were split into two classes, the 5S low-methylation class was deepest over the 5S gene and its 5′ flank, a shape absent from D4Z4 4q (Figure S4K, L). When the total rise of each molecule was decomposed by step size, 5S rose largely through steps of 0.3 or more (56% of the rise) and D4Z4 4q largely through smaller ones (Figure S4M). The two arrays thus share a proximal gradient, but in 5S rDNA it is produced by a changing proportion of units that switch between two states, and in D4Z4 4q by a continuous rise in the methylation of the whole unit. Taken together, 5S rDNA methylation has three features, the bimodality of units, the element-specific methylation within the unit and the change along the array, and these features are not a simple consequence of sequence or of repetitive structure but a property of the 5S rDNA array itself.

### Methylation state of the 5S rDNA repeat unit tracks chromatin accessibility and nucleosome occupancy

CpG methylation of 5S rDNA units had thus been shown to vary systematically with position within the array, and that feature is tied to a trait specific to the array, one that distinguishes it from dispersed 5S loci. The next question was whether the methylation differences mapped across units track chromatin architecture, that is, whether they relate to the functional state of the units in the array. To connect this variation to the role of the units, Fiber-seq was used, which reports chromatin state along individual long molecules.

Fiber-seq uses a non-sequence-specific adenine methyltransferase, the m6A-MTase Hia5, to deposit m6A on adenines not occluded by protein (Figure 5A). DNA wrapped in nucleosomes is largely devoid of m6A, and within accessible stretches the positions occupied by transcription factors remain as short unmethylated gaps (62). Reading each molecule at full length therefore yields alternating methylated and unmethylated intervals, which resolve into two layers. Unmethylated blocks of nucleosomal length are assigned to nucleosomes, and the intervening methylated intervals, the MTase-sensitive patches (MSPs), to accessible DNA (63). Smaller gaps nested within MSPs are read as footprints of non-nucleosomal proteins such as transcription factors (62,65). Unlike population-averaging assays, Fiber-seq resolves nucleosome placement and accessible DNA on the same molecule, and can therefore distinguish whether molecules at one position exist in a mixture of states.

**Figure 5.**
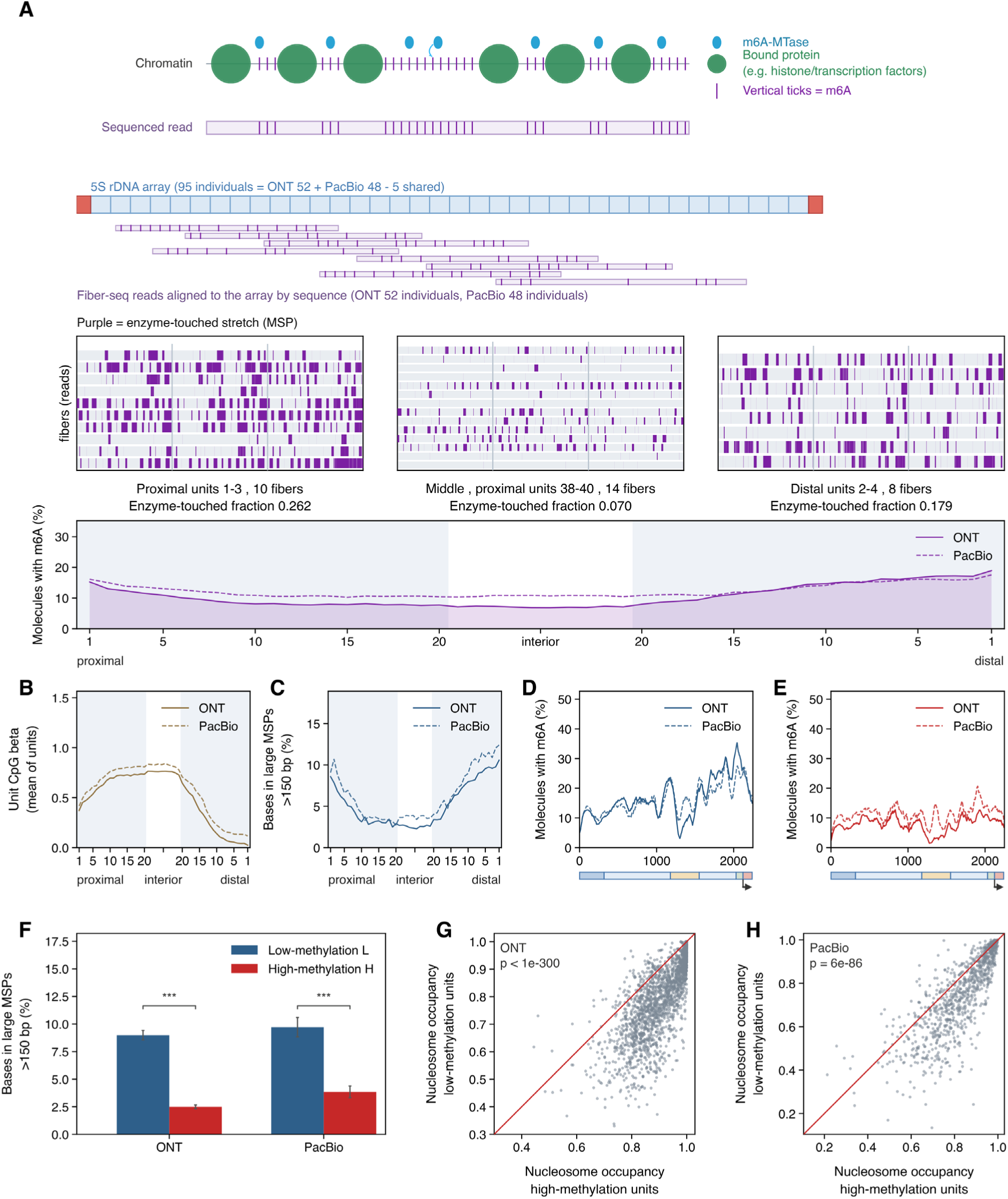
Chromatin architecture and CpG methylation across the 5S rDNA array in single Fiber-seq reads. In (A) to (E) ONT is the solid line (52 individuals) and PacBio the dashed line (48 individuals). (A) Principle of Fiber-seq and m6A per array position. Top, a schematic of chromatin and of the sequenced read, in which the blue ellipses are the m6A-MTase Hia5, the green circles are bound protein (histones and transcription factors) and the purple vertical ticks are m6A. Middle, the array reference with Fiber-seq reads aligned to it by sequence (purple), and below it three windows of one array (HG01891 haplotype B, 113 units) drawn as fiber stacks, the proximal end, the middle and the distal end, with purple marking enzyme-touched stretches (MSPs). Bottom, the fraction of molecules carrying m6A at each array position, taken as occurrences at a base divided by the coverage at that base. PacBio m6A was called with the provider’s jasmine model, 95 distinct individuals in all with 5 of them shared between the platforms. In (A) to (C) the shaded margins are the 20 absolute unit positions at either end and the white center holds the interior as relative positions. (B) CpG methylation per array position. The unit CpG β averaged over units. All units are pooled. (C) Density of large MTase-sensitive patches per array position. Bases covered by MSPs longer than 150 bp divided by called bases, with MSPs called by FiberHMM. (D, E) Within-unit m6A per unit position. Low-methylation units in (D, blue) and high-methylation units in (E, red), classified at a unit CpG β of 0.508. The colored band under each panel marks the elements of the unit, which run from left to right as the AluY element, spacer-1, the GA-rich region, spacer-2, the 5′ flank and the 5S gene. A bent arrow under the band marks the transcription start site of the 5S gene and the direction of transcription, and the band carries color only with no element names. (F) Large-MSP density in low-methylation and high-methylation units, paired within single molecules. Low-methylation (L, blue) against high-methylation (H, red) units, ONT and PacBio separately, with MSPs called by FiberHMM. Both bars are means over the same molecules, those traversing units of both classes. Error bars, 95% confidence intervals. Asterisks, a two-sided Wilcoxon signed-rank test over the paired per-molecule means (***, *P* < 0.001, ONT 2,111 molecules from 52 individuals, PacBio 1,010 molecules from 48 individuals). (G, H) Per-molecule nucleosome occupancy in low-methylation against high-methylation units, ONT in (G) and PacBio in (H). Each point is one molecule, averaged over the units of each class that it traverses, and only molecules traversing units of both classes are included. The red diagonal marks equality. *P* is a two-sided Wilcoxon signed-rank test on the per-molecule differences (ONT, n = 2,111 molecules from 52 individuals, 1,964 of 2,111 in the same direction, *P* < 1 × 10⁻¹⁰; PacBio, n = 1,010 molecules from 48 individuals, 758 of 1,008, *P* < 1 × 10⁻¹⁰).

Fiber-seq data were analyzed for those HPRC individuals with a 5S rDNA array reference, 52 individuals for ONT and 48 for PacBio and 95 distinct individuals in total, and the reads of each molecule were placed onto that array reference. Both ends of the array, proximal and distal, carried more enzyme-accessible DNA than the array interior. The same pattern appeared in the per-position m6A profile and held for ONT and PacBio alike (Figure 5A).

CpG methylation was low at both ends and high in the interior, as in Figure 3 and also in the 52 ONT and 48 PacBio individuals analyzed here (Figure 5B). The density of accessible patches, the fraction of called bases covered by large MSPs, that is MSPs longer than a nucleosome, was likewise elevated at the same two ends (Figure 5C). The large-MSP threshold was taken from Dubocanin et al. (68). The pattern was not explained by differences in read coverage (Figure S7A).

Units were then partitioned by their methylation level, at a unit CpG β of 0.508. Within-unit m6A was highest over the 5′-flanking region and the 5S gene in low-methylation units, and was uniformly low across all elements in high-methylation units (Figure 5D and E). Accessible-patch density was also higher in low-methylation units, and this difference was maintained on both platforms when the two classes were compared in a paired fashion within the same molecule (Figure 5F). Resolved into five windows of the unit, the difference was largest over the 5′ upstream window and the 5S gene and smallest over the AluY element, without a monotonic ordering in between, and was significant in 10 of 10 window-by-platform combinations (Figure S7C).

Nucleosome occupancy gave an answer in the same direction. Occupancy, for each molecule and unit, is the number of bases assigned to nucleosomes divided by the total called bases of that unit, that is the nucleosome and MSP bases, with the full criteria given in the Supplementary Methods. High-methylation units showed higher occupancy than low-methylation units (Figure 5G and H). To exclude a per-molecule difference in labeling efficiency, the comparison was repeated within single molecules, where enzyme exposure is shared and both classes carry the same repeat sequence and hence the same adenine density. The difference did not collapse (Figure S7B), and the paired comparison uses the 44.6% of ONT molecules and the 21.4% of PacBio molecules that span units of both classes. The sign was unchanged in all fourteen combinations of seven ways of splitting the two classes and two platforms, and in all fourteen combinations of seven large-MSP thresholds spanning 100-300 bp and two platforms, with *P* ≤ 1 × 10⁻²³ throughout. The seven splits are the single cut-off used above and six two-sided gaps, which leave units of intermediate methylation out, the narrowest keeping units at or below 0.45 against those at or above 0.60 and the widest at or below 0.20 against those at or above 0.85. Both sweeps were computed from the same FiberHMM calls as the figures (Supplementary Methods).

Taken together, low-methylation 5S rDNA units carry more accessible DNA and fewer nucleosomes than high-methylation units, and both differences persist when the two classes are compared within the same molecule.

### Footprint distribution differs with the methylation state of the 5S rDNA repeat unit

The next question was whether occupancy at the transcription-factor binding sites differs between low- and high-methylation units. The binding architecture of a canonical type 1 Pol III promoter (97), the class to which the 5S gene belongs, is drawn in Figure 6A. The Pol III machinery that occupies promoters of this class in human cells has been mapped genome wide (98). To call footprints from Fiber-seq data, FiberHMM was used (63,65), which partitions each molecule, from its m6A pattern and its segment lengths, into nucleosomes, methyltransferase-sensitive patches (MSPs) and the shorter protected intervals nested within them (Figure 6B, Figure S8A). Because whether a footprint edge is resolvable at all depends on the local adenine density, no position was nominated in advance and every offset and every size class across the unit was allowed to compete (Figure 6B, Figure S8C, F, and Methods).

**Figure 6.**
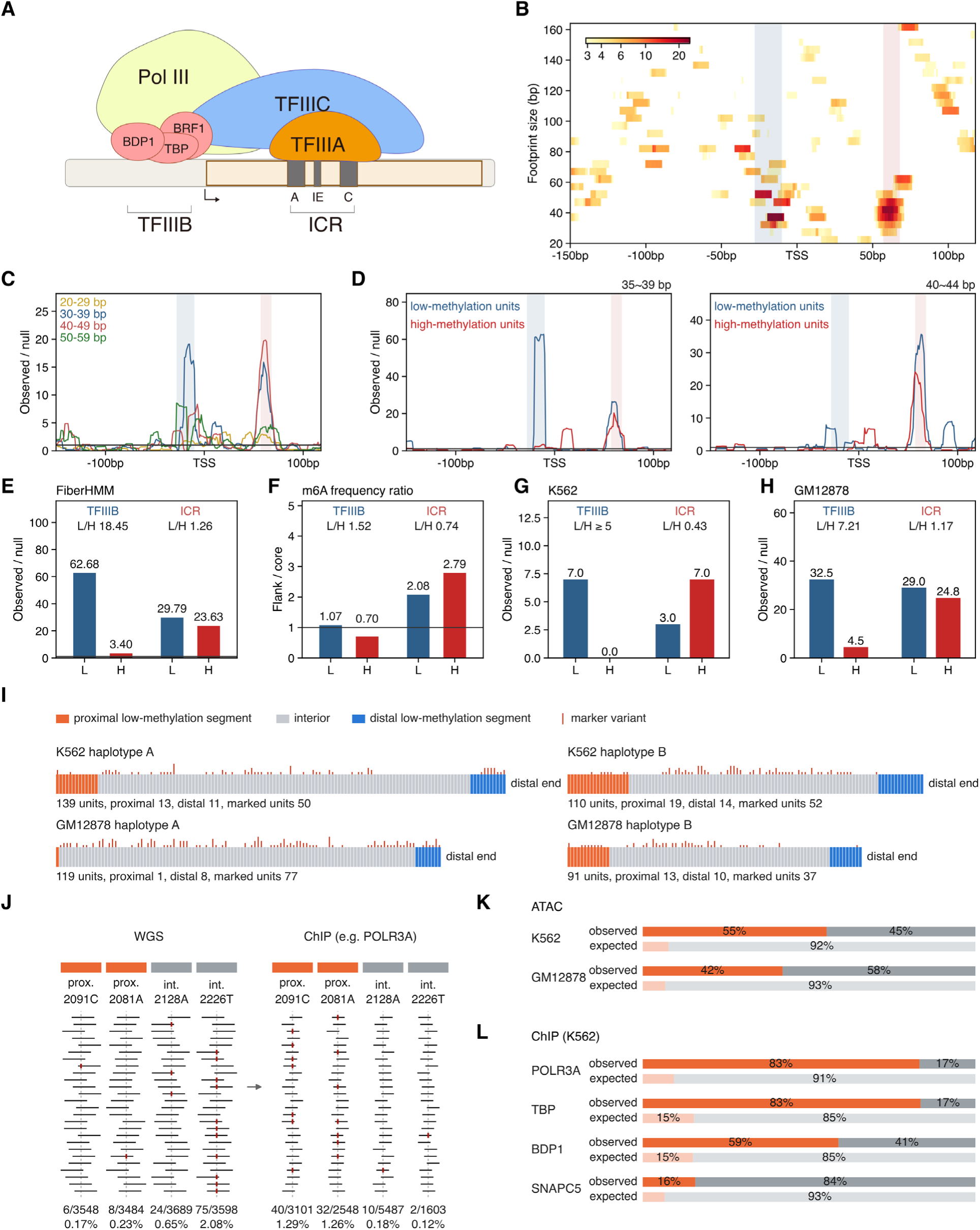
Methylation-related transcription-factor footprints across the 5S rDNA repeat unit. (A) Promoter architecture of the unit. Schematic of a type 1 Pol III promoter, showing the TFIIIB-binding site upstream of the gene and, within the gene, the internal control region with its A box and C box. (B) Grid scan of footprint position against footprint size. HPRC PacBio Fiber-seq, 30,330 molecules. FiberHMM m6A-protected intervals, every position against every size (20-164 bp, 5 bp bins) over −150 to +118. Only the window was fixed in advance. Color gives fold excess over a circular-shift null within each unit. Cells below the permutation threshold are gray. Bands mark TFIIIB −28 to −10 (blue) and the internal control region +57 to +68 (red). (C) Footprint density by size class. The same scan in 10 bp classes, pooled regardless of methylation, observed over null. The solid horizontal line, null level of 1, at which observed equals expected. The 30-39 bp class peaks at −15 (19.1-fold) and the 40-49 bp class at +62 (19.8-fold). These are the 10 bp pooling, while the finer 5 bp bins used in (D) peak at −11 and +58. All 89 cells clearing the threshold are ranked in Figure S8C. (D) Position profiles of the two leading size bins, by methylation state. The 5 bp bins ranking first and second in that methylation-blind scan, 35-39 bp (left) and 40-44 bp (right), for low- and high-methylation units, 7,451 low against 22,879 high of the 30,330 molecules, split at unit CpG β 0.508. The solid horizontal line is again the null level of 1, and the dashed verticals mark −11 and +58. (E) Footprint frequency at the two positions, by methylation state. Low over high methylation, TFIIIB 62.68/3.40 = 18.45 (173/16 footprints) and internal control region 29.79/23.63 = 1.26 (49/82). The solid horizontal line, null level of 1. Ratios here and in (F), (G) and (H) are computed from the unrounded values, not from the rounded occupancies printed beside them. (F) Caller-free flank-to-core m6A ratio at the two positions. Flank m6A frequency (−35 to −27 and +22 to +27) over core m6A frequency (−7 to +1), unsmoothed and with no null. TFIIIB 1.07/0.70 = 1.52 and internal control region 2.08/2.79 = 0.74. The solid horizontal line is at 1, which here is the level at which flank equals core and is not a null level. The axis here is flank over core rather than observed over null, so the panel is read within itself and not against (E). (G) The same two positions in K562. Observed over null, with the solid horizontal line at 1. Here the internal control region is read at +62, the position the cell-line scan gives, not the +58 of (A)-(F). TFIIIB 7.0/0.0, written L/H ≥ 5 because the high side holds no footprint, and internal control region 3.0/7.0 = 0.43. Units 2,334 (1,106/1,228), labeled by gene-body β 0.5 over the last 119 bp with ≥ 5 CpG, so the denominator is units and not the molecules of (A)-(F). K562 is presence/absence (15/1,106 against 0/1,228), bounded from below. (H) The same two positions in GM12878. Observed over null, with the solid horizontal line at 1, the internal control region again read at +62 and the unit label as in (G). TFIIIB 32.5/4.5 = 7.21 and internal control region 29.0/24.8 = 1.17. Units 5,501 (5,428 labeled, 1,572/3,856). Both positions clear the null in both lines (grid-wise *P* ≤ 0.0010). (I) Zones and zone-exclusive markers on the four cell-line haplotypes. Zone boundaries are change points in gene-body methylation (Figure S9A), not a fixed β cut. Markers number 70 in K562 (7 proximal, 59 interior, 4 distal) and 78 in GM12878 (3, 68, 7). K562 haplotype B is drawn in its 110-unit marker frame, and its methylation frame has 98 units. (J) Marker read fractions in WGS and ChIP-seq, with POLR3A as the example and marker bases counted at base quality ≥ 20. Reads over two proximal and two interior markers are drawn schematically with the marker allele in red, and the measured marker reads over depth are given under each column. The expected count of a zone is the depth at each of its markers times the WGS marker fraction, summed over the markers. (K) Zone composition of ATAC-seq marker reads in both cell lines. Observed shares of proximal and interior marker reads are set against the shares expected from WGS, with marker bases counted without a base-quality filter. Reads aligned to the array number 16,041 in K562 and 14,701 in GM12878, of which 15,278 and 14,686 pass the alignment gate. (L) Zone composition of ChIP-seq marker reads in K562, for the Pol III machinery and the negative control SNAPC5. Shares are drawn as in (K), with marker bases counted at base quality ≥ 20. Reads aligned to the array number 61,226 for POLR3A, 158,494 for TBP, 21,059 for BDP1 and 6,035 for SNAPC5, of which 18,877, 102,695, 3,491 and 5,931 pass the alignment gate.

Protection was concentrated in the 0-60 bp size range at two positions, around the TFIIIB-binding site upstream of the gene and the internal control region (ICR) within it (99), where TFIIIA binds (Figure 6B, Figure S8A, B). Footprints of comparable length and site have been reported for TFIIIA and TFIIIB on the human 5S rRNA gene or on 5S and other Pol III genes in other species (100,101). These two ranked first and second among the 89 grid peaks that cleared the grid-wide 0.05 criterion (Figure S8C). In untreated control material the TFIIIB window was labeled more heavily than the unit average (1.95) and the ICR slightly less (0.77), so the TFIIIB signal is not a property of the sequence itself, although the ICR may be in part (Figure S8D).

Subdividing by footprint length separated the two positions by size as well (Figure 6C). The 5 bp classes carrying the strongest enrichment were 35-39 bp at TFIIIB and 40-44 bp at the ICR, peaking at −11 and +58 respectively. Both classes were fixed in the methylation-blind scan and only then split into low- and high-methylation units (Figure 6D).

Splitting at a whole-unit CpG β of 0.508, TFIIIB occupancy was 18.45-fold higher in low-methylation units (62.68 against 3.40), whereas the ICR was only 1.26-fold higher (29.79 against 23.63) (Figure 6E). The asymmetry is supported statistically. The TFIIIB direction held in all 45 individuals carrying a footprint at that site (one-sided sign test *P* = 2.8 × 10⁻¹⁴), whereas the ICR held in 27 of 46 individuals (*P* = 0.15, not significant).

A caller-independent readout, the ratio of raw m6A density in the flanking windows to that in the central window, recovered both positions as footprints, each exceeding its own permutation null (both *P* < 5 × 10⁻⁴) (Figure 6F, Figure S8E). On this axis too TFIIIB was higher in low-methylation units (1.07 against 0.70), whereas at the ICR the ratio was 2.08 in low-methylation units and 2.79 in high-methylation units. The stronger TFIIIB footprint in low-methylation units is therefore consistent across both axes, while the ICR is not. The two strata were not tested against each other on this axis. The ratio has no fixed baseline in the absence of protection, because the flanking and central windows differ in adenine content, and protection can lift it above that baseline by at most a factor of 1/(1 - f), where f is the fraction of protected molecules. At the TFIIIB site the measured value sits at that ceiling, whereas at the internal control region it exceeds it about twofold and is therefore not explained by protection of individual molecules. Its magnitude accordingly reflects window composition and occupancy rather than the strength of protection.

The contrast is not accounted for by intrinsic under-labeling, by nucleosome occlusion or by caller sequence context (Figure S8G). Nor does it depend on the choice of label. Moreover, restricting the denominator to molecules carrying no nucleosome call over the position under test left the TFIIIB footprint in place (Figure S8I). Over the site windows (−28 to −10 for TFIIIB and +57 to +68 for the ICR) this conditional occupancy gave a low-to-high ratio of 12.3 at TFIIIB, far above the 1.0 that pure occlusion predicts, while the ICR fell to 1.05 (Figure S8J). The direction was the same on the caller-independent axis, where a molecule is excluded if a nucleosome call overlaps −35 to +27 about the center (Figure S8K). Lastly, selecting units whose gene body alone is low-methylated, rather than the whole unit, gave the same split, 14.1-fold at TFIIIB against 1.2-fold at the ICR (Figure S8H).

Taken together, the scan recovered footprints at the two positions that the promoter architecture predicts, the TFIIIB-binding site and the internal control region where TFIIIA binds. Of the two, the footprint at the TFIIIB-binding site was consistently more occupied in low-methylation units, on the caller-based and on the caller-independent axis alike.

### Cell-line ATAC-seq and ChIP-seq place accessibility and Pol III machinery binding in the low-methylation zone

Low- and high-methylation units thus differ in both chromatin accessibility and footprint pattern. The next question was whether the same conclusion is supported in parallel by other assays, such as ATAC-seq and ChIP-seq, using two cell lines in which multiple such assays have been performed by ENCODE (102), K562 and GM12878. To this end the HPRC results were first tested for reproduction in the two cell lines and then, in those where they were, this analysis examined the methylation-related zone distribution of the ATAC-seq and ChIP-seq signal.

Each array was first divided by the methylation of the cell line itself. In K562, however, the difference between end and interior units was weak for whole-unit methylation but was retained for gene-body methylation (Supplementary Table S2), so gene-body methylation was used. On haplotype-resolved array contigs assembled from the long reads of each cell line, the units from each array end to a change point in gene-body β formed the proximal and distal zones, and the remainder the interior zone (Figure S9A).

In both cell lines the end zones were largely low in methylation and the interior high, by gene-body methylation in K562 and also by whole-unit methylation in GM12878, except at the proximal end of GM12878 haplotype A, a single unit whose values are unstable (Figure S9B and Supplementary Table S3). The end zones also had more m6A and more gene-body bases covered by large MSPs (Figure S9B), and footprints were enriched at the TFIIIB position and the internal control region (Figure S9C, D, F, G). The TFIIIB position was more protected in low-methylation units in both cell lines, both in FiberHMM footprints and in the caller-free m6A flank-to-core ratio (Figure 6G, H and Figure S9D, E, G, H). The internal control region showed a small or line-dependent difference in footprints (Figure 6G, H), and by the flank-to-core ratio it was slightly higher in high-methylation units, but at this position that ratio does not report the strength of protection (Figure S9E, H). Fiber-seq in the two cell lines therefore goes in the same direction as in HPRC (Figure 6E, F).

The relationship of methylation to chromatin accessibility and to Pol III machinery binding was next confirmed in parallel using ATAC-seq and ChIP-seq from these cell lines. ATAC-seq and ChIP-seq fragments are short, however, and cannot be assigned to one unit among units that are nearly identical. The relationship was instead tested indirectly, by comparing the proportion of fragments that carry zone-exclusive variants between whole-genome sequencing (WGS) and ATAC-seq or ChIP-seq (Figure 6I, J).

Figure 2 showed that variants within the array have a characteristic distribution across its regions. The question was therefore whether there are region-exclusive variants that link a fragment to a position within the arrays of these two cell lines. The within-unit variants of each zone were first identified in the array of each cell line (Figure 6I), a variant found only in units of one zone being an exclusive marker of that zone (proximal, interior or distal), excluding the AluY and GA-rich segments and hypervariable units and pooling the two haplotypes. Because the assemblies may be incomplete, contigs were rebuilt from long reads carrying the marker allele alone and each marker was tested for lying in its original zone (Figure S10A-C). None of the significant contigs lay in another zone (0 of 115 in K562 and 0 of 153 in GM12878), and the few away from their original site (7 and 1) lay elsewhere in the same zone or could not be placed. As a further check, independent of array position, K562 reads were cut into units and the markers related to gene-body methylation (Figure S10D). Of the units carrying a proximal marker 89% of 64 were low in methylation, as were 85% of the 149 carrying a distal marker, against 25% of the 887 units carrying an interior marker. In these cell lines the markers therefore represent the state of low- and high-methylation units, independently of unit alignment. This yielded proximal- and interior-exclusive markers, 70 in K562 (7 proximal, 59 interior and 4 distal) and 78 in GM12878 (3, 68 and 7), listed in Supplementary Table S4. The distal zone also carries markers, but it was not scored in either cell line. When WGS reads were cut to fragment length and passed through the same alignment gate, the distal zone alone was inflated, 1.5- to 1.6-fold in GM12878 and 1.3- to 1.4-fold in K562, and in K562 75-100% of the distal reads came from a single marker (365T) that is also carried by a dispersed 5S locus on chr12. Only the proximal and interior zones were therefore compared.

The distribution of ATAC-seq and of ChIP-seq within the unit was then examined for the targets of 5S rRNA transcription (Figure S10E, F). Only fragments explained by the 5S rDNA array alone, that is, fragments that match the array better than the dispersed 5S loci or the rest of CHM13, were retained. The reads expected at a marker are the read depth at that position multiplied by the marker-allele fraction in WGS, that is, what the assay would place there if it sampled the array copies as WGS does. Each zone was then scored as its observed marker reads, summed over its markers, divided by that expectation summed over the same markers (Figure 6J). Scored in this way, ATAC-seq gave more proximal-marker reads than expected in both GM12878 and K562 (Figure 6K).

The same scoring was next applied to ChIP-seq. It could be scored in K562 alone, because the read depth covering the proximal marker positions in the GM12878 control track is too low. That depth multiplied by the marker fraction in WGS comes to fewer than one read, so the track cannot serve as the denominator of the interaction. Likewise BRF1 and GTF3C2 in K562 were not scored, because their read depth covering the proximal marker positions is too low, leaving more than about a 5% chance that no proximal read would be observed and a zone ratio would not exist (Figure S10G).

Each ChIP target was compared with the control track and with the negative control SNAPC5 (a subunit of SNAPc, which acts at snRNA-type promoters but not at the 5S promoter (97)). For each target the proximal-to-interior ratio is given, which divides the proximal zone ratio by the interior one, both against WGS. In K562 it was 47.8-fold for POLR3A (the largest subunit of Pol III), 27.9-fold for TBP and 8.1-fold for BDP1 (both subunits of TFIIIB), against 2.7-fold for SNAPC5 (Figure 6L). To exclude the uneven recovery that chromatin accessibility, fragmentation and mappability give a ChIP library, the proximal-to-interior ratio of a target was then divided by that of the control track, giving the interaction, so that a value near 1 means no more proximal excess than in the control track. The interaction was 20.9 [4.6-112.7] for POLR3A and 12.1 [3.6-45.5] for TBP, intervals that lie above 1, whereas SNAPC5 gave 1.2 [0.1-3.9], no more than the control track, and BDP1 gave 3.5 [0.8-21.5], an interval that includes 1. The zone ratios and interactions against the control track are summarized in Figure S10H (ATAC-seq) and S10I (ChIP-seq). In K562 the proximal ChIP reads were concentrated on the top three markers, but with these three removed the proximal zone ratio remained at 8.36 for TBP and 5.34 for POLR3A. Together, these results reproduce the Fiber-seq result in both cell lines and show a difference in zone distribution, in ATAC-seq in both cell lines and in ChIP-seq in K562, that supports it in parallel.

### The methylation pattern of the 5S rDNA array in different species

Finally, whether the methylation pattern found in the human array is evolutionarily conserved was asked of the 5S rDNA arrays of the eight species for which long reads are available, namely five apes, zebra finch, zebrafish and *Arabidopsis thaliana*. Figure 7A shows every chromosome that carries a 5S array, one block per species and drawn to scale, with the centromere marked. Across the eight species, 65 arrays on 21 chromosomes were cataloged and 49 of them analyzed. The arrays drawn in gray were cataloged but not analyzed, and each of them carries three to five copies, which is the range of that gray set, since an array entered the analysis when a repeat unit frame could be built for it. Each chromosome is drawn with the lower assembly coordinate at the left, and the horizontal axis of the landscape panel in Figure 7B follows the same rule. The five ape chromosome 1 have used here store the q terminus as base 1, the opposite of the human reference.

**Figure 7.**
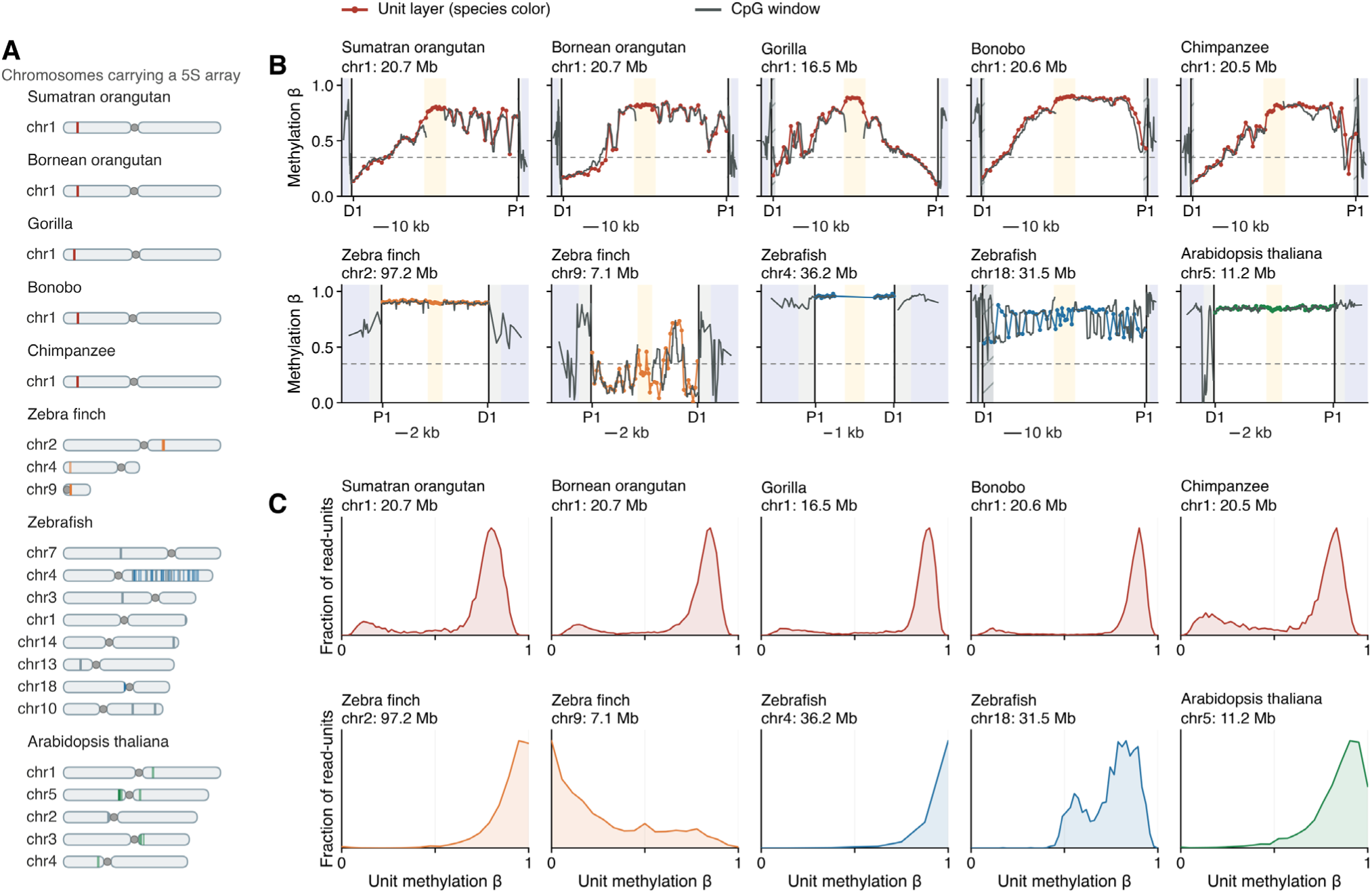
Methylation pattern in the 5S rDNA arrays of eight species. (A) Chromosomal position of every cataloged array. One block per species, drawn to scale, showing the chromosomes that carry a 5S rDNA array. Each chromosome is drawn in the orientation of its species reference with position 1 at the left. Vertical marks are arrays, all of the same length and thickness. Full color marks the ten arrays shown in B and C, pale color the other analyzed arrays, and gray the cataloged arrays for which no repeat unit frame could be built from the assembly, all of them carrying three to five copies. 8 species, 21 chromosomes, 65 arrays, 49 of them analyzed. (B) Methylation landscape of ten representative arrays and of their flanks. The colored line, unit layer, one point per unit slot, and the dark line is a CpG window that also covers the flanks. Both lines are the same statistic, the call-weighted mean β, and inside the array the window is one repeat unit wide so the two coincide. Shading marks the flank, the flank beyond 2 kb with the axis compressed, and the array interior between the twenty end units with the axis compressed. Hatching marks the partial unit at each end of the array, which is excluded from the analysis. The dashed line is β = 0.35. (C) Per-unit methylation of the ten arrays. Distribution of per-unit β for the same ten arrays of (B) in the same order. Bins are the values β can take for that array. Each array is counted on its own and the ten are not pooled. Complete units per array run from 1,583 to 93,323, and the mean number of CpG calls per unit from 8.3 to 190.5. CG context only for the animal species.

The arrangement itself differs by an order of magnitude. Each of the five apes carries exactly one array, in every case on chromosome 1, whereas zebra finch carries four arrays on three chromosomes per haplotype, *A. thaliana* ten arrays on five chromosomes of which nine were analyzed, and zebrafish 46 arrays on eight chromosomes of which 31 were analyzed. The repeat unit differs between species and between arrays. Figure S11A draws the repeat unit of the representative array of each species on a single base-pair scale. Unit length runs from 177 bp to 2,312 bp, while the gene itself is 118-119 bp in every species and lies at the end of the unit, so the entire difference is in the spacer.

The methylation landscape within the array (Figure 7B) places the apes with the human array and separates every other species from it. In all five apes methylation of the array is depleted at both ends and high in the interior, and in all but the gorilla the wider band of low methylation lies at the distal end. In the gorilla the wider band lies at the proximal end. That exception is because the gorilla is the only one of the five whose 5S genes are transcribed toward the distal end. Among the other species, zebra finch and zebrafish vary between the arrays of one genome, while *A. thaliana* is uniformly high in CG and almost flat along the array. Figure S12 draws, in the same way, the 39 analyzed arrays that are not in Figure 7B, and none of those 39 shows the edge depletion seen in the apes.

At the level of the unit (Figure 7C) the distribution of per-unit β separates the species sharply. All five apes are bimodal, as the human array is, with fitted components at β 0.195-0.252 and 0.787-0.881 and a low-methylation fraction of 8.8-26.2%. In zebra finch the arrays separate by level, three of them lying above mean β 0.898 and the chromosome 9 array at 0.274. In zebrafish 1 of 31 arrays is bimodal, the array on chromosome 18, against 6 unimodal, 23 not testable and 1 reserved. Not testable is a resolution limit and not a negative result, because a unit in those arrays carries only 7-8 CpGs, so β can take too few distinct values for the test to stand. None of the 39 arrays outside the main figure is bimodal, all of them being unimodal and high, with median array β 0.910 in zebra finch, 0.956 in zebrafish and 0.859 in *A. thaliana* (Figure S11B). For *A. thaliana* the CHG and CHH contexts were also measured and are given separately in Figure S11C, and the β used in Figure 7 and in panels A and B of Figure S11 is the CG value alone. Methylation along the repeat unit is given for all 49 arrays in Figure S13. A gene window methylated below the rest of the unit, the feature that organizes the human unit, appears with appreciable amplitude in only two places, the five apes and zebrafish chromosome 18.

Taken together, the bimodal methylation of units and the position-dependent difference within the array found at the human locus are conserved within the apes, differ from species to species outside them, and in zebra finch and zebrafish differ between the arrays of one genome, at the level of the unit and at the level of the array alike.

## Discussion

This work describes the sequence and methylation features of the human 5S rDNA array and shows that units are bimodal in methylation, a property related to chromatin architecture.

For sequence variation, earlier studies that did not use long reads could not separate individual units and mostly reported how often a variant occurs among the copies of a person (32,33). A recent study using long-read assemblies mapped variants in the array (32), but how variation is organized by position from the proximal to the distal end of a whole array was not appreciated. These features show characteristic patterns of variation within the array (Figure 2).

Three components make up this landscape. They are concerted evolution (Figure 2B-D) (103), a region-specific tendency of variant accumulation (Figure 2E-H and L), and a class of specialized units (Figure 2E and Figure S2I). Remarkably, the array kept relatively constant numbers of proximal-type and distal-type units as it became longer or shorter (Figure 2I-K). Why variants accumulate differently between regions, and how the proximal and distal zones keep their size regardless of array length, remain to be studied. The regional differences may be linked to the other properties that change along the array. For example, the distinct CpG bias of C>T substitutions on the distal side (Figure 2L) may reflect the low methylation of the distal end (Figure 3A), since methylated cytosine at CpG readily deaminates to thymine (104,105).

Like the zone by sequence variants, methylation along the array also marks out a proximal and a distal feature, and on average that zone’s length changes little with the total unit number of the array. The basic rule that forms the framework of the methylation pattern is a positional one, low at the ends and high in the interior. However, there appear to be more factors that contribute to the methylation of individual units on the array, which together make up the methylation pattern of an individual haplotype. By comparing different haplotypes (Figure 3G), transmission in trios (Figure 3H), different cell types (Figure S3E, F) and molecules within the same sample (Figure 3E, F), array sequence traits and cellular state prove able to affect the methylation pattern beyond position within the locus.

Among the states of the cell, unit methylation being related to chromatin architecture, those that can bear on 5S rDNA activity are candidates that can influence methylation. One is the cell cycle and proliferation of the individual cells, which regulate 5S rRNA gene transcription directly (3) and the activity of RNA polymerases I and III (12), and are therefore expected to influence 5S rRNA expression. Another is a change of the cell state. For example, Epstein-Barr virus immortalization induces active polymerase III transcription (10), and in cancer, polymerase III transcription is frequently elevated (8,106,107) and inhibiting polymerase III attenuates colorectal tumorigenesis (9). Long-read analysis of defined cell states, for example a comparison after cell cycle arrest, or comparing the methylation of arrays in samples before and after immortalization or transformation can test whether those states of the cell influence 5S rDNA methylation.

The bimodality of unit methylation is one of the methylation features that set the 5S rDNA array apart from the dispersed 5S units and from the other repeat arrays examined here (Figure 4G and J). The two unit states defined by that bimodality, low-methylation and high-methylation, are associated respectively with relatively high chromatin accessibility and relatively frequent transcription-factor-sized footprints, and with relatively low accessibility and relatively infrequent footprints of that kind. Just as the low-methylation stretches are concentrated at the proximal and distal ends of the array, accessibility was likewise higher at those two ends than in the array interior (Figure 5A to 5C). The same held at the level of the elements within a unit, where the within-unit methylation and m6A profiles of high-methylation units differed comparatively little from element to element (Figure 4G and Figure 5E), whereas in low-methylation units both the methylation and the m6A values differed considerably more between elements (Figure 4G and Figure 5D). Methylation and chromatin accessibility in the 5S rDNA array are therefore associated at the level of the array, of the unit, and of the elements within a unit.

The Fiber-seq scan places footprints at two positions in the Pol III binding apparatus, the TFIIIB-binding site and the internal control region, and only the TFIIIB footprint tracks methylation state reproducibly. Earlier work on 5S rDNA and on Pol III transcription is compatible with this pattern. Most directly, TFIIIA, which binds the internal control region, was shown to bind the actively transcribed somatic-type and the largely inactive oocyte-type 5S genes of *Xenopus laevis* alike (108) and with equal affinity (109). Separate regulation of TFIIIB and TFIIIC, which TFIIIA recruits in 5S rDNA, has also been reported under inhibition of Pol III transcription. Prolonged repression by nutrient deprivation in yeast reduced TFIIIB occupancy at tRNA genes while TFIIIC occupancy rose (110), and Pol III transcription inhibition through p53 also reduced TFIIIB occupancy at human tRNA genes while TFIIIC occupancy was largely unchanged (111). In yeast, TFIIIB was found on only 23 to 47% of the 150 to 200 chromosomal 5S gene copies in vivo, and a smaller fraction, 8 to 17%, carried an open complex (112), so partial and copy-to-copy TFIIIB occupancy at 5S genes is not itself new, and the measurement here resolves it to single units along one array. The asymmetry found here therefore suggests that a similar separate regulation of TFIIIB and TFIIIA, which recruits TFIIIC, may act between units along the array. What differs between low-methylation and high-methylation units is then the binding of TFIIIB, which binds stably upstream of the start site and directly recruits Pol III (113,114), rather than that of TFIIIA, which acts as an assembly factor (97,115).

The ATAC-seq and ChIP-seq results from the cell lines went in the same direction as the Fiber-seq accessibility and footprint pattern (Figure 6K, L). SNAPC5 (2.7-fold) and the control track (2.3-fold) also had a higher zone ratio in the proximal than in the interior zone, although by less than the Pol III machinery (Figure S10I), which may be due to a difference in accessibility between the zones, because sonicated chromatin is itself enriched at accessible regions (116). One caveat is that the long-read assemblies and the short-read data did not come from cells of a single laboratory, so although the pattern was observed at multiple markers and separately for different proteins, their variant distributions may differ (117). Observing the relationship between transcription and methylation directly will require methods such as DiMeLo-seq (118) to confirm this finding, which would also resolve the fact that the proteins at the footprints called from the Fiber-seq data cannot be named. In addition, unlike at 45S rDNA, where DNA methylation follows NoRC-mediated histone modification and impairs preinitiation complex formation (14), it remains to be elucidated at 5S rDNA whether methylation or transcription comes first and which influences the other.

The two features reported here, the bimodality of unit methylation and the dependence of methylation on position within the unit, are largely shared among the apes. Outside the apes they differ from species to species and between the arrays of one species, so these methylation features are not a universal property of a 5S array. Among those species the result for the zebrafish arrays is interesting. Of the zebrafish arrays analyzed, the chromosome 18 array is the only one called bimodal (Figure 7C), while the chromosome 4 arrays are unimodal and high, at a median array beta of 0.956. Figure S13 places the difference in the gene. In the chromosome 18 array beta falls to 0.372 over the gene window against 0.753 over the whole unit, whereas in the chromosome 4 arrays the gene window sits 0.007 to 0.019 above the unit mean, and the same gene-centered separation distinguishes the two unit states in the human array. Zebrafish 5S rRNA genes fall into a maternal-type and a somatic-type family that are expressed at different developmental stages (119), and that literature assigns the two families to chromosomes, placing the somatic-type genes on chromosome 18 and the maternal-type genes on chromosome 4 (119,120). The reads used here come from adult whole brain at 9 months post-fertilization, which is somatic tissue at a stage in which the somatic-type family is the one in use. Unit methylation and gene expression may therefore be related in zebrafish as well.

Recovering the full 5S rDNA array of every haplotype and reading it unit by unit reaches what a shorter view of the locus cannot. It shows the properties of whole units, and the way those properties are organized by position from the proximal to the distal end. On that view the array carries a methylation pattern of its own, that within a single array the units are bimodal in methylation rather than graded, falling into low-methylation and high-methylation classes, and that which class a unit belongs to goes with chromatin architecture. By showing that the methylation of a human 5S rDNA unit goes with its chromatin accessibility and with footprint occupancy, this links the control of expression within the human 5S rDNA array to epigenetic change. It also opens the way to see the regulatory landscape of 5S rDNA, that is, which units in an array carry the marks of active transcription and which do not, and where along the array they sit.

## Data availability, Supplementary data, Funding

## Data availability

Cell-line short-read data are ENCODE experiments (102,121,122); the file accessions used for K562 (TBP, POLR3A, GTF3C2, BDP1, BRF1, SNAPC5, the control tracks, ATAC-seq and whole-genome sequence) and for GM12878 (TBP, POLR3G, the control tracks and ATAC-seq) are listed in Supplementary Table S5. The GM12878 whole-genome denominator is 1000 Genomes run ERR3239334. The human 5S rDNA arrays and reads come from public long-read resources. From the HPRC come assemblies (release 2 index assemblies_release2_v1.0.index.csv) and ONT reads (BioProject PRJNA701308; s3://human-pangenomics/working/HPRC/; release 2 data (34) and, for 100 individuals, data not yet part of a release (35), of which 37 are also deposited as nanopore runs under that BioProject while the remaining 63 carry no INSDC accession; four of the 100 (HG01204, HG01516, HG02470 and HG03919) contributed reads to the assemblies only and not to the read placement), together with the year-1 HG002 assembly (23). The other resources are the 1kGP-LRSC ONT release (s3://1000g-ont/PROCESSED_DATA/, including its GIAB HG003 assembly; described by its providers as possibly unpublished) (38) with the publication its providers request (24), Oxford Nanopore reads of the GIAB samples (s3://ont-open-data/giab_2023.05/; distributed under the Creative Commons Attribution-NonCommercial 4.0 license, CC BY-NC 4.0) (36,37), GIAB HG001 PacBio HiFi (s3://giab/data/NA12878/PacBio_CCS_15kb_20kb_chemistry2/) (36,70) and Oxford Nanopore reads of GM12878 (ENA ERR17530411, BioProject PRJEB114757, which is GIAB HG001 read data downsampled to 30x and haplotagged by that submission rather than an independent GM12878 deposit) and of HG001 from the 2025.01 open-data release (s3://ont-open-data/giab_2025.01/analysis/wf-human-variation/sup/HG001/, flow cells PAW79146 and PAW81754) (37), and Oxford Nanopore reads of K562, which carries a prime-editing construct (BioProject PRJNA1371648). Population labels are the 1000 Genomes phase 3 panel (123) and the 3,202-sample pedigree (124). The matched normal and tumor pairs are three releases. Five cell line pairs are from the CASTLE release (125), BioProject PRJNA1086849, in which every one of the ten samples used here carries Oxford Nanopore whole-genome runs. The submission metadata of that study lists Oxford Nanopore among its platforms. Because several of these samples carry more than one such run, the study is named rather than a run. The COLO829 pair is the 2024 Oxford Nanopore open-data release (126), s3://ont-open-data/colo829_2024.03/, whose basecalls hold the tumor and the matched lymphoblastoid line in separate folders. The HG008 pair is from the GIAB somatic release, ftp-trace.ncbi.nlm.nih.gov/ReferenceSamples/giab/data_somatic/ HG008/Liss_lab/, which holds four Oxford Nanopore directories, of which Northeastern_ONT-std_20240422 holds the normal pancreas sample. The normal and the tumor both carry CpG methylation calls. HPRC Fiber-seq, the largest Fiber-seq dataset used here, is released through the Human Pangenome Reference Consortium on AWS Open Data rather than through an INSDC archive: the raw submissions read here are under s3://human-pangenomics/submissions/, in the submissions named UCSC_HPRC_ONT_YEAR1_FIBERSEQ (ONT, under the prefix 5ECA1D3E-1C37-44B7-BA20-576417F786F0--) and WUSTL_HPRC_PacBio_FiberSeq (PacBio, under the prefix 4f00ec2d-4533-40e3-b0ca-624ab5a559d9--), one folder per sample under its HPRC sample identifier. The raw submitted BAMs were read rather than the processed track, because the processed hprc-epigenome modbed files carry m6A only and the analyses here need 5mC on the same molecule. K562 Fiber-seq is ENA project PRJEB121523, run ERR17608832. GM12878 Fiber-seq is the FIRE release (127). That work distributes its processed Fiber-seq results on AWS, one folder per sample, and the file read here is the processed s3://stergachis-manuscript-data/2024/Vollger_et_al/FIRE/GM12878/fire/ GM12878.fire.bam, which is 326,249,442,731 bytes and carries its own index. The raw reads of that release are deposited as BioProject PRJNA1233341 (run SRR32614294, sample SAMN47268219, cell line GM12878, PacBio Sequel II, 25,311,792 reads) and were not read here. That file is a merge: read names in it carry eleven distinct PacBio movie identifiers, found in all fourteen intervals sampled, and the single run identifier in its header is one of the eleven. Four of the eleven match deposited runs - SRR17281093 and SRR17281094 (GEO GSE186009, BioProject PRJNA771694) and SRR24282070 and SRR24282071 (BioProject PRJNA956114) - and for the remaining seven no separate run was found in the records searched, all eleven being present in the merged deposit SRR32614294; the two GSE186009 movies supply 15.7% of 6,203 sampled reads and 21.2% of the reads sampled from chr1:228.6-228.8M. An earlier version of this manuscript said that GSE186009 is a different deposit and not the file read here. That was wrong: it is part of it. The file holds 24,771,424 mapped reads and none unmapped, which does not reconcile by subtraction with the 25,311,792 spots of SRR32614294, because ft fire --skip-no-m6a and the per-chromosome view each drop reads.

Cross-species Nanopore reads were obtained from public archives and are listed per run in Supplementary Table S6. Zebrafish is ENA project PRJEB53713 (128), runs ERR12118112-ERR12118117 (six wild-type controls, whole brain, 9 months post-fertilization, female). That project holds both the transcriptome and the methylome of the same study, which is how its key resources table describes it; the data-availability statement of the paper names the accession for the RNA-seq, and the runs used here are the Oxford Nanopore methylome deposited under the same project. The ape read sets were produced for a study of the ape sex chromosomes (129) and the ape reference assemblies for a separate study (130); the two are different studies and both are used here. For the apes and zebra finch: *Pan troglodytes* (mPanTro3) PRJNA976701; *Pan paniscus* (mPanPan1) PRJNA986878; *Gorilla gorilla* (mGorGor1) PRJNA986879; *Pongo pygmaeus* (mPonPyg2) PRJNA976699; *Pongo abelii* (mPonAbe1) PRJNA976700; *Symphalangus syndactylus* (mSymSyn1) PRJNA976702; *Taeniopygia guttata* (bTaeGut7) PRJNA1241433. *Arabidopsis thaliana* (Col-CC) raw signal is ENA PRJEB79853 (131), run ERR17795792.

Six bonobo PromethION flowcells basecalled with Guppy 6.3.7 are not deposited in SRA and were retrieved from GenomeArk; the bonobo value reported in the main text uses the Guppy 6.3.8 flowcells only. Reference assemblies, each read as its GenBank (INSDC) record: *Pan troglodytes* mPanTro3 GCA_028858775.3; *Pan paniscus* mPanPan1 GCA_029289425.3; *Gorilla gorilla* mGorGor1 GCA_029281585.3; *Pongo pygmaeus* mPonPyg2 GCA_028885625.3; *Pongo abelii* mPonAbe1 GCA_028885655.3; *Symphalangus syndactylus* mSymSyn1 GCA_028878055.3; *Taeniopygia guttata* bTaeGut7 GCA_048771995.1 (maternal plus Z, bTaeGut7.mat) and GCA_048772025.1 (paternal autosomes, bTaeGut7.pat); *Danio rerio* GRCz12tu GCA_049306965.1; and *Arabidopsis thaliana* Col-CC GCA_028009825.2. The six ape assemblies and the maternal zebra finch assembly are the current NCBI reference genomes for their species; GRCz12tu and Col-CC are not, the designated references being GRCz13ab (GCA_052040795.2) for *D. rerio* and TAIR10.1 (GCA_000001735.2) for *A. thaliana*. Zebra finch reads were re-basecalled with Dorado 2.1.2 from 9 of the 13 flowcells screened, and *A. thaliana* raw signal was re-basecalled with the same version (models hac@v5.0.0 and hac@v6.0.0). Siamang reads are as released (dorado 0.9.1); the primate read sets were not re-basecalled, and their per-flowcell Guppy versions (6.3.7 and 6.3.8) are recorded in the data catalog of this work for all 45 flowcells. The zebrafish reads are as deposited (R10.4.1, Guppy 6.4.2 per the depositors), realigned from GRCz11 to GRCz12tu. The arrays lie on *A. thaliana* contigs CP116280.1, CP116281.2, CP116282.1, CP116283.2 and CP116284.1 (chromosomes 1 to 5), on zebrafish contigs CM111182.1 (chromosome 4) and CM111196.1 (chromosome 18), on zebra finch maternal contigs CM109790.1, CM109792.1 and CM109798.1 (chromosomes 2, 4 and 9) with paternal counterparts CM109751.1, CM109753.1 and CM109759.1, and, one array per species, on CM054434.2, CM055472.2, CM055446.2, CM054630.2, CM054679.2 and CM054520.2 in the apes, in the order given above.

Upstream processing was identical for all nine species and the versions were read back from the outputs themselves: 5S genes were located with cmsearch (Infernal 1.1.5, September 2023) (132) against RF00001.cm of Rfam 15.0 (133) (family 5S_rRNA; CM length 120, gathering threshold 38.00 bits, checksum 242171328, built with Infernal 1.1.4 on 4 July 2024) with --noali --cut_ga, that is at the default gathering threshold with no parameter changed; reads were aligned with minimap2 2.26-r1175 (48,49) (-y -ax map-ont -- secondary=no), filtered and sorted with samtools 1.19.2 (50) (view -b -F 0×900), and methylation was called with modkit v0.6.4 (134) (--cpg --combine-strands for the animals, per-motif for the plant).

Supplementary data. Supplementary Data are available at NAR Online. Supplementary Table S1 lists the panel definitions and, on a second sheet, the cohort counts, the sequencing chemistries and the software releases, Supplementary Table S5 the ENCODE file accessions and Supplementary Table S6 the cross-species read and assembly accessions, and Supplementary Table S7 gives within-array resemblance, its controls and the methylation of terminal-type units. Supplementary Table S8 gives the per-marker TFIIIC (GTF3C2) counts in K562, which are reported as a table rather than a panel because the proximal estimate rests on seven reads, and Supplementary Table S4 lists the zone-exclusive markers of the two cell lines with their unit numbers in each haplotype. Supplementary Table S9 gives the three nucleosome controls for the cell-line footprint and zone measurements, the TFIIIB contrast restricted to nucleosome-free units (S9A), the nucleosome deficit at the array ends (S9B), and the label dependence, deviance decomposition and by-element accessibility that bound the zone interpretation (S9C). These were supplementary figures in an earlier version and are given as a table here because each is a short list of numbers. The Index sheet of the Source Data file lists, for every panel of every figure, the sheet that holds the values that panel draws, with the number of rows in that sheet and, for the few panels that have no sheet, the reason.

## Acknowledgements

The author acknowledges the Human Pangenome Reference Consortium (BioProject ID: PRJNA730823) and its funder, the National Human Genome Research Institute (NHGRI). The GIAB cell lines/DNA samples HG001-HG004 (GM12878, GM24385, GM24149 and GM24143) behind the Oxford Nanopore open data used here were obtained from the NIGMS Human Genetic Cell Repository at the Coriell Institute for Medical Research. The author thanks the Stergachis laboratory (University of Washington) for the publicly released Fiber-seq data used here, including the GM12878 FIRE release (127). The author thanks the ENCODE Consortium and the ENCODE production laboratories of Michael Snyder (Stanford University), Kevin Struhl (Harvard Medical School) and Sherman Weissman (Yale University) for the ChIP-seq and ATAC-seq data used here. The two K562 whole-genome sequencing experiments used here for a denominator are on the ENCODE portal but are not ENCODE consortium products, and the author thanks the laboratories of Feng Yue (Pennsylvania State University) and Alexander Urban (Stanford University) for them.

## Supplementary Material

### Supplementary figure legends

**Supplementary Figure S1.**
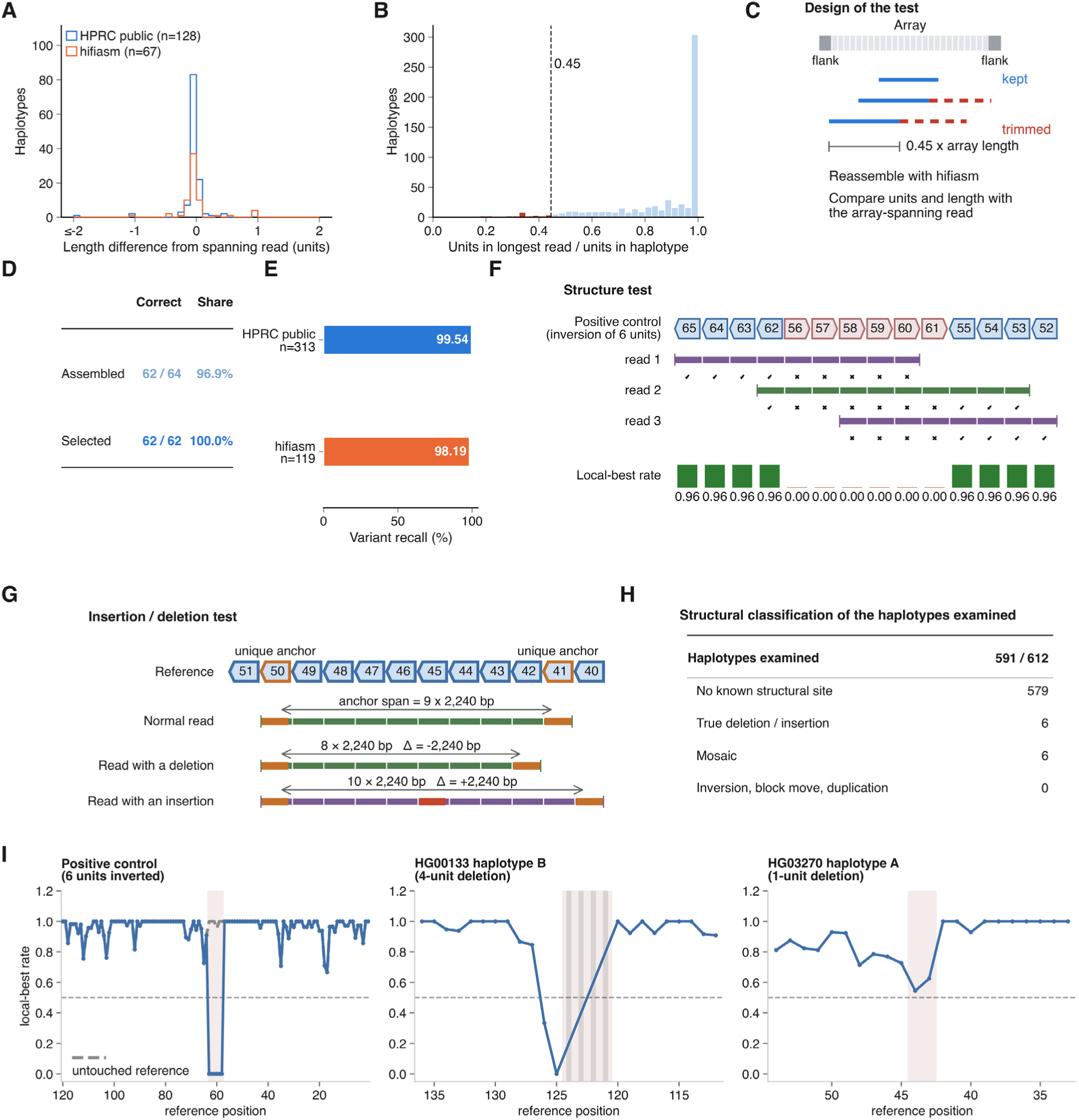
Validation of the array references. (A) Reference unit count against an array-spanning read. Agreement between the unit count of the reference and of an array-spanning ONT read. n = 195 HPRC haplotypes. (B) The longest-read span ratio. For each haplotype the span of its longest array read is divided by the array length, and the threshold drawn is the fifth percentile of that ratio. (C) Design of the trimming and reassembly test. Every read that covers more than 0.45 of the array length is trimmed back to that limit. A read carrying a flank is trimmed at its non-flank end, and a read lying wholly inside the array is trimmed at one of its two ends, chosen at random. The array is then reassembled. (D) Outcome of the trimmed reassembly. The test of panel C run across 32 individuals and 64 haplotypes. (E) Recovery of the variant content of the reference. The reference is compared with a PacBio HiFi read consensus. (F) Design of the structure test. The local-best rate asks, for every placed read unit, whether the reference unit at the position it was placed on is the best match among the reference units within ±5 positions. (G) Design of the insertion and deletion test. The second and independent axis is the base-pair distance between two unique anchor units, measured against the distance the reference predicts. (H) Structural classification of the haplotypes examined. The 591 haplotypes the structure test reached, classified by re-examining read by read the 13 candidate structural sites flagged in 12 of them. (I) Worked examples of the local-best rate. Three examples spanning the range, including an in-silico inversion of six units. Each panel’s denominator is the set of haplotypes that met that panel’s own data requirement, so it differs between panels. Definitions as Figure 1.

**Supplementary Figure S2.**
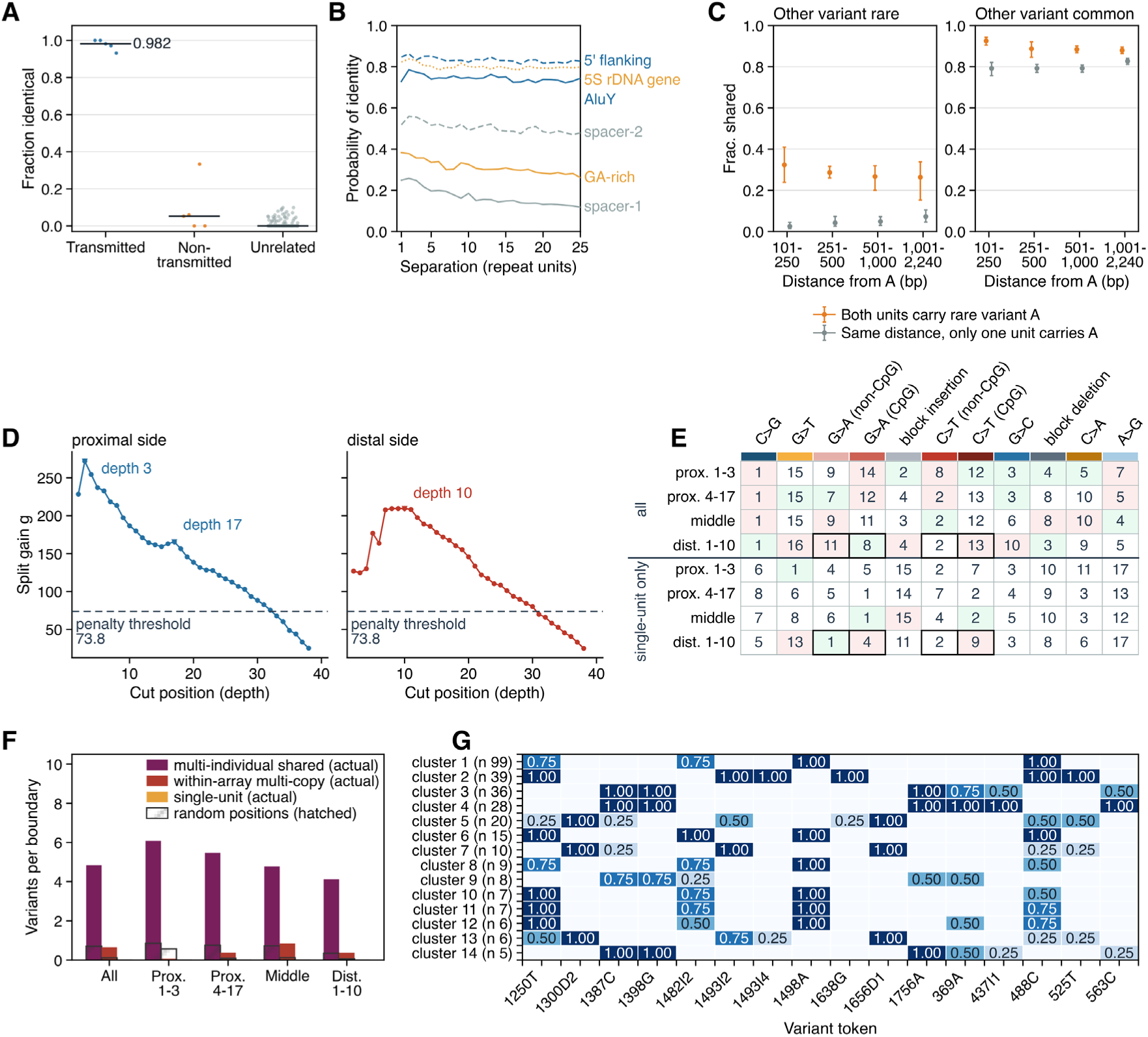
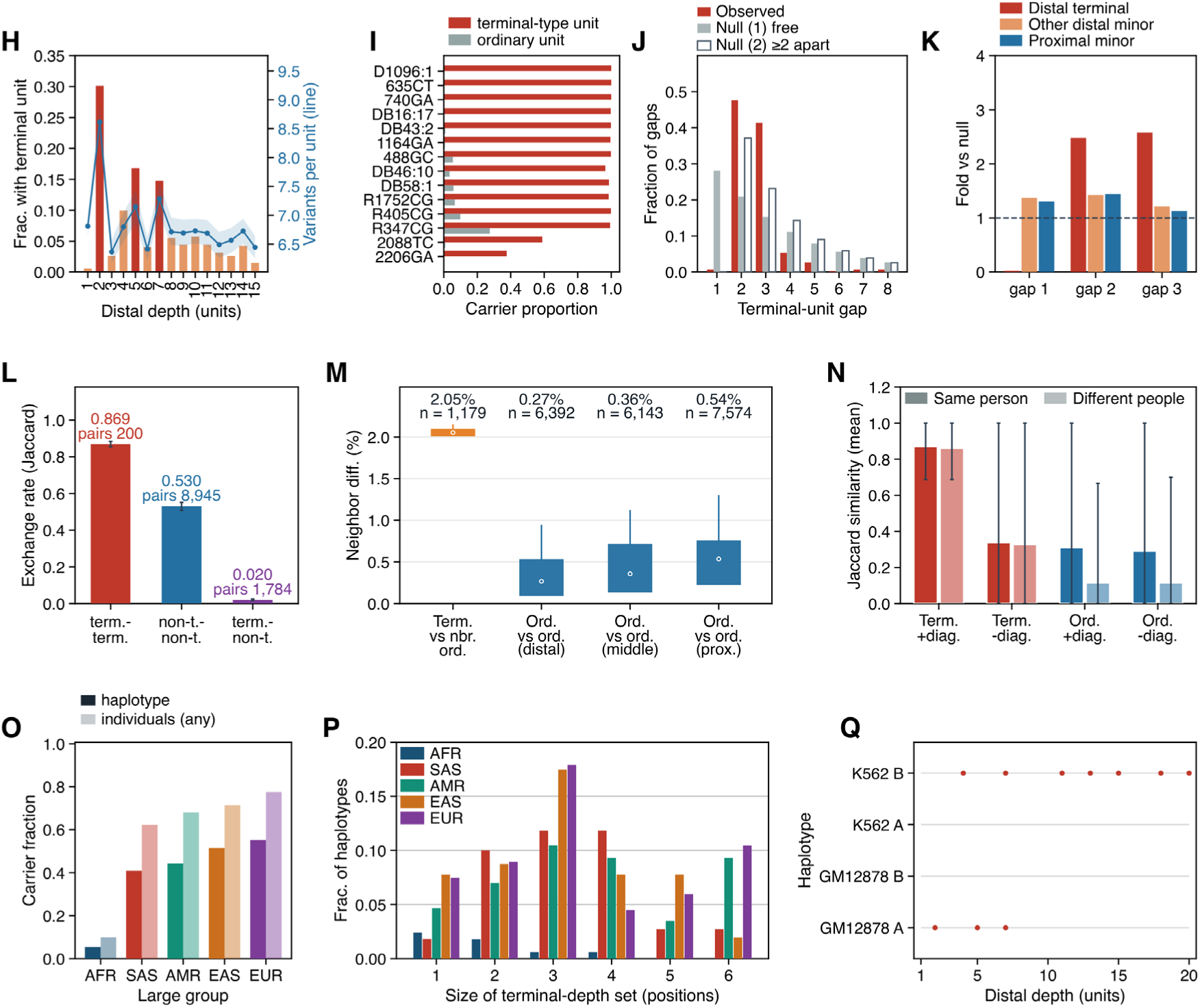
Unit transmission and resemblance, array zoning and types, and the distal terminal-type unit. (A) Trio transmission of units. Fraction of aligned units identical at the matched position, per pair. Transmitted, offspring against the transmitting parent’s haplotype. Non-transmitted, against that parent’s other haplotype. Unrelated, other families. The number printed on the panel is the median of the transmitted pairs. Transmitted pairs were selected at Jaccard ≥ 0.95, hence no *P* value. n = 5 transmitted (447 units), 5 non-transmitted (297) and 150 unrelated pairs (9,730). The pair counts are fewer than the haplotypes they are drawn from, because only pairs passing the Jaccard cut enter: the comparison drew on 12 offspring haplotypes of 6 individuals and 15 parental haplotypes of 8 individuals, in trios. (B) Whole-region identity by separation. Probability that a whole unit region is identical between two units of one array, by separation (1-25). Color and line style mark regions, named at line ends. n = 541 haplotypes of 299 individuals, with 2,000 pairs drawn per separation. (C) Variant co-sharing around a rare variant. Fraction of other variants B shared by two units, by distance from a rare variant A (< 1%) to B. Orange, both units carry A. Gray, one does, at the same separation. Left, B rare (< 1%). Right, B common (≥ 10%). Bars, haplotype-cluster bootstrap 95% intervals. Central window, 0.2-0.8 of the array. Distances ≤ 100 bp not drawn. n = 541 haplotypes of 299 individuals, with 110,974 (orange) and 98,257 (gray) unit pairs. (D) Split gain by cut depth. Proximal end (left) and distal end (right). Line, penalty threshold 73.8 (1.0 × 20 features × ln 40). The distal marker (depth 10) is the analysis boundary. The split gain is flat, with depths 7 to 11 within 99% of the maximum. n = 541 haplotypes of 299 individuals. (E) Variant class ranks by zone. Rank of variant classes by variants per unit in each zone, for all variants (left four columns) and single-unit variants (right four). Bold, top four. Green, z ≥ +5 against 200 within-haplotype unit shuffles. Red, z ≤ −5. Rows, the 11 classes of Figure 2F. n = 1,623, 7,574, 36,612 and 5,410 units in zone order, from 541 haplotypes of 299 individuals. (F) Origin of variants at zone boundaries. Variants distinguishing a zone boundary, by origin class (as Figure 2E), for all boundaries and by the zone in which the boundary falls, proximal 1-3 (69), proximal 4-17 (150), central (713) and distal 1-10 (181). Change-point segmentation by criterion A of Figure 2I. A boundary in two zones goes to the shallower. Hatched, null of random non-boundary positions in the same haplotype. Single-unit variants cannot distinguish a boundary. n = 1,113 boundaries in 541 haplotypes of 299 individuals. (G) Token fingerprints of array types. A token is one variant treated as present or absent in a haplotype, as in Figure 2N. Rows, the 14 clusters of Figure 2M (haplotype count). Columns, the 16 most separating tokens. Value, fraction of the four zones in which the token separates the cluster (within carriage ≥ 0.80, within minus outside ≥ 0.30), so 1.00 means all four zones. n = 295 of 510 haplotypes, from 213 individuals. (H) Terminal-type units by distal depth. Fraction of haplotypes whose unit at each distal depth (1-15) is a terminal-type unit (bars, left axis), with variants per unit at that depth (line, right axis). Terminal-type units are defined in all panels as units carrying at least 10 of the 12 diagnostic variants in (I). n = 541 haplotypes of 299 individuals. (I) Diagnostic variants of the terminal type. Carrier fraction of each variant among terminal-type and ordinary units at distal depths 1-15, for the 14 variants with the largest difference. The 12 with a difference of 0.70 or more are the diagnostic variants, and units carrying at least 10 of them are terminal-type units. Because the type is defined by these same variants, their high carriage among terminal-type units follows in part from that definition. n = 598 terminal-type and 7,517 ordinary units, from 541 haplotypes of 299 individuals. (J) Gaps between terminal-type units. Gap (units) between adjacent terminal-type units of one haplotype, observed and under two nulls, (1) terminal-type units of each haplotype placed freely and (2) placed at least two positions apart. The 3 gaps above 8 are not drawn. n = 414 gaps from 160 of the 541 haplotypes of 299 individuals. (K) Gap frequencies of minority units. Frequency of gaps 1, 2 and 3 between adjacent minority units of the same kind, divided by a label-shuffle null (1 = null), for distal terminal-type, other distal minority and proximal minority units. n = 354, 2,401 and 1,934 pairs from 163, 1,442 and 1,331 runs of adjacent units of the same kind, within 541 haplotypes of 299 individuals. One haplotype can hold more than one run, so the run counts are not haplotype counts. (L) Exchange between terminal-type and neighboring units. Variant Jaccard (exchange rate) of unit pairs within a gap of 2 at distal depths 1-15, by pair type, that is by the classes of the two units making up the pair, over the 19 of 3,869 sites polymorphic within both terminal-type and non-terminal units (at least 5 units each). Numbers, mean and pair count. Bars, haplotype bootstrap 95% intervals. n = 200, 8,945 and 1,784 pairs in bar order from 125, 465 and 184 of the 541 haplotypes of 299 individuals. (M) Sequence difference between adjacent units. Difference (%) between directly adjacent units by pair type. Point, median. Thick bar, interquartile range. Line, to the 90th percentile only; the 10th percentile is not drawn. Numbers, median and pair count. Only 3 pairs of directly adjacent terminal-type units exist, too few to show. n = 1,179, 6,392, 6,143 and 7,574 pairs in bar order from 541 haplotypes of 299 individuals. (N) Terminal-type resemblance within and between people. Mean variant Jaccard of random unit pairs at distal depths 1-15, within and between people, for terminal-type (Term.) and ordinary (Ord.) units, with (+diag.) and without (−diag.) the 12 diagnostic variants. The −diag. comparison leaves out the 12 diagnostic variants, so it does not rest on the variants that define the type. Bars, 2.5th to 97.5th percentiles. n = 2,831 and 2,838 within-person pairs of terminal-type units (+diag., −diag.), 1,392 and 1,353 of ordinary units, and 20,000 between-person pairs per group, within 541 haplotypes of 299 individuals. (O) Terminal-type carriage by superpopulation. Carriage per haplotype and per individual (carried on either haplotype). Denominators (haplotypes/individuals), AFR 167/91, SAS 110/61, AMR 86/47, EAS 103/56 and EUR 67/40. n = 533 haplotypes of 295 individuals, those of the 541 matched to the 1000 Genomes population table. (P) Number of depths occupied by terminal-type units. For each size (1-6) of the depth set occupied by terminal-type units, the fraction of a population’s haplotypes carrying a set of that size. Color, superpopulation. Denominators (haplotypes, carriers), AFR 167 and 9, SAS 110 and 45, AMR 86 and 38, EAS 103 and 53, EUR 67 and 37. The 2 carriers without a population label are omitted. n = 182 carriers among 533 haplotypes of 295 individuals. (Q) Terminal-type units in cell-line assemblies. Distal depths 1-20 in four cell-line assemblies (GM12878 A and B, K562 A and B, the two haplotypes of each line). Red points, terminal-type units, called by the same rule after aligning each unit to the reference unit. Cell lines only, so no statement about inheritance follows. n = 80 units from 4 assemblies of 2 cell lines.

**Supplementary Figure S3.**
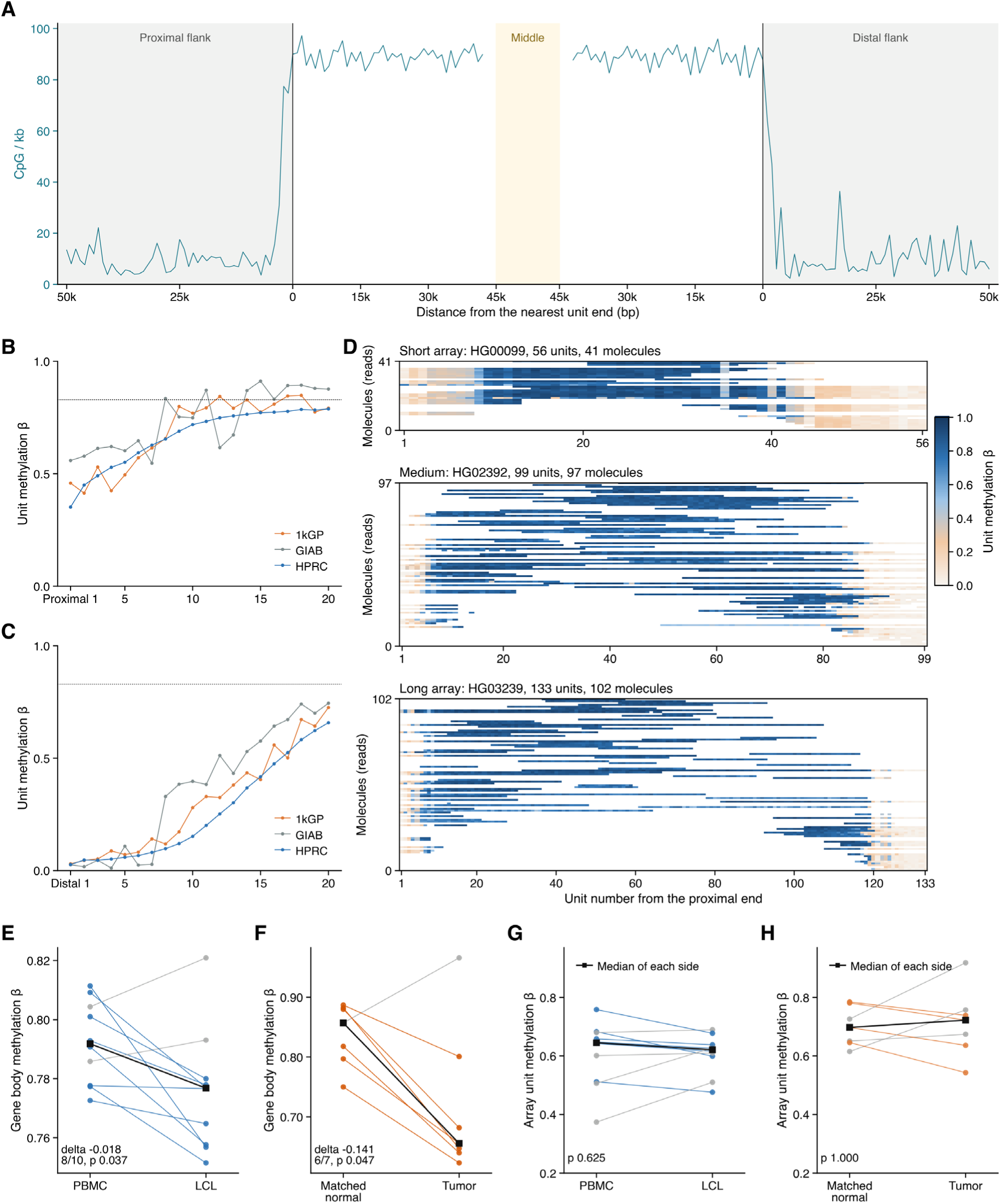
The positional methylation pattern checked against unit sequence, array length, data source and cell state. (A) CpG density of the array sequence. CpG per kb in 1,000 bp windows, on the same folded axis as Figure 3A. One bin merges the last 1 kb inside the array with the first 1 kb outside it, because the binning takes the floor of the absolute distance and then restores the sign. n = 186 windows of 1,000 bp from 487 haplotypes of 268 individuals. (B) and (C) Mean unit methylation by position in three data sources. Unit mean β against distance from each end for 1kGP, GIAB and HPRC. (B) counts from proximal 1 and (C) from distal 1. The dotted line, mean of the interior zone (0.829). The three sources are not three sets of people. Of the 10 1kGP individuals 9 are also in HPRC. Both GIAB haplotypes of (B) come from one individual, so the roughness of that trace is from sample size. n = 47,206 (HPRC), 485 (1kGP) and 165 (GIAB) units in (B), from the 507, 6 and 2 haplotypes whose proximal end is observed, and 47,483 (HPRC), 1,265 (1kGP) and 228 (GIAB) units in (C). (D) Single molecule maps of three haplotypes of different length. Drawn as Figure 3E, for HG00099, HG02392 and HG03239. Within each length stratum the haplotype with the most molecules was taken, from the 220 haplotypes that passed fixed thresholds. n = 41, 97 and 102 molecules over 56, 99 and 133 units. (E) Blood and cell line pairs from the same donor. Gene-body β within high-methylation units, peripheral blood mononuclear cells against the lymphoblastoid cell line of the same person. Within-pair median change −0.018, 8 of 10 fall, *P* = 0.037. n = 10 pairs of 10 donors, the HPRC Year 7 matched set. (F) Paired normal and tumor samples. The same measurement in 7 matched normal-tumor pairs. The tumor side is a cell line in all seven and the normal side is a matched cell line in six and patient tissue in one. Within-pair median change −0.141, 6 of 7 fall, Wilcoxon signed rank *P* = 0.047 one sided and *P* = 0.094 two sided. n = 7 pairs of 7 donors, five from the CASTLE R10 ONT release, one from the 2024 COLO829 ONT release and one from the GIAB HG008 ONT release. The gene body is the 119 bp at the end of the unit. (G) and (H) The same two sets of pairs measured on whole-array β. The 10 pairs of (E) and the 7 pairs of (F) with the y axis changed from gene-body β to the unit β of the whole array. Heavy black line, median of each side. Blood and cell line, 0.645 to 0.621, within-pair median change −0.021, 6 of 10 fall, *P* = 0.625. Normal and tumor, medians 0.698 to 0.723 with a within-pair median change of −0.046, 4 of 7 fall, *P* = 1.000; the median of the paired changes and the difference of the two medians need not share a sign. n = 10 pairs of 10 donors (G) and 7 pairs of 7 donors (H), as in (E) and (F).

**Supplementary Figure S4.**
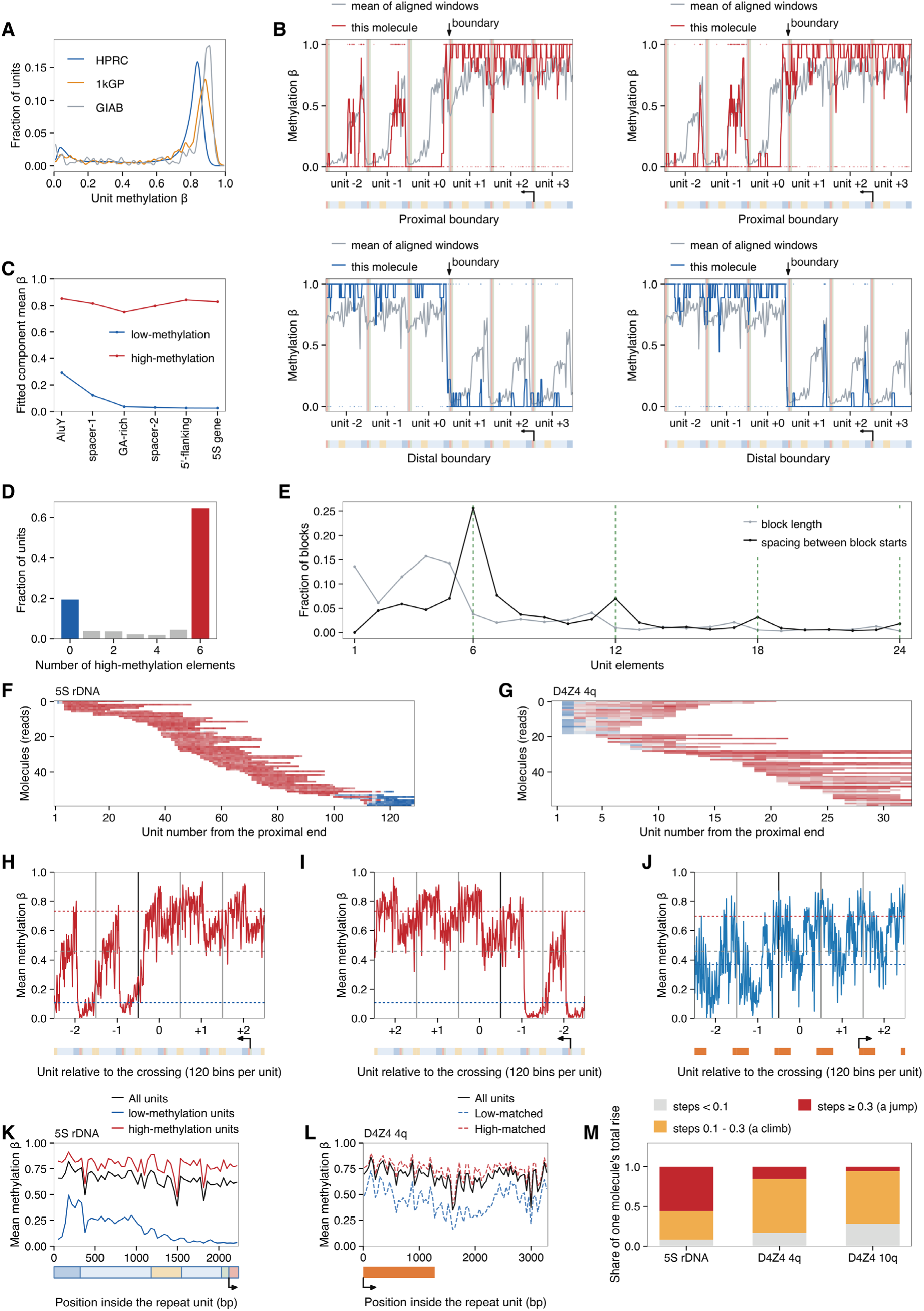
Detailed unit methylation states in 5S rDNA and D4Z4. (A) Unit methylation distribution in three datasets. HPRC (blue), 1kGP (orange) and GIAB (gray). n = 47,483 (HPRC), 1,265 (1kGP) and 228 (GIAB) units. (B) Single molecules through a boundary. Two molecules through the proximal boundary (top, red) and two through the distal boundary (bottom, blue). Each unit is drawn from the 5S gene to AluY. n = 4 molecules. (C) Fitted state levels by sub-unit element. The two components are fitted separately for each of the six elements. n = 833,387 to 843,297 read-units per element from 576 haplotypes of 297 individuals, HPRC. (D) Number of high-methylation elements per unit. Units with all six elements called, of which 83.9% are wholly in one state (0 or 6 elements). n = 816,172 read-units from 576 haplotypes of 297 individuals, HPRC. (E) Spacing between block starts of the same state. Black, spacing between successive starts of same-state blocks within one read, with peaks at 6, 12, 18 and 24 sub-unit elements, multiples of one unit. Gray, length of blocks that contain the 5S gene element. Dashed green lines, multiples of six elements. Of all spacings, 42.4% are multiples of six against 16.7% expected by chance. n = 260,044 spacings (black) and 169,011 blocks (gray). (F) Example read-by-unit map of 5S rDNA. n = 60 reads, the 1kGP set. (G) Example read-by-unit map of D4Z4 4q. n = 60 reads, the 1kGP set. (H) Transition at the proximal end of 5S rDNA at base resolution. Molecules are aligned on their own crossing, defined as in Figure 4, and averaged in 120 bins per unit over the two units on each side, so the trace is a superposition and not a single molecule. Units are numbered from the crossing, negative toward the array end. Red and blue dotted lines, the high and the low fitted state levels (0.7324 and 0.1078), and the gray dashed line the cut, β = 0.4611, which is the likelihood ratio split between the two fitted components and so does not fall midway between their levels. n = 104 molecules, the 1kGP set. (I) Transition at the distal end of 5S rDNA at base resolution. The array interior is on the left and the distal end on the right, with the state levels of (H). n = 139 molecules, the 1kGP set. (J) Transition at the proximal end of D4Z4 4q at base resolution. Dotted lines, the D4Z4 4q state levels (0.6970 and 0.3679), the gray dashed line is the 5S rDNA cut, drawn here for reference only, and orange bars mark the transcribed part. n = 53 molecules, the 1kGP set. (K) Within-unit methylation of the two unit classes in 5S rDNA. All units (black), low-methylation units (blue) and high-methylation units (red). n = 25,644 units, the 1kGP set. (L) Within-unit methylation of the two unit classes in D4Z4 4q. All units (black) and the two sides of the rank split of Figure 4J (dashed, blue low-matched and red high-matched), the labels of that split carried over unchanged. n = 7,966 units, the 1kGP set. (M) Decomposition of each molecule’s total rise by step size. 5S rDNA, D4Z4 4q and D4Z4 10q. n = 290, 179 and 122 rising molecules, the 1kGP set.

**Supplementary Figure S5.**
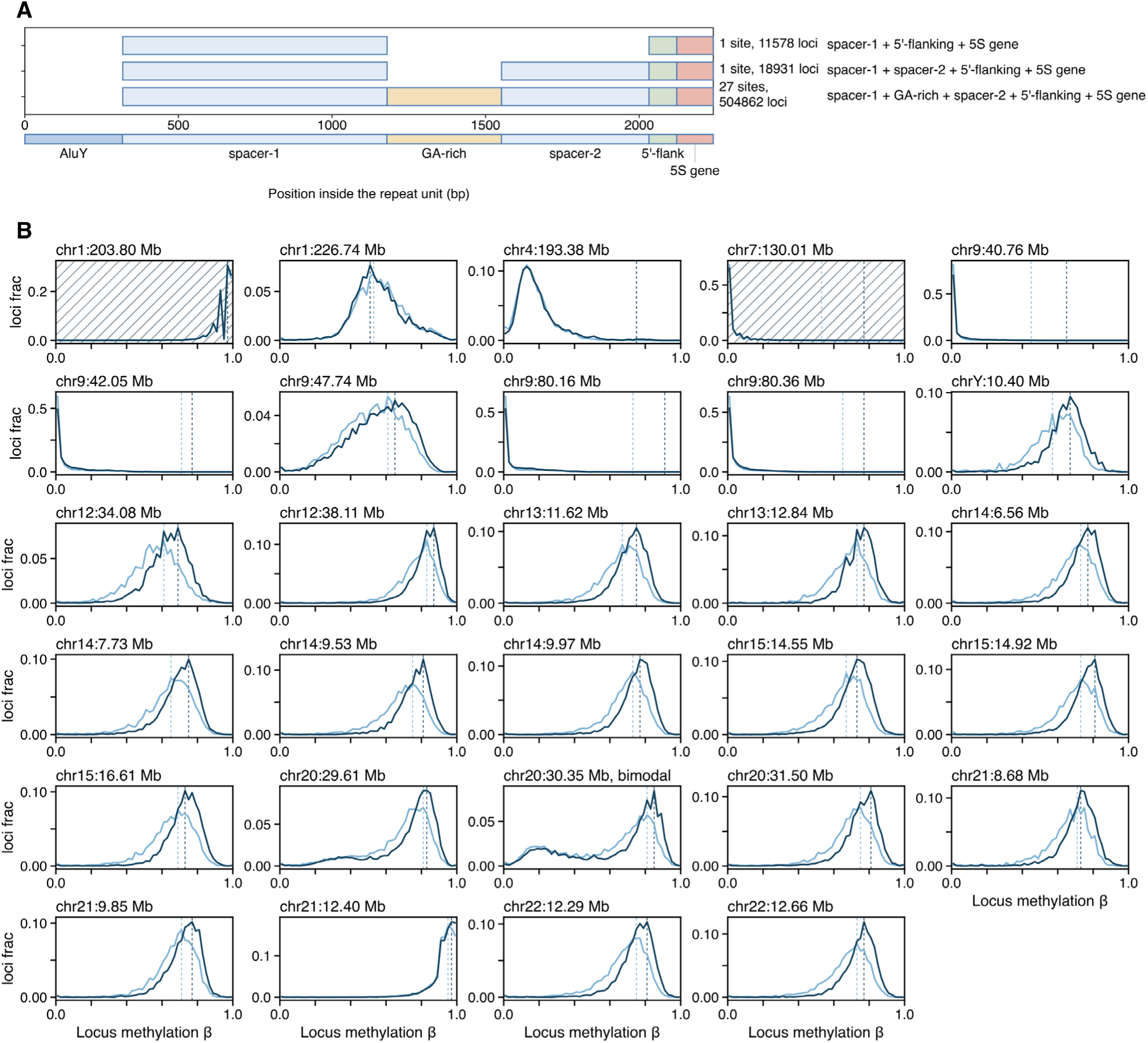
Dispersed 5S copies outside the array. (A) Element content of the dispersed 5S copies. Copies at the 29 sites that carry at least three elements of the 5S rDNA repeat unit, drawn on the coordinate of the array unit. Most sites, 27 of 29 (504,862 loci), carry spacer-1, the GA-rich segment, spacer-2, the 5′ flank and the 5S gene, that is, the whole unit except the AluY element. The other two lack the GA-rich segment, and one of them also lacks spacer-2. n = 535,371 loci at 29 sites from 305 individuals, the set from which (B) takes its loci. (B) Locus methylation at each dispersed site by flank tone. Distribution of locus methylation β at each of the 29 sites, drawn separately for individuals in the lowest third (light blue) and the highest third (dark blue) of flank tone, an individual’s mean methylation over the CpGs on the same reads that lie more than 500 bp outside the 5S-like segment. The middle third is not drawn. β is the hard call (each CpG counted 0 or 1 at ML ≥ 128), and a locus needs at least 10 CpGs. Dashed lines, the high-methylation mode of each group. Hatched panels are the two sites whose loci carry a median of fewer than 50 CpGs, so that locus β moves in steps wider than the 0.02 bin and the curve is jagged. n = 355,564 of the 535,371 loci of (A), 162,372 in the lowest and 193,192 in the highest tone third.

**Supplementary Figure S6.**
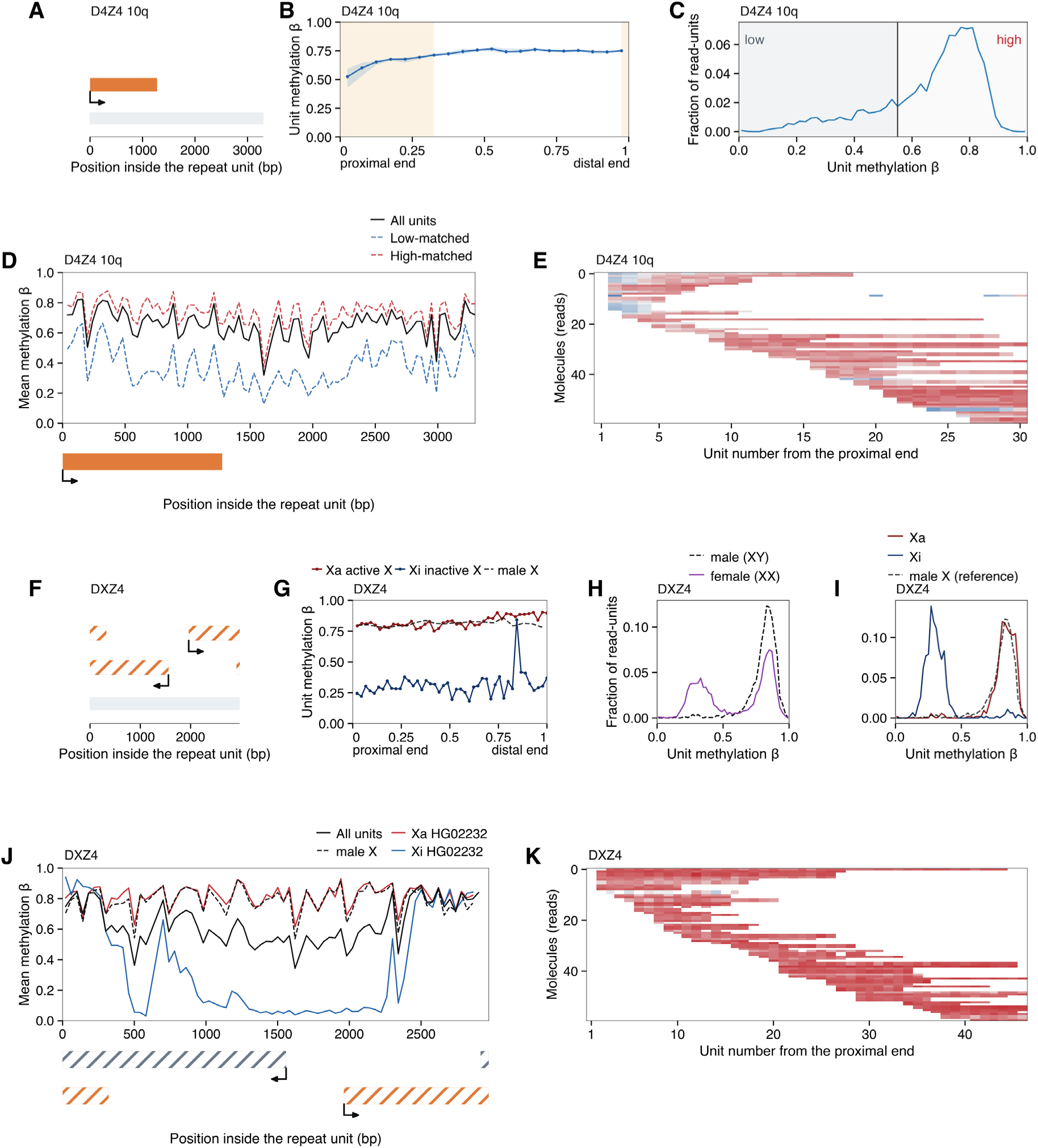
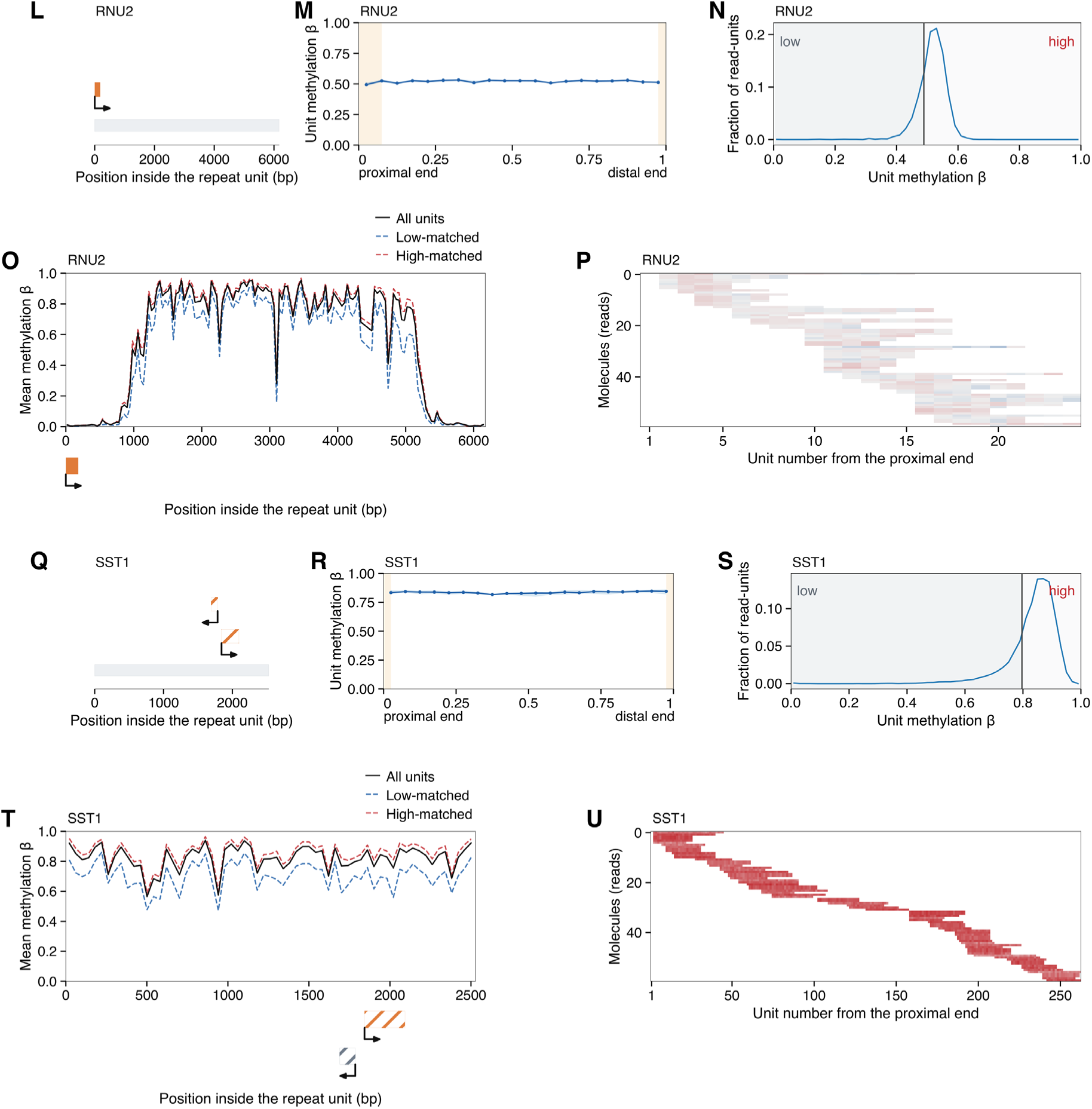
Unit methylation of four other human tandem arrays. (A) D4Z4 10q, transcribed parts of the repeat unit. (B) D4Z4 10q, methylation along the array. n = 6 segments from 6 haplotypes of 4 individuals, the 1kGP set. (C) D4Z4 10q, unit methylation distribution. n = 4,998 units, the 1kGP set. (D) D4Z4 10q, within-unit methylation. n = 4,998 units, the 1kGP set. (E) D4Z4 10q, read-by-unit map. n = 60 reads, the 1kGP set. (F) DXZ4, transcribed parts of the repeat unit. (G) DXZ4, methylation along the array for the active and the inactive X of one woman. n = 4 haplotypes (the Xa and the Xi of HG02232 and 2 male X), the 1kGP set. (H) DXZ4, unit methylation distribution in men and in women. n = 2,118 units in men and 7,693 in women other than HG02232, the 1kGP set. (I) DXZ4, unit methylation distribution for the active and the inactive X of one woman. n = 534 units on the Xa and 467 on the Xi, the 1kGP set. (J) DXZ4, within-unit methylation. n = 10,812 units, the 1kGP set. (K) DXZ4, read-by-unit map. n = 60 reads, the 1kGP set. (L) RNU2, transcribed parts of the repeat unit. (M) RNU2, methylation along the array. n = 4 segments from 4 haplotypes of 3 individuals, the 1kGP set. (N) RNU2, unit methylation distribution. n = 4,950 units, the 1kGP set. (O) RNU2, within-unit methylation. n = 4,950 units, the 1kGP set. (P) RNU2, read-by-unit map. n = 60 reads, the 1kGP set. (Q) SST1, transcribed parts of the repeat unit. (R) SST1, methylation along the array. n = 18 segments from 16 haplotypes of 9 individuals, the 1kGP set. (S) SST1, unit methylation distribution. n = 42,554 units, the 1kGP set. (T) SST1, within-unit methylation. n = 42,554 units, the 1kGP set. (U) SST1, read-by-unit map. n = 60 reads, the 1kGP set. In the transcription panels (A, F, L, Q), as in Figure 4H, the gray bar is the repeat unit, orange bars the transcribed parts and the bent arrow the transcription start and its direction. Under each within-unit profile (D, J, O, T) every transcript has its own row, orange for the plus and gray for the minus strand. Hatched bars are transcripts whose annotated extent is not established at base resolution, so their drawn ends are approximate, and a transcript that runs past the end of the unit appears as two blocks on one row (F, J). In the profiles along the array (B, M, R) points are the median over array segments in 20 equal bins of relative position, 0 proximal and 1 distal with the ends not folded, the band is the interquartile range and shading marks the transition zones as in Figure 4I. In the unit β distributions (C, N, S) none of these arrays has a valley to cut at, so the vertical line is a rank split that leaves 21.9% of units below it, the share of low-methylation units in 5S rDNA (Figure 4J), and the within-unit profiles (D, O, T) show all units (solid black) and the two sides of that split (dashed, blue low-matched and red high-matched). The read-by-unit maps (E, K, P, U) show, for the haplotype with the most reads, the reads that cover the most units, ordered by their first unit and colored by unit β from blue (0) through gray (0.5) to red (1). DXZ4 is laid out by X chromosome, because its two modes are the active and the inactive X. In (G) and (I) the active (Xa) and the inactive (Xi) X are those of an example woman, HG02232, and male X in (G) is the median of the two male haplotypes whose segments reach flanking sequence at both ends. In this woman each X sits at its own level, and the difference is maintained along the array (G, I). Panel (H) compares men (XY) and women (XX), and panel (J) shows all units, male X, and the Xa and the Xi of HG02232. The figure continues over two pages, (A)-(K) on the first and (L)-(U) on the second.

**Supplementary Figure S7.**
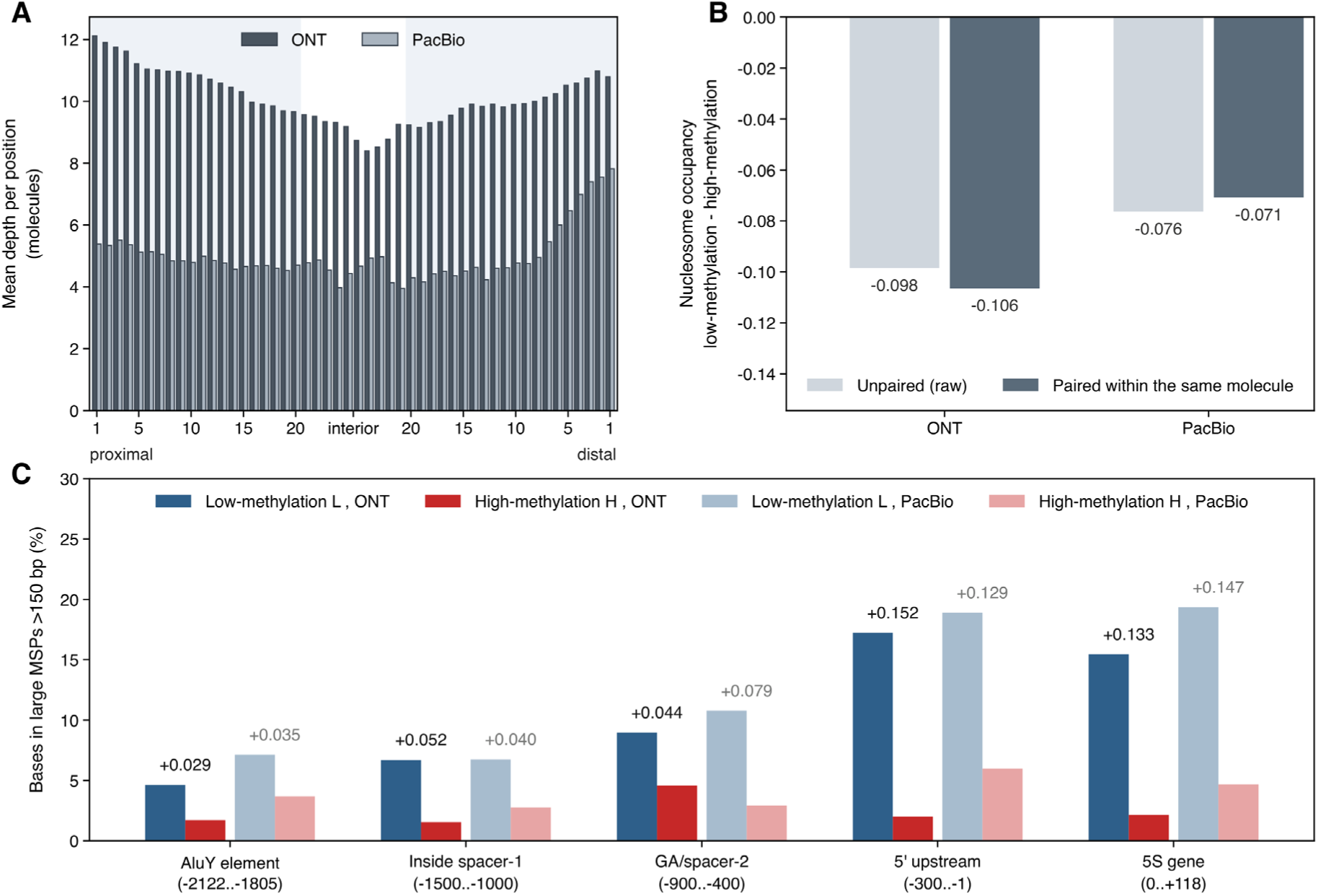
Molecule depth, the effect of pairing on the occupancy difference, and element-wise MSP density across the 5S rDNA array. (A) Mean molecule depth per array position. The number of read-by-unit observations in a bin divided by the number of unit positions pooled in that bin, ONT in dark bars (52 individuals) and PacBio in light bars (48 individuals), the two platforms counted separately. The axis runs from the proximal to the distal end of the array. The 20 bins at either end are absolute positions, one bin per unit and numbered from the corresponding end starting at 1, and the 10 central bins hold the interior as relative positions. This panel is the denominator of the per-position panels of Figure 5 and pools all molecules. (B) Effect of pairing on the nucleosome occupancy difference between low-methylation and high-methylation units. Low-methylation minus high-methylation, shown both unpaired (raw, light bars) and paired within the same molecule (dark bars), ONT and PacBio separately. Units were classified at a unit CpG β of 0.508 and occupancy was taken from FiberHMM segments. Low-methylation minus high-methylation is −0.098 unpaired and −0.106 paired on ONT, and −0.076 unpaired and −0.071 paired on PacBio. The value of each bar is printed beside it. Bar heights are not compared between the unpaired and the paired estimate, because changing the boundary alters both the control on labeling and the methylation contrast within the paired molecules. The paired estimate is the test of Figure 5G and H. The unpaired estimate was tested over individuals, each contributing the mean of its low-methylation and of its high-methylation units, by a two-sided Wilcoxon signed-rank test on the exact null distribution, *P* = 4 × 10⁻¹⁶ on ONT and *P* = 5 × 10⁻¹⁴ on PacBio. ONT 52 individuals and PacBio 48 individuals. (C) Element-wise large-MSP density in low-methylation and high-methylation units. Low-methylation L in blue and high-methylation H in red, ONT in dark bars and PacBio in light bars, resolved into five windows of the unit, the AluY element, inside spacer-1, GA and spacer-2, the 5′ upstream region and the 5S gene. The coordinates in parentheses are relative to the transcription start site of the 5S gene. The numbers above the bars are the low-methylation minus high-methylation difference. Tests were unpaired, over all units of each class, and the paired estimates are in Figure 5F and in (B) above. The AluY window is counted using only bases belonging to the unit itself. Units were classified at a unit CpG β of 0.508, MSPs were called with FiberHMM, and ONT 52 individuals and PacBio 48 individuals were counted separately.

**Supplementary Figure S8.**
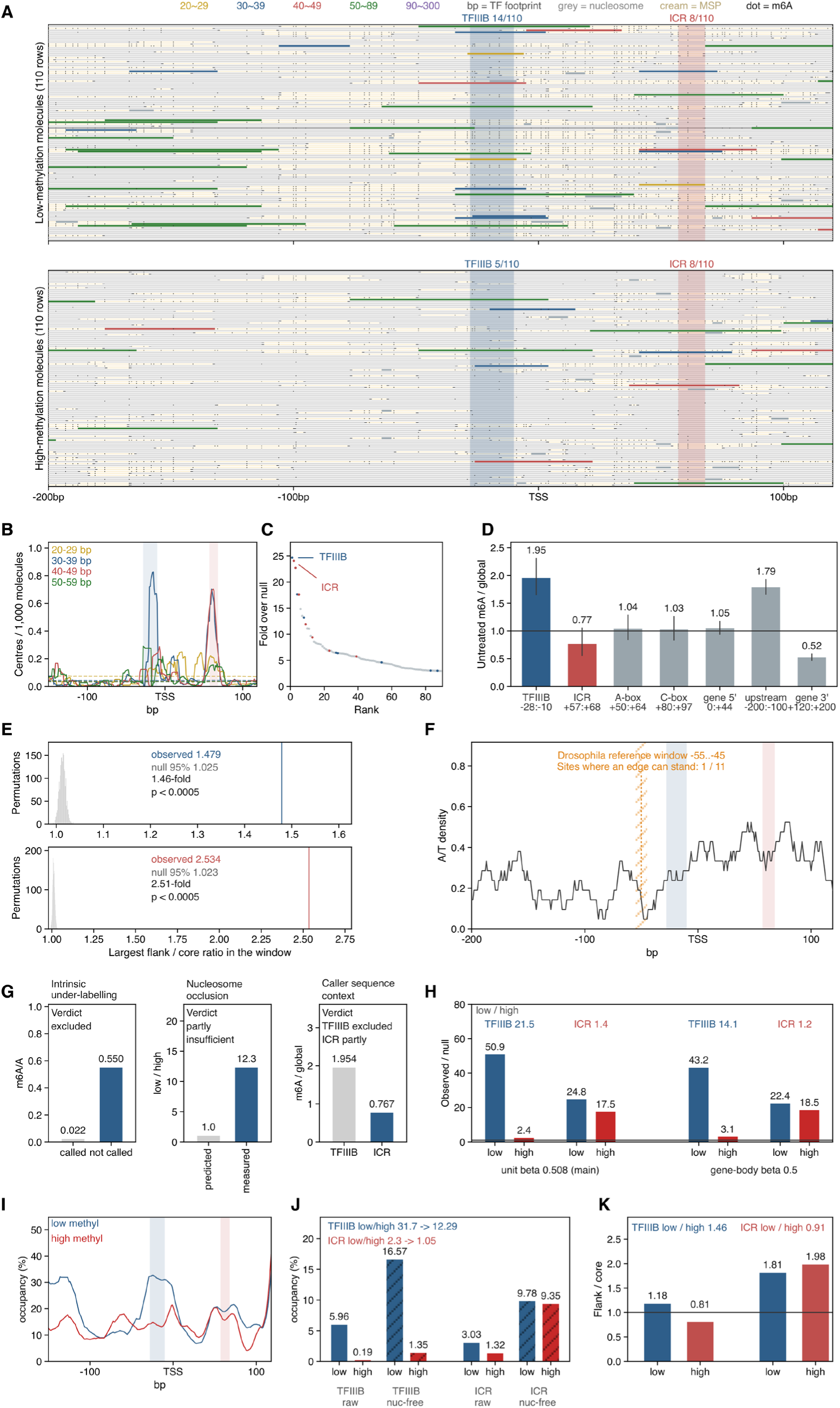
Single-molecule views, controls and alternative explanations for the footprint analysis. (A) Single molecules by methylation stratum. 110 molecules per stratum, drawn at random by reservoir sampling from the molecules that span the displayed window in full, the same number taken from each stratum and left unsorted in the order drawn. This is a display sample and not a denominator. 14 of 110 low-methylation molecules carry a TFIIIB call against 5 of 110 high, and the internal control region gives 8 against 8. (B) Footprint-center density by size class, without null normalization. Per 1,000 molecules for four size classes, pooled and without the normalization over the null used in Figure 6C. The dashed horizontal lines are the per-class mean densities, not a null level. (C) Ranking of every grid cell that cleared the threshold. All 89 grid cells from Figure 6B that cleared the threshold 2.94, ranked by fold excess, with TFIIIB and the internal control region in color and the other 87 gray. The two bins used in Figure 6D are ranks 1 and 2 here (24.68, 24.07). They were fixed before methylation was looked at, and the ranking runs over the whole scan with no position specified in advance. (D) Untreated control material from an ONT run. m6A over the global rate by window. TFIIIB 1.95, the opposite of what a sequence-context artifact predicts, against 0.77 at the internal control region, which is partly consistent with it. (E) Permutation null at the two positions. The null is built from 2,000 permutations at the two sites alone, not over the whole grid. TFIIIB 1.479 observed against a null median 1.013 and 95th percentile 1.025 (*P* < 5 × 10⁻⁴), internal control region 2.534 against 1.011 and 1.023. (F) Adenine opportunity across the scan window. Hia5 methylates adenine only, so a footprint edge cannot fall where there is none, and positions were evaluable only above 1,000 adenine opportunities summed over both strata. A window at −55 to −45, the position reported in *Drosophila* and carried here as a reference point, meets that cut at 1 of 11 positions, against 4 of 19 for TFIIIB (−28 to −10) and 3 of 12 for the internal control region (+57 to +68). (G) The three alternative explanations and their verdicts. Intrinsic under-labeling gives m6A per adenine of 0.022 inside called footprints against 0.550 outside, and is excluded. Nucleosome occlusion, a low-to-high ratio of 1.0 predicted against 12.3 measured, partly insufficient. Caller sequence context, untreated m6A over global on ONT 1.954 at TFIIIB against 0.767 at the internal control region, the same measurement as in (D) printed there to two decimals, excluded for TFIIIB and partly for the internal control region. (H) The same measurement under the two unit labels. Whole-unit β 0.508 and gene-body β 0.5. TFIIIB 50.9/2.4 = 21.5 and 43.2/3.1 = 14.1, internal control region 1.4 and 1.2. The labels agree for 91.6% of 29,855 units but share no denominator, 7,451/22,879 molecules for whole-unit β against 9,255/20,600 for gene-body β, so ratios taken under different labels are not comparable. The disagreement runs one way. Of the units the two labels place differently, 2,212 are high on the whole unit and low in the gene body against 290 the other way round. (I-K) Molecules carrying no nucleosome call. In each of these three panels the nucleosome is handled by restricting the denominator rather than by subtracting a contribution, and the solid horizontal line marks the null level of 1. (I) Footprint frequency per position on nucleosome-free molecules. Fraction of molecules carrying a TFIIIB-sized call at each position, low against high methylation, among molecules with no nucleosome call over that position. (J) Occupancy at the two site windows, raw and nucleosome-free. The same as occupancy at the two positions. TFIIIB 5.96/0.19% and 16.57/1.35%, the low- to-high ratio falling from 31.7 to 12.3, far above the 1.0 that pure occlusion predicts. Internal control region 3.03/1.32% and 9.78/9.35%, falling from 2.31 to 1.05. (K) The caller-free axis on nucleosome-free molecules. TFIIIB 1.18 against 0.81 (1.46-fold), internal control region 1.81 against 1.98 (0.91-fold). Ratios in this figure are computed from the unrounded values, not from the rounded occupancies printed beside them.

**Supplementary Figure S9.**
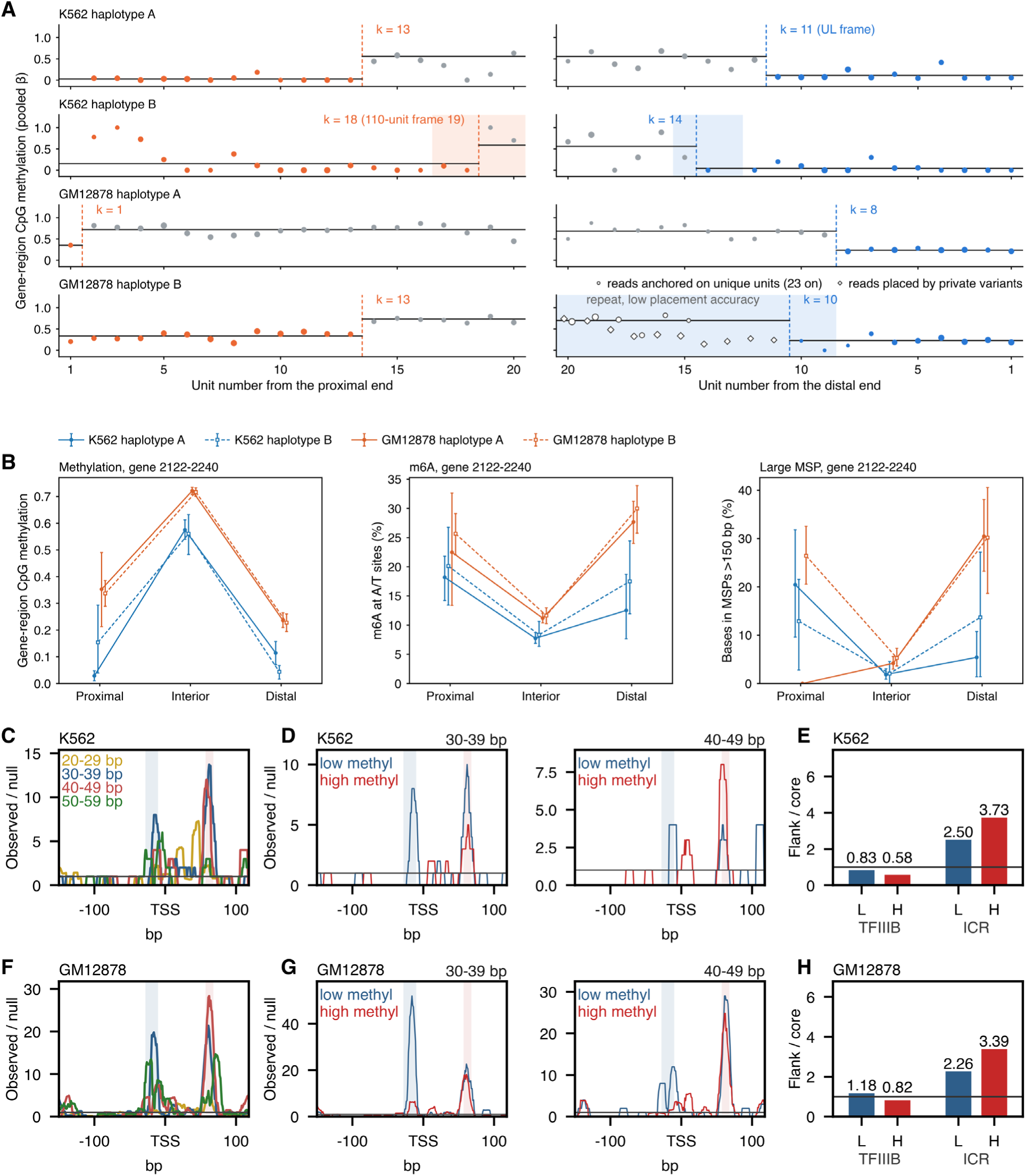
End zones and Fiber-seq footprints in K562 and GM12878. (A) Change points in gene-region methylation at the array ends. Gene-region CpG methylation (pooled β) of the 20 units nearest each end is plotted against unit number from the proximal end (left) and from the distal end (right) for the four cell-line haplotypes. Point size shows the number of CpG calls in the unit, its area growing with the square root of that number relative to the largest unit in the panel, so larger points rest on more calls. Orange (proximal) and blue (distal) points are units inside the low-methylation stretch, and gray points are units beyond it. The dashed line marks the fitted change point k and the horizontal lines the pooled β on either side of it. The shaded span, orange at the proximal end and blue at the distal end, is the 95% interval of k and is drawn only where it is wider than one unit, at both ends of K562 haplotype B and at the distal end of GM12878 haplotype B, where it runs past the 20 units shown. In GM12878 haplotype B the distal end lies in a repeat where reads are placed with low accuracy, and open symbols mark units there that received no placed read and are measured instead from reads anchored on unique units (circles) or placed by private variants (diamonds). K562 haplotype A is shown at its distal end on the unit frame of a second K562 assembly built with the ONT reads added, the UL frame, because in the assembly used elsewhere units 14-39 at that end received no placed read. (B) Gene-body measures by zone. From left, CpG methylation over the gene body (2,122-2,240), m6A at A/T sites over the gene body and the share of gene-body bases covered by a large MSP (> 150 bp), with 95% intervals. Methylation is the quantity the zones were cut on and is shown only as a check. (C) Footprint profiles by size class in K562. Curves for the four size classes, 20-29, 30-39, 40-49 and 50-59 bp, over all units, as observed over a circular-shift null with 11 bp smoothing. Blue and red bands, TFIIIB (−28 to −10) and the internal control region (+57 to +68). All reads touching the array are used without read placement. (D) Footprint profiles in low- and high-methylation units in K562. Shown for the 30-39 bp class (left) and the 40-49 bp class (right), drawn as in (C). Units are labeled low or high at gene-body β 0.5 (1,106 low and 1,228 high), and the bar summary is Figure 6G. (E) Caller-free flank-to-core m6A ratio at the two positions in K562. The core window is c−7 to c+1 and the flanks c−35 to c−27 and c+22 to c+27, the asymmetric geometry of Dubocanin and colleagues kept unchanged, around centers −11 (TFIIIB) and +58 (internal control region). (F-H) The same three views for GM12878, footprint profiles by size class (F), footprint profiles by methylation state (G, 1,572 low and 3,856 high units, bar summary in Figure 6H) and the flank-to-core ratio (H).

**Supplementary Figure S10.**
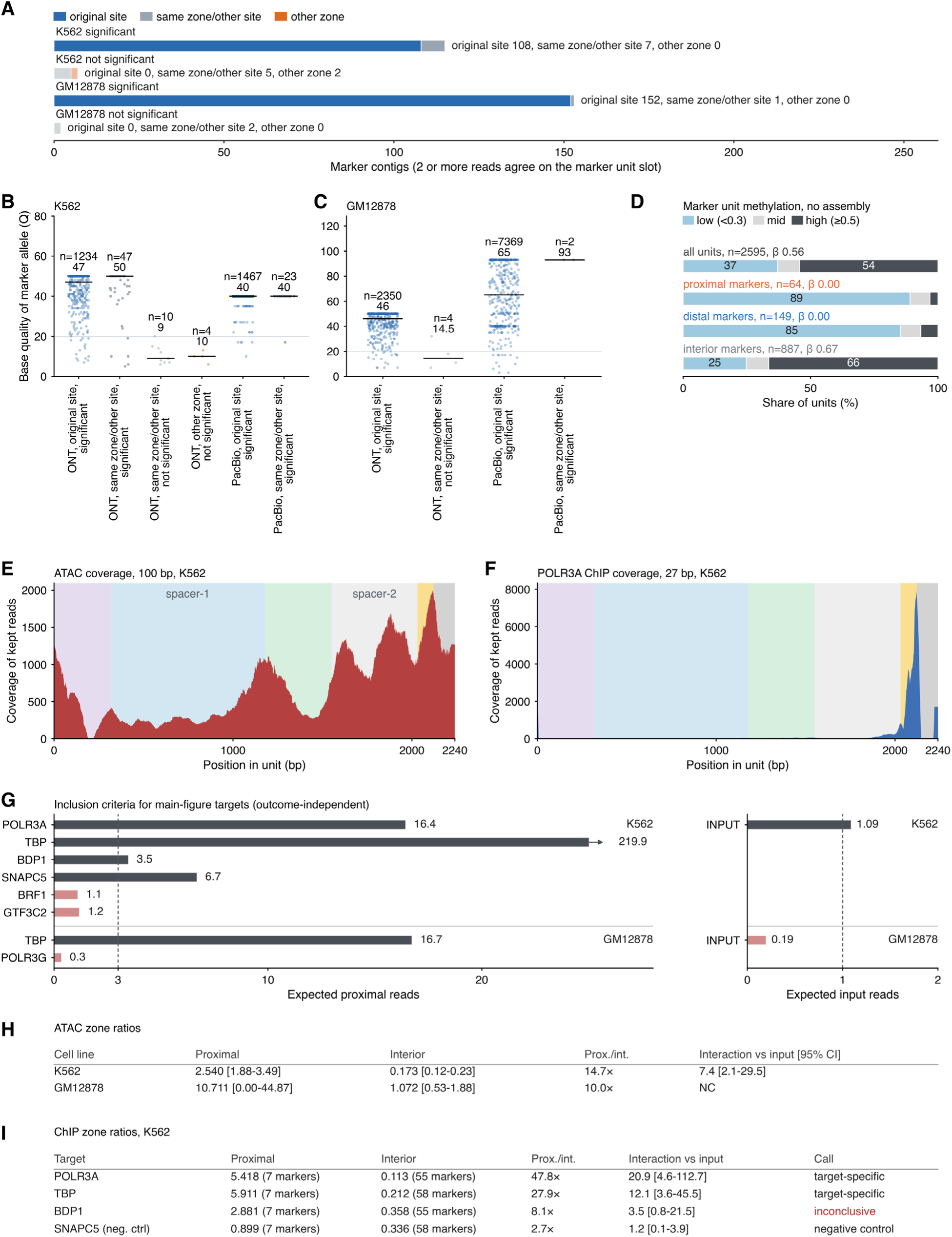
Validation of zone-exclusive markers and zone ratios of ATAC-seq and ChIP-seq. (A) Reassembly from marker-allele reads. Contigs assembled from marker-allele reads alone, with at least two reads agreeing on the marker’s unit slot, are classed as lying at the original site, at another site in the same zone (pooled with contigs outside the assembly or ambiguous) or in another zone. A contig is counted only when it rests on at least two independent reads, has at least two reads with marker-base quality ≥ 20 and carries at most 20% discordant pairs. (B, C) Base quality of the marker allele in the reassembled contigs, K562 (B) and GM12878 (C). Reads are grouped by platform and class, black bars mark medians and the gray line marks quality 20. (D) Methylation of marker-carrying units. A unit enters with at least three methylation calls and is called low-methylation below a unit CpG β of 0.3. Units are called from placed reads alone, without an assembly. (E) ATAC-seq coverage within the unit in K562. Coverage is shown in a 100 bp window, after the alignment gate described in Materials and methods. (F) POLR3A ChIP-seq coverage within the unit in K562. The window is 27 bp. (G) Inclusion criteria for the main-figure ChIP-seq targets in K562 and GM12878. The criteria, fixed before any outcome was known, ask whether a target has enough proximal-marker reads for a zone ratio. The expected count sums, over the proximal markers (seven in K562 and three in GM12878), the ChIP-seq depth at the marker position times the WGS marker-allele fraction (base quality ≥ 20), without using the observed marker reads. Targets need at least 3, and the control track, scored the same way, needs at least 1 so that the interaction against the control track can be computed. In K562 expected proximal reads are 16.4 for POLR3A, 219.9 for TBP, 3.5 for BDP1 and 6.7 for the negative control SNAPC5, against 1.1 for BRF1 and 1.2 for GTF3C2, and 1.09 for the control track. In GM12878 they are 16.7 for TBP and 0.3 for POLR3G, and 0.19 for the control track. (H) ATAC-seq zone ratios in both cell lines. Zone ratios are observed over expected marker reads with 95% bootstrap intervals, Prox./int. is the proximal zone ratio over the interior one, counted without a base-quality filter as in Figure 6K. The interaction is the proximal-to-interior ratio over that of the control track. NC, not computable, because the GM12878 control track has one proximal observation. (I) ChIP-seq zone ratios in K562. Columns are as in (H), counted at base quality ≥ 20 as in Figure 6L. A target is called target-specific when the 95% interval of its interaction lies above 1, and inconclusive otherwise. BRF1 and GTF3C2, a TFIIIC subunit, are not shown because they fail the criterion of (G); their counts are given in Supplementary Table S8.

**Supplementary Figure S11.**
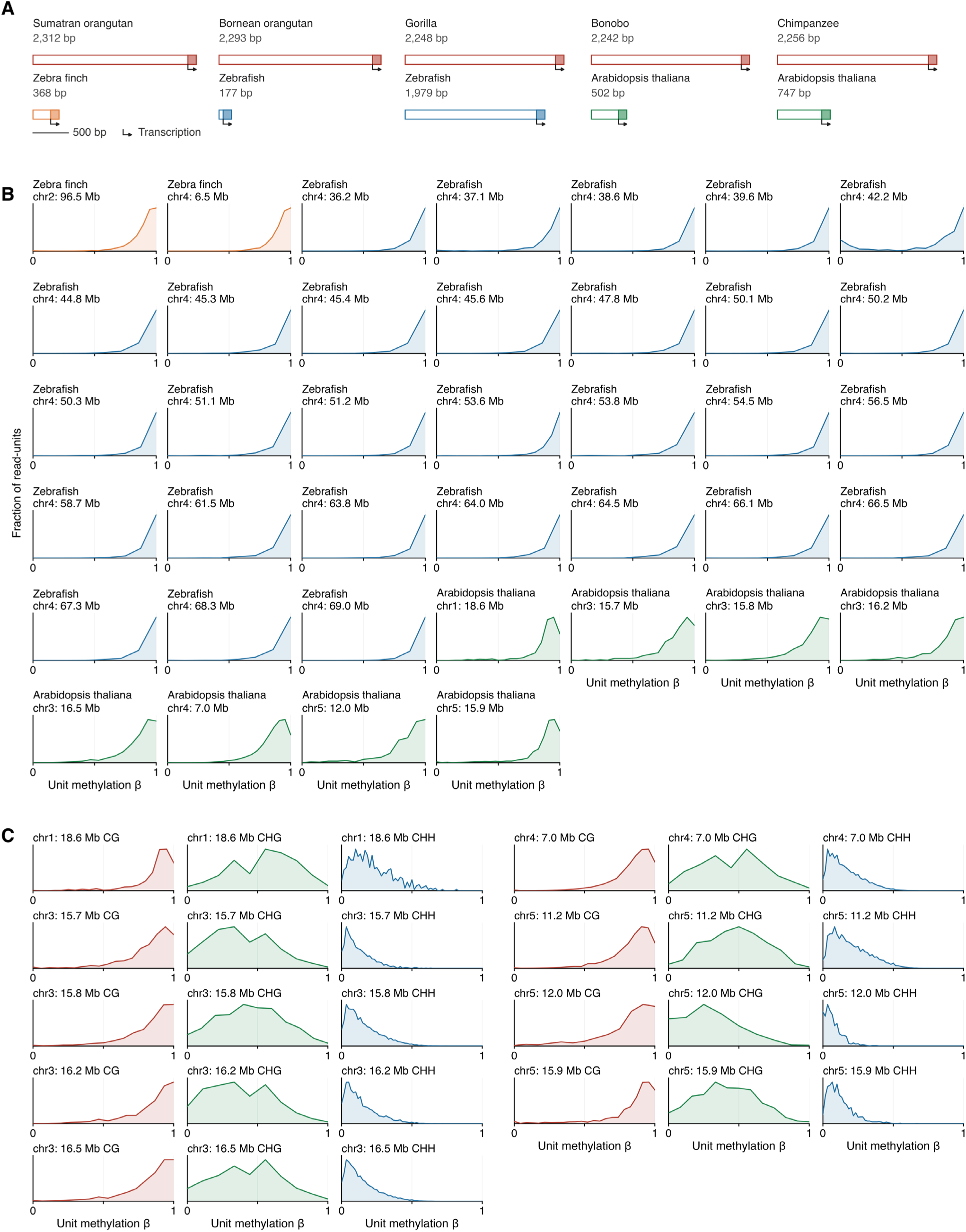
Repeat unit of the representative arrays, per-unit methylation of the arrays not shown in Figure 7, and the three sequence contexts of *Arabidopsis thaliana*. (A) Structure of the repeat unit in each species. Repeat unit of the representative array of each species, all on one bp scale, each unit headed by the species and by its own length in bp. The block is the 5S rDNA gene, which lies at the end of the unit in every species and is drawn in a lighter shade of that species color. A species whose arrays fall into two clearly different unit lengths is drawn twice, once for each representative length. Unit length is 2,242-2,312 bp in apes, 368 bp in zebra finch, 177 bp in zebrafish with one array of 1,979 bp, and 482-747 bp in *Arabidopsis thaliana*, while the gene itself is 118-119 bp in all of them. (B) Per-unit methylation of the 39 arrays not shown in Figure 7C. The 39 are zebra finch 2, zebrafish 29 and *A. thaliana* 8, drawn as Figure 7C, over 662,884 complete units. Per species that is 52,647 units in zebra finch, 430,819 in zebrafish and 179,418 in *A. thaliana*, and a single array holds from 508 to 67,526. None of the 39 is bimodal. Median array β is 0.910 in zebra finch, 0.956 in zebrafish and 0.859 in *A. thaliana*, and the lowest single array is 0.741. (C) The three sequence contexts of the nine *Arabidopsis thaliana* arrays. Per-unit methylation measured separately in the CG, CHG and CHH contexts. Each row is one array and each column one context, the nine arrays are split into two blocks of five and four drawn side by side, and the vertical axis is the fraction of read-units and is not labeled on the panels. β is the hard-called methylation of one read × unit in that context, binned as in Figure 7C. Averaged over the nine arrays, each array taken as its call-weighted mean β (Σ5mC / Σcalls), the unit methylation is CG 0.844, CHG 0.423 and CHH 0.146, with a median of 17 CG, 9 CHG and 69 CHH calls per unit.

**Supplementary Figure S12.**
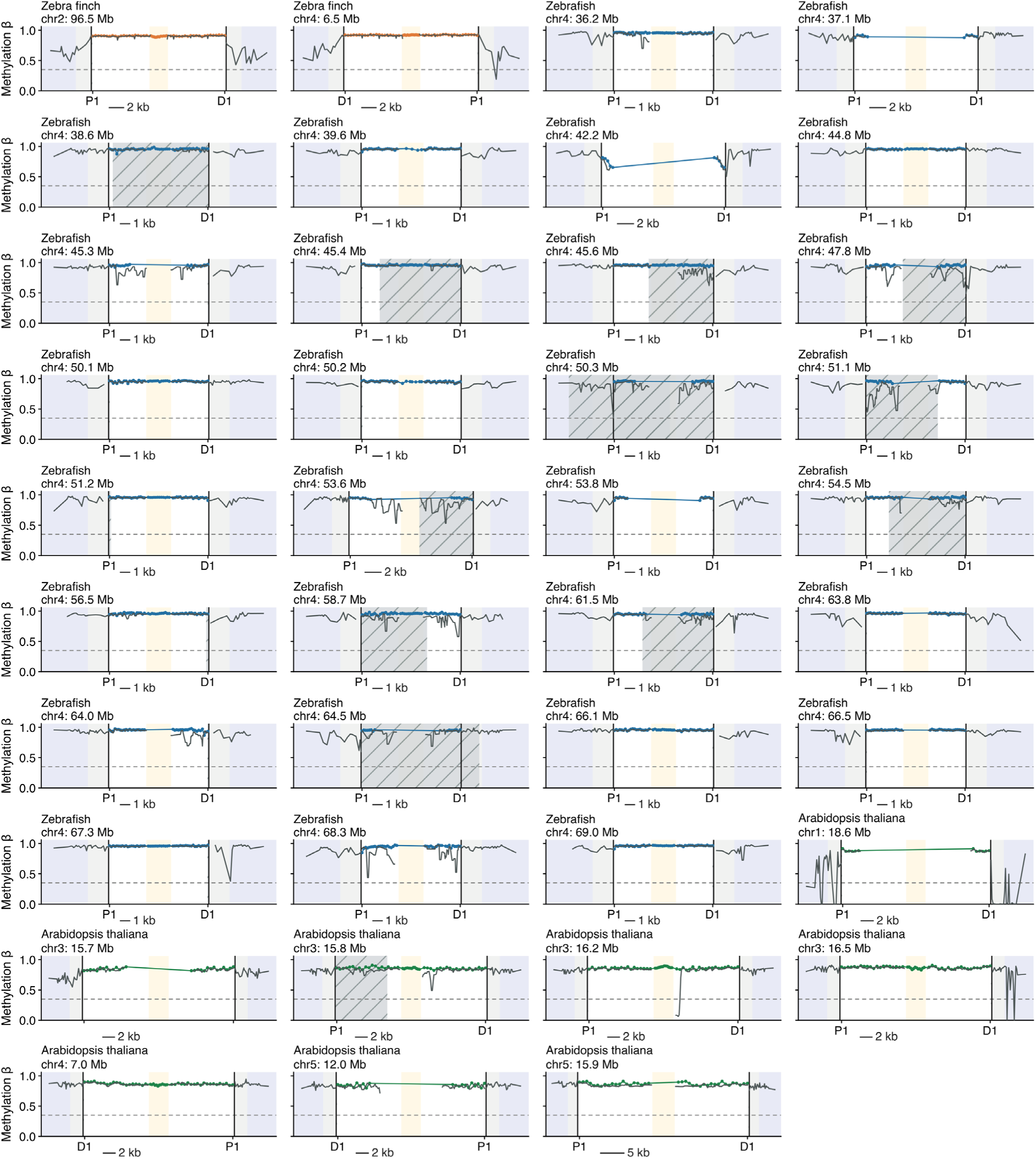
Methylation landscape of the 39 analyzed arrays not shown in Figure 7B. The same 39 arrays, drawn as Figure 7B. Each panel is one array, headed by the species and by the chromosome with the array position in Mb, carrying its own scale bar. Axes and the P1/D1 labels follow Figure 7. The proximal end falls on the left in 35 of the 39 panels and on the right in three. In the remaining panel (*A. thaliana* chr3, 15.7 Mb) the array lies inside the centromeric interval, so proximal and distal are not defined for it and P1/D1 are not shown.

**Supplementary Figure S13.**
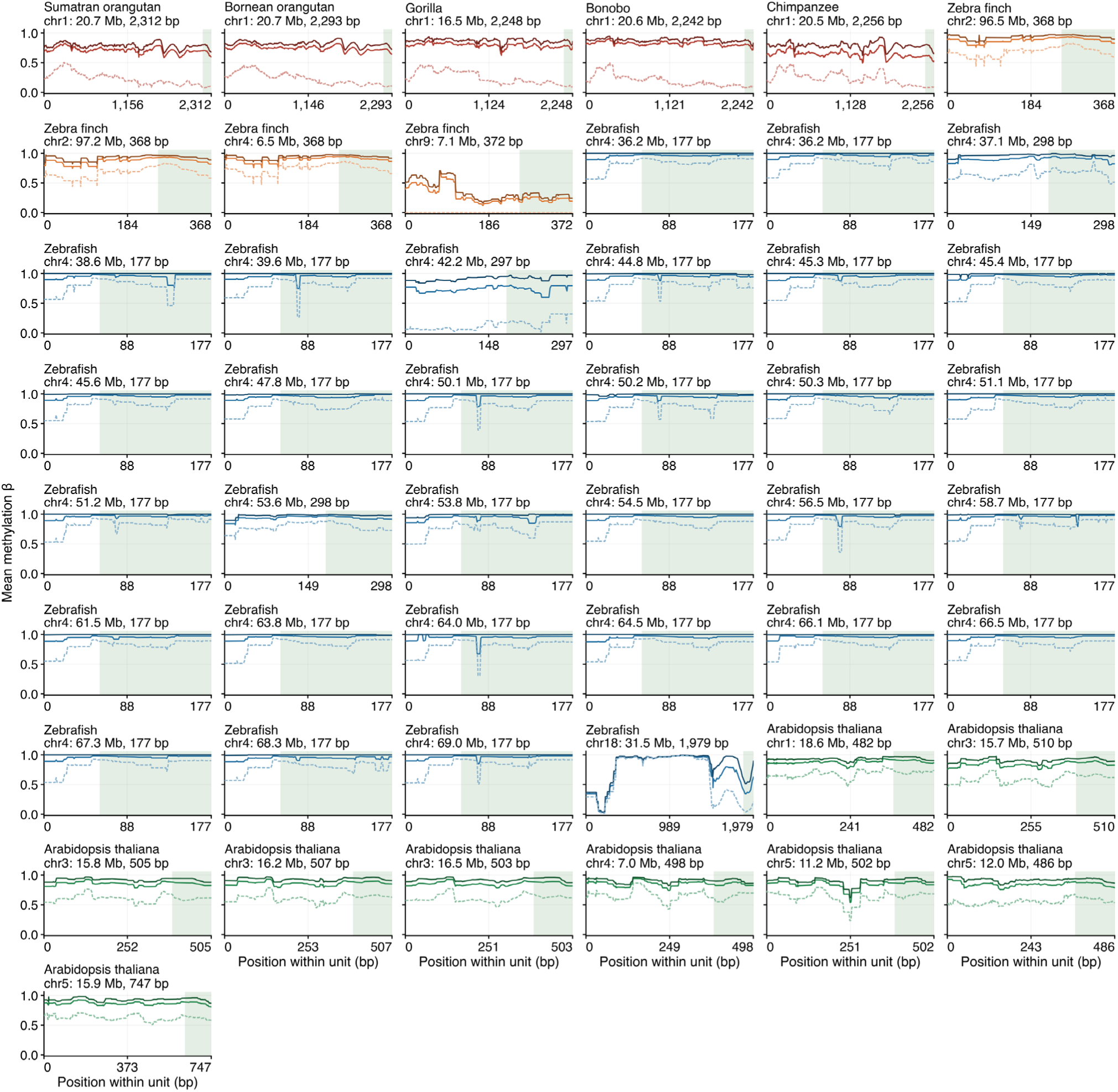
Methylation along the repeat unit of all 49 analyzed arrays. Each panel is one array, headed by the species and by the chromosome with the array position in Mb and the unit length in bp. Black, all complete units. Red, the high-methylation group. Blue, the low-methylation group. The green band marks the 5S rDNA gene, which lies at the end of the unit. The two groups are set per array and are defined in Materials and methods. Panels are ordered with the apes first.

